# Nasal tissue-resident memory CD4^+^ T cells persist after influenza A virus infection and provide heterosubtypic protection

**DOI:** 10.1101/2024.07.06.602325

**Authors:** Nimitha R. Mathew, Romain Gailleton, Lydia Scharf, Karin Schön, Josue Enriquez, Hannes Axelsson, Anneli Strömberg, Nils Lycke, Mats Bemark, Ka-Wei Tang, Davide Angeletti

**Affiliations:** Department of Microbiology and Immunology; Institute of Biomedicine; University of Gothenburg; Gothenburg; Sweden.; Department of Clinical Immunology and Transfusion Medicine, Sahlgrenska University Hospital, Gothenburg, Sweden; Department of Translational Medicine – Human Immunology, Lund University, Malmö, Sweden; Department of Infectious Diseases, Institute of Biomedicine, University of Gothenburg, Gothenburg, Sweden; Region Västra Götaland, Sahlgrenska University Hospital, Department of Clinical Microbiology, Gothenburg, Sweden; SciLifeLab, Institute of Biomedicine; University of Gothenburg, Gothenburg, Sweden

## Abstract

CD4 tissue-resident memory T cells (TRM) are crucial adaptive immune components involved in preventing influenza A virus (IAV) infection. Despite their importance, their physiological role in the upper respiratory tract, the first site of contact with IAV, remains unclear. Here, we find that, after IAV infection, antigen-specific CD4 TRM persist in the nasal tissue (NT) compartment after infection and provide protection upon heterosubtypic challenge. Single cell RNA sequencing analysis reveals that NT CD4 TRM are heterogeneous and transcriptionally distinct as compared to their lung counterparts. Mechanistically, we demonstrate that the CXCR6-CXCL16 axis promotes CD4 TRM residency in the NT. Furthermore, we show that the NT of mice and humans contains a high frequency of Th17 CD4 TRM that aid in local viral clearance and in reducing tissue damage. Collectively, our results support a robust physiological role for nasal tissue CD4 TRM in local protection during heterosubtypic IAV infection.

## INTRODUCTION

Influenza A virus (IAV) continues to pose a threat to society by causing yearly morbidity, deaths and providing a fertile ground for opportunistic bacterial infections (Paget et al., 2019). Seasonal vaccines elicit antibodies against the immunodominant surface glycoprotein hemagglutinin (HA) which is, however, prone to antigenic drift. Therefore, vaccines show limited effectiveness and are updated annually (Campbell et al., 2020). In addition, there is a hovering risk of pandemics caused by the recombination of animal and human viruses (The, 2024). Under such circumstances, antibody-mediated protection is usually inefficient. However, memory T cells recognize IAV internal proteins which are relatively conserved across different strains (Vijayanand et al., 2020), thus mediating heterosubtypic responses.

Among the memory T cells, tissue resident memory T cells (TRM) are gaining traction as a ‘ready to deploy’ memory arm that is formed at the site of infection and hence acts as a barricade before the pathogens reach the circulation or lymphoid organs where effector memory or central memory T cells can be activated (Marchesini Tovar et al., 2024). Indeed, during IAV infection, both CD4 and CD8 TRM in the lungs protect against heterologous viruses (Teijaro et al., 2011; Zheng et al., 2023). Compared to CD8 TRM, CD4 TRM are understudied, despite their multifunctionality. They can mediate viral clearance, act by direct cytotoxicity on virus-infected cells, guide the formation of IAV-specific lung CD8 TRM and orchestrate local B cell responses (Borkner et al., 2021; Brown et al., 2012; Son et al., 2021; Swarnalekha et al., 2021; Teijaro et al., 2011). Specifically, lung CD4 TRM are diverse and exhibit different functions depending on their antigenic specificity. For example, two subsets of IAV NP (nucleoprotein)- specific lung CD4 TRM were recently identified: T_RM_1 which are Th1 like, and T follicular helper like cells (T_RH_) which are involved in promoting local antibody response and delivering help to local CD8 T cells (Son et al., 2021; Swarnalekha et al., 2021). On the other hand, M2e (ectodomain of virus matrix protein)-specific lung CD4 TRM were shown to be mostly of the Th17 subtype and protect from tissue injury during IAV infection (Omokanye et al., 2022). However, our knowledge about CD4 TRM in other tissues relevant for respiratory viral infection is still limited.

The upper respiratory tract (URT) is the port of entry for any inhaled pathogen and the first site of contact with the immune cells of the body. The immune system within the mouse URT is a complex network of draining cervical lymph nodes (cLN), nasal-associated lymphoid tissues (NALT) and immune cells dispersed within the nasal chamber outside the NALT (referred as nasal tissues; NT) (Pizzolla et al., 2017a; Randall). The NALT is the site for the recall expansion of memory T cells and priming of naïve T cells, whereas cLN act as a priming site (Liu et al., 2024; Pizzolla et al., 2017b). Increasing evidence suggest that protective CD4 TRM are generated during bacterial and SARS-CoV-2 infection in the URT (Borkner et al., 2021; Diallo et al., 2023a; Kinnear et al., 2018; Lim et al., 2022; O’Hara et al., 2020; Wimmers et al., 2023; Xu et al., 2023; Yount et al., 2023). Nevertheless, the cues needed for URT CD4 TRM formation and tissue residency remain elusive. For IAV, some studies have demonstrated the protective role of URT CD8 TRM in reducing viral burden and transmission (Pizzolla et al., 2017a; Richard et al., 2020; Uddbäck et al., 2024). Surprisingly, despite the importance of CD4 TRM and the extensive spread of IAV, the role of IAV-specific CD4 TRM in URT has not been explored.

Here, we investigate antigen-specific CD4 TRM within the URT of IAV-infected mice and healthy human volunteers. We demonstrate that CD4 TRM form in NT of the URT and protect upon a heterologous IAV infection. Furthermore, we discover transcriptional differences between TRM populations in NT *vs* lungs and define factors required for the formation of antigen-specific CD4 TRM in the NT. Our data provide a comprehensive characterization of previously unexplored, IAV-specific CD4 TRM cells that reside in the URT.

## RESULTS

### IAV infection generates long-lived antigen specific CD4 TRM in the nasal tissue

To assess whether IAV-specific CD4 TRM arise in the URT, splenic CD4 T cells derived from Ovalbumin (OVA)-specific OT-II T cell receptor (TCR) transgenic mice that could be identified as CD45.1^+^ or tdTomato^+^ were transferred into congenic C57BL/6J (CD45.2^+^) mice. Subsequently, we infected the mice intranasally (i.n.) with a mouse-adapted IAV strain PR8 H1N1 that expresses the OVA_323-339_ peptide (PR8-OVA) and examined NALT and NT for OVA-specific CD4 TRM (OT-II CD4 TRM) on day 30 post infection. We used lungs as a positive control because IAV specific CD4 TRM are well known to form there (Teijaro et al., 2011). Throughout all our experiments, we administered fluorochrome-conjugated CD4 antibody intravenously (i.v.) 5 minutes before sacrifice to distinguish CD4 T cells in the tissue from those in the vasculature (Fig 1A). CD4 memory T cells were identified as being CD3^+^, CD4^+^, CD44^+^ and CD62L^-^. We differentiated CD4 TRM from their circulating counterparts as CD3^+^CD4^+^CD44^+^CD62L^-^ i.v.^-^ CD69^+^ (Fig S1A). CD69 is a well-known tissue retention molecule that prevents egress of T cells from the tissue(Shiow et al., 2006; Szabo et al., 2019b). OT-II CD4 TRM were present in the NT at a similar frequency as lungs. However, we found less OT-II CD4 TRM in the NALT (Fig 1B-C).

**Fig 1:**
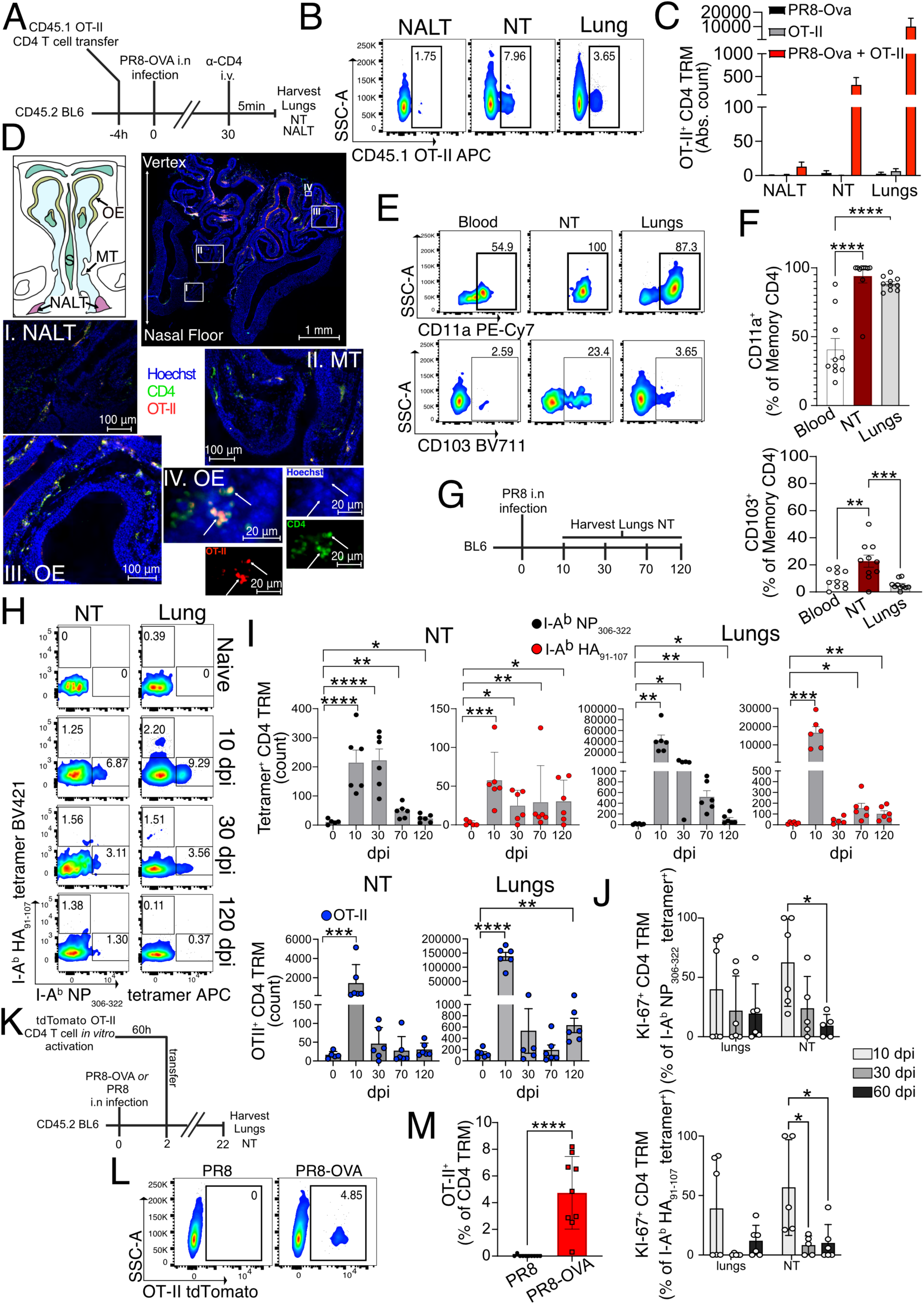
IAV-specific CD4 TRM are formed at the URT and is dependent on presence of cognate antigen. A) Schematic diagram showing the procedure of transfer of OVA specific (OT-II) CD4 T cells, infection of mice with PR8-OVA and subsequent harvesting of organs. B-C) CD45.1^+^ or tdtomato^+^ OT-II CD4 TRM (gated on live CD3^+^CD4iv^-^CD4 tissue^+^CD44^+^CD62L^-^CD69^+^ as shown in Fig S1A) in different organs of CD45.2^+^ recipients on day 30 following PR8-OVA infection. B) Representative flow cytometry plots indicating the percentages of CD45.1^+^ OT-II CD4 TRM. The data are representative of one experiment out of three independent experiments. C) Bar graph indicating the absolute number of CD45.1^+^ or tdtomato^+^ OT-II CD4 TRM. The experiment was repeated thrice and the results (mean ± s.e.m.) were pooled. D) Representative microscopic images of OT-II^+^ (red) CD4^+^ (green) T cells in different sections of the NT and NALT. The sections are derived from day 30 following PR8-OVA infection of mice that received OT-II CD4 T cells as described in fig. 1 A. Scale bars are indicated in each image. MT= Maxilloturbinate, OE= Olfactory epithelium, S= Nasal septum. E-F) Frequency of CD103^+^ and CD11a^+^ cells among OT-II CD4 TRM in NT and lungs (gated on live CD3^+^CD4iv^-^CD4 tissue^+^CD44^+^CD62L^-^CD69^+^), and among total CD4 effector memory T cells (CD3^+^CD4 iv^+^CD44^+^CD62L^-^) in blood of CD45.2^+^ recipients on day 30 following IAV infection. E) Representative flow cytometry plots indicating the percentages of CD103^+^ and CD11a^+^ cells. The data is representative of one experiment out of two independent experiments. F) Scatter plot indicating the percentages of CD103^+^ and CD11a^+^ cells. Each data point indicates an individual mouse. The experiment was repeated twice and the results (mean ± s.e.m.) were pooled. NS, not significant, \*\*\*\**P*<0.0001; \*\*\**P*<0.001; \*\**P*<0.01; \**P*<0.05 by one-way ANOVA, with Dunnet’s multiple comparison test. G-I) Kinetics of I-A^b^ NP_306-322_ tetramer, I-A^b^ HA_91-107_ tetramer-specific CD4 TRM and OT- II^+^ CD4 TRM in the NT and lungs on indicated days following PR8-OVA IAV infection. G) Schematic representation of the experimental setup. H) Representative flow cytometry plots indicating the percentages of I-A^b^ NP_306-322_ tetramer and I-A^b^ HA_91-107_ tetramer-specific CD4 TRM are shown. I) Scatter plot indicating the absolute numbers of I-A^b^ NP_306-322_ tetramer, I-A^b^ HA_91-107_ tetramer-specific CD4 TRM and OT-II^+^ CD4 TRM. Each data point indicates an individual mouse. The experiment was performed once and the results (mean ± s.e.m.) were plotted. NS, not significant, \*\*\*\**P*<0.0001; \*\*\**P*<0.001; \*\**P*<0.01; \**P*<0.05 by lognormal one-way ANOVA. J) Scatter plot showing the percentage of Ki-67^+^ cells among I-A^b^ NP_306-322_ tetramer^+^ and I-A^b^ HA_91-107_ tetramer^+^ CD4 TRM in lung and NT isolated on day 10, 30 and 60 post PR8 infection. The experiment was performed twice, and the results (mean ± s.e.m.) were pooled. NS, not significant, \*\*\*\**P*<0.0001; \*\*\**P*<0.001; \*\**P*<0.01; \**P*<0.05 by two-way ANOVA, with Tukey’s multiple comparison test. K-M) Frequency of OT-II CD4 TRM in the NT on day 22 following infection with PR8 or PR8-OVA. K) Schematic representation of the experimental setup. L) A representative flow cytometry plot indicating the frequency of OT-II CD4 TRM. M) Scatter plot showing the frequency of OT-II CD4 TRM. The experiment was performed twice and the results (mean ± s.e.m.) are pooled. NS, not significant, \*\*\*\**P*<0.0001; \*\*\**P*<0.001; \*\**P*<0.01; \**P*<0.05 by unpaired two-tailed t-test.

To localize OT-II CD4 T cells anatomically within the URT, we isolated, fixed and decalcified heads of PR8-OVA infected mice on day 30 post infection. OT-II CD4 T cells were found throughout the nasal chamber, including the olfactory epithelium (OE), respiratory epithelium lining the nasal septum (S), maxilloturbinates (MT) and NALT (Fig 1D). Given the localization of OT-II CD4 T cells in proximity to the epithelium of the NT, we examined the expression of CD103, an integrin involved in the adhesion of T cells to E-cadherin on epithelial cells. CD103 is often associated with CD8 TRM (Beura et al., 2019; Szabo et al., 2019b; Zens et al., 2017) but its association with CD4 TRM in the respiratory tract has not been elucidated, in mice. Indeed, CD103 was poorly expressed on lung CD4 TRM in comparison to NT CD4 TRM (5% *vs* 22%) (Fig 1E-F, Fig S1B). Apart from CD103, we analyzed NT CD4 TRM for the expression of CD11a, another marker associated with TRM residency at late days after infection (Szabo et al., 2019b; Teijaro et al., 2011). Nearly all OT-II CD4 TRM in the NT and lungs expressed CD11a (Fig 1E-F, Fig S1B).

Having demonstrated that NT CD4 TRM are established in a transfer model, we wanted to verify whether they would also form when precursor frequency is at physiological levels. Hence, we took advantage of MHC-II tetramers, presenting immunodominant peptides I-A^b^ HA_91-107_ and I-A^b^ NP_306-322_ (Besavilla et al., 2023; Miller et al., 2015). To follow the kinetics of antigen specific CD4 TRM, we infected mice with PR8 and harvested the lungs and NT at different time points (Fig 1G). We found that I-A^b^ HA_91-107_ tetramer^+^ and I-A^b^ NP_306-322_ tetramer^+^ CD4 TRM arose as early as 10 day post infection (dpi). The number of tetramer^+^ CD4 TRM declined from day 10 to day 30. Although, there was a steady decrease in tetramer-specific CD4 TRM in NT and lungs, we could still detect antigen-specific CD4 TRM in the NT on 120 dpi in most of the mice, suggesting this to be a relatively stable population (Fig. 1H-I, Fig S1C-E). We observed similar long-term kinetics for OT-II^+^ CD4 TRM in the OT-II transfer model (Fig 1I, Fig S1C-D).

IAV-specific CD4 TRM in the lungs are self-sustaining, without being replenished from the lymphoid reservoirs(Turner et al., 2014). To determine whether the same was true for NT CD4 TRM, we analyzed their expression of Ki-67, a marker for ongoing cell division (Miller et al., 2018). As expected, at the early stage after infection (10 dpi) approximately 60% of the IAV-specific NT CD4 TRM expressed Ki-67 (Fig 1J, Fig S1F), suggesting that some of those may still be in the effector phase. Conversely, at early (30 dpi) and late (60 dpi) memory time points only 10-15% of I-A^b^ HA_91-107_ tetramer^+^ and I-A^b^ NP_306-322_ tetramer^+^ NT CD4 TRM expressed Ki-67, indicating a low level of homeostatic proliferation (Fig 1J, Fig S1F). The importance of cognate antigen for the formation of TRM varies depending on tissue and T cell type; cognate recognition is a crucial factor for lung CD8 TRM formation(Takamura et al., 2016) but it is not required by NT CD8 TRM(Pizzolla et al., 2017a). Thus, we sought to assess the importance of cognate antigen maintenance for NT CD4 TRM formation. We generated effector tdTomato^+^ OT-II T cells *in vitro* and transferred them to C57BL/6 mice that were infected with either PR8 or PR8-OVA two days prior (Pizzolla et al., 2017a) (Fig 1K). In this model, even though effector OT-II cells can access the NT in both PR8 and PR8-OVA infected groups, we found that OT-II CD4 TRM were only retained in the NT of the PR8-OVA-infected mice (Fig 1L-M). This was also observed for lung CD4 TRM (Fig S1G-H). Our results demonstrate that cognate antigen, rather than only an inflammatory milieu, is required for the conversion of effector OT-II T cells to OT-II CD4 TRM in respiratory mucosal tissues, in contrast to NT CD8 TRM (Pizzolla et al., 2017a).

In summary, we found that IAV-specific CD4 TRM are generated and persist in the NT following infection.

### NT CD4 TRM are multifunctional and produce cytokines upon antigen stimulation

A rapid and efficient response to viral re-infection is a key feature of memory T cells, including CD4 TRM. Upon recognition of their specific antigen, memory T cells are reactivated and produce cytokines, some of which have a direct impact on virus-infected cells while others influence the recruitment of additional immune cells(Künzli and Masopust, 2023). To evaluate the IAV-specific response in NT CD4 TRM, we stimulated NT and lung cells including CD4 TRM from PR8-infected mice with an IAV-derived nucleoprotein (NP) peptide pool. Compared to unstimulated controls or naïve NT CD4 TRM, a higher proportion of peptide-stimulated NT CD4 TRM produced IFN-γ and IL-2, while no change was observed for TNF (Fig 2A-B and Fig S2A). Similarly, CD4 TRM sorted from either NT or lungs expressed higher amounts of intracellular IFN-γ when stimulated with splenocytes pulsed with NP peptide pool compared to SARS-CoV-2 nucleocapsid peptide pool, ruling out a role for ongoing inflammation in IFN-γ production (Fig 2C-D). NT and lung CD4 TRM displayed a similar immunodominance profile with regards to different IAV-derived peptides stimulation (Fig S2B). In general NT CD4 TRM harbored a lower frequency of cytokine producing cells than lung CD4 TRM (Fig 2A-D, Fig S2A). To exclude the possibility that this was due to that NT CD4 TRM had a more exhausted phenotype we measured PD-1 expression. CD4 TRM from both organs had a similar frequency of PD-1^+^ CD4 TRM indicating this was not the case (Fig 2E-F, Fig S2C) but suggesting a role for PD-L1-PD1 axis in CD4 TRM regulation, similarly to what was described for IAV-specific CD8 TRM(Wang et al., 2019). Surprisingly, even though NT CD4 TRM had overall lower frequency of cytokine producing cells, ICOS, a T cell co-stimulatory molecule (Moguche et al., 2015; Peng et al., 2022), was expressed on a higher proportion of NT CD4 TRM in comparison to lung CD4 TRM (Fig 2E-F, Fig S2C).

**Fig 2:**
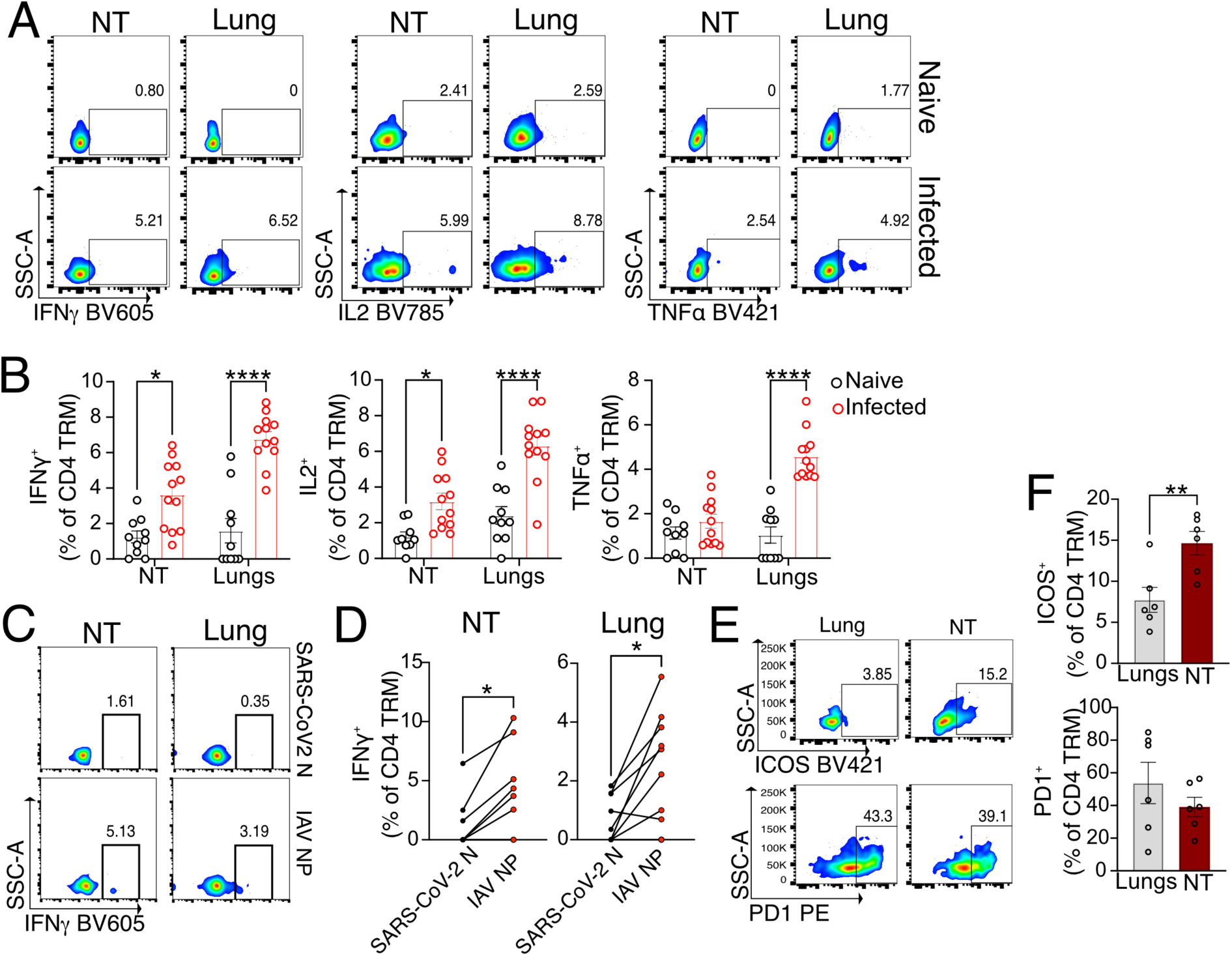
NT CD4 TRM express cytokines upon antigen stimulation. A-B) Expression of intracellular cytokines in CD4 TRM of NT and lungs upon restimulation with IAV NP peptide pool. The organs are isolated on day 30 following PR8 IAV infection of mice or from naïve mice. A) Flow cytometry plots indicating the percentage of CD4 TRM expressing IFN-γ, IL-2 and TNF. B) Scatter plot showing the percentage of CD4 TRM expressing IFN-γ, IL-2 and TNF. The experiment was repeated thrice and the results (mean ± s.e.m.) are pooled. NS, not significant, \*\*\*\**P*<0.0001; \*\*\**P*<0.001; \*\**P*<0.01; \**P*<0.05 by two-way ANOVA, with Tukey’s multiple comparison test. C-D) Frequency of IFNγ^+^ cells among all lung and NT CD4 TRM isolated on day 30 post PR8 infection and that are stimulated with peptide pool (IAV NP versus SARS-CoV-2 nucleocapsid) pulsed splenocytes. C) Representative flow cytometry plots showing the expression of IFNγ^+^ cells among all lung and NT CD4 TRM. D) Graph indicating the IFNγ^+^ cells among CD4 TRM stimulated by IAV NP versus SARS-CoV-2 nucleocapsid pulsed splenocytes connected by a line. Each line indicates an individual mouse. The experiment was performed three times and the results were pooled. NS, not significant, \*\*\*\**P*<0.0001; \*\*\**P*<0.001; \*\**P*<0.01; \**P*<0.05 by two-sided Wilcoxon matched-pairs signed-rank test. E-F) Expression of ICOS and PD-1 on CD4 TRM on d30 post PR8 IAV infection. E) Representative flow cytometry plots of CD4 TRM expressing ICOS and PD-1 in lungs and NT. F) Scatter plot showing the percentage of CD4 TRM expressing ICOS and PD-1. The experiment was repeated twice and the results (mean ± s.e.m.) are pooled. NS, not significant, \*\*\*\**P*<0.0001; \*\*\**P*<0.001; \*\**P*<0.01; \**P*<0.05 by unpaired two-tailed t-test.

### NT CD4 TRM provide protection upon heterologous IAV challenge

CD4 TRM promote viral clearance by recruiting and activating B cells, CD8 T cells and innate cells and by exerting direct cytotoxic effects on virus-infected cells(Künzli and Masopust, 2023). Given that NT CD4 TRM displayed an IAV-specific response (Fig 2A-D, Fig S2A), we set out to elucidate their function upon heterosubtypic challenge. To test this, we infected mice with PR8 and challenged them with heterologous X31 H3N2 virus on day 30 post primary infection. X31 has identical internal proteins to PR8 (including NP) but distinct surface glycoproteins (e.g. HA). First, we verified whether CD4 TRM respond to and expand locally at 6 days post rechallenge (Fig 3A). To rule out any contribution of circulating lymphocytes, we treated the mice with FTY720 during the experiment (Brinkmann et al., 2002). Indeed, frequencies of I-A^b^ NP_306-322_ tetramer^+^, but not I-A^b^ HA_91-107_ tetramer^+^, CD4 TRM sharply increased in both NT and lungs (Fig 3B-C and Fig S3A). The increase in I-A^b^ NP_306-322_ tetramer^+^ CD4 TRM was not due to expansion of naïve cells as it was not observed upon primary X31 infection (Fig S3B).

**Fig 3:**
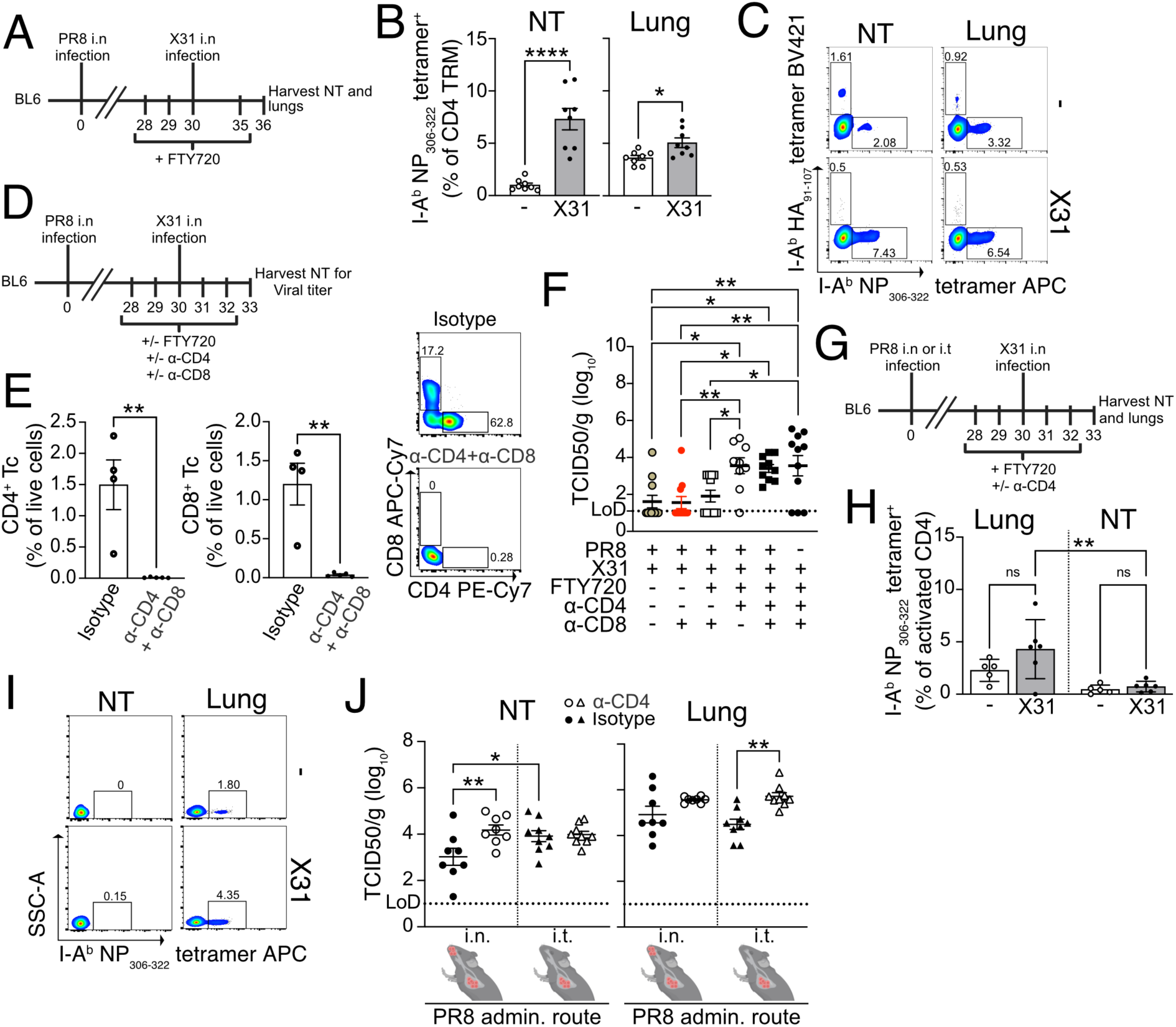
NT CD4 TRM provide protection against heterosubtypic challenge. A-C) I-A^b^ NP_306-322_ tetramer and I-A^b^ HA_91-107_ tetramer-specific CD4 TRM in the NT and lungs of mice that were infected with X31 intranasally or left uninfected on day 30 following PR8 infection. The mice were treated with FTY720 and the organs were analyzed 6 days post X31 infection. A) Schematic representation of the experimental setup. B) Scatter plot indicating the percentage of I-A^b^ NP_306-322_ tetramer ^+^ cells among CD4 TRM of lungs and NT. The experiment was done twice and the results (mean ± s.e.m.) were pooled. NS, not significant, \*\*\*\**P*<0.0001; \*\*\**P*<0.001; \*\**P*<0.01; \**P*<0.05 by unpaired two-tailed t-test. C) Representative flow cytometry plots indicating the percentages of I-A^b^ NP_306-322_ tetramer and I-A^b^ HA_91-107_ tetramer-specific CD4 TRM are shown. D) Schematic representation of primary and secondary infection of mice, their treatment with anti-CD4 antibody, anti-CD8 antibody or/and FTY720 on the indicated days and collection of NT. E) Left panel: Scatter plot showing the frequency of CD4 Tc and CD8 Tc among all live cells in the NT of mice treated with isotype control antibody or anti-CD4 and anti-CD8 antibody. The experiment was performed once. The result is shown as (mean ± s.e.m.) NS, not significant, \*\*\*\**P*<0.0001; \*\*\**P*<0.001; \*\**P*<0.01; \**P*<0.05 by unpaired two-tailed t-test. Right panel: A representative flow cytometry plot for the frequency of CD4 and CD8 Tc. F) Scatter plot indicating viral titers (TCID_50_/g) in NT of mice on day 3 following secondary infection with X31 IAV. The experiment was repeated thrice and the results (mean ± s.e.m.) are pooled. NS, not significant, \*\*\*\**P*<0.0001; \*\*\**P*<0.001; \*\**P*<0.01; \**P*<0.05 by one-way ANOVA, with Dunnet multiple comparison test. The outliers were identified and removed using Grubbs method (alpha=0.05). One outlier each removed from all groups except for groups: PR8 + anti-CD8 + FTY720 and PR8 + anti-CD4. G-J) Viral titer and frequency of I-A^b^ NP_306-322+_ CD4 TRM in the NT and lungs of mice who have undergone intratracheal or intranasal PR8 infection followed by intranasal infection with X31 on day 30 post primary infection. The mice were treated with FTY720+/-anti-CD4 antibody or left untreated. G) Schematic representation of the experimental set up. H) Scatter plot indicating the frequency of I-A^b^ NP_306-322_ tetramer^+^CD4iv^-^CD44^+^CD62L^-^ cells in the NT and lungs of mice on day 3 post-secondary X31 infection +FTY720 treatment vs mice without secondary X31 infection and not treated with FTY720 as shown in fig 3G. The experiment was repeated thrice and the results (mean ± s.e.m.) are pooled. NS, not significant, \*\*\*\**P*<0.0001; \*\*\**P*<0.001; \*\**P*<0.01; \**P*<0.05 by two-way ANOVA, with Tukey’s multiple comparison test. I) Representative flow cytometry plots indicating percentages of I-A^b^ NP_306-322_ tetramer^+^CD4iv^-^CD44^+^CD62L^-^ cells in NT and lung of mice with and without secondary X31 infection (related to 3H). J) Scatter plot showing viral titers (TCID_50_/g) in NT and lungs of mice on day 3 following secondary infection with X31 IAV and who were infected with PR8 intranasally or intratracheally 30 days prior to X31 infection. The mice were treated with FTY720 ip as indicated in fig 3G. The experiment was repeated twice and the results (mean ± s.e.m.) are pooled. NS, not significant, \*\*\*\**P*<0.0001; \*\*\**P*<0.001; \*\**P*<0.01; \**P*<0.05 by one-way ANOVA, with Dunnet multiple comparison test.

To define a possible role in protection, we performed the same experiment as above but instead measured the viral titer in NT on day 3 post X31 infection. In addition to FTY720 treatment, we depleted either CD4 T cells, CD8 T cells or both (Fig 3D). To effectively clear TRM from respiratory mucosal tissues we injected mice with a high dose of depleting mAbs (Goplen et al., 2020; Son et al., 2021) (Fig 3E and Fig S3C-D). In the absence of any intervention, the secondary infection was cleared from the NT by cross-reactive T cells (Fig 3F), as PR8-antibodies did not neutralize X31 (Fig S3E). Surprisingly, depletion of CD8 T cells did not affect the viral clearance in the NT (with or without FTY720 administration) bolstering the role of CD4 TRM in local protection. Conversely, CD4 T cell depletion severely impaired viral clearance, with viral titers similar to those of the mice that did not undergo a primary infection (Fig 3F). To further exclude any possible contribution of lung TRM, despite FTY720 administration, we used an intratracheal infection (i.t.) model for the primary IAV inoculation (Fig 3G). Upon i.t. infection, IAV-specific CD4 TRM formed only in the lungs but not in the NT (Fig S3F). Accordingly, heterologous i.n. X31 challenge, combined with FTY720, administration resulted in the expansion of I-A^b^ NP_306-322_ tetramer^+^ CD4 T cells only in lungs but not in NT (Fig 3H-I). When compared to local expansion after i.n. prime (Fig 3B), the results strongly suggest that TRM are the major players involved in the local response. In addition, we repeated the CD4 depletion experiment (Fig 3G) combined with i.n. or i.t. prime and X31 challenge (Fig 3J): as expected, CD4 depletion affected viral titers in lungs. Conversely, i.n. primed mice (both NT and lung CD4 TRM) had lower viral titers in NT when compared to i.t. primed mice (only lung TRM) and CD4 treatment impaired protection in NT only when local CD4 TRM were present (Fig 3J). Our overall results (Fig 3F-J) strongly support that NT CD4 TRM contribute to protection independently of lung CD4 TRM.

To verify whether NT CD4 TRM were also induced by vaccination, we established an i.n. vaccination model using CTA1-3M2e-DD as an immunogen. CTA1-3M2e-DD contains 3 tandem repeats of M2e linked to CTA1-DD adjuvant; M2e is highly conserved across IAVs and immunization with CTA1-3M2e-DD has been shown to induce M2e specific lung resident CD4 T cells that protect against lethal IAV challenge in mice(Eliasson et al., 2008; Eliasson et al., 2018). For these experiments, we used BALB/c rather than C57BL/6 as CTA1-3M2e-DD induces poorer M2e specific T cell responses in C57BL/6(Eliasson et al., 2018). To maximize the formation of CD4 TRM in NT and not lung, we modified the immunization protocol and reduced the volume to 5μl. We vaccinated mice with three i.n. doses of CTA1-3M2e-DD or CTA1-DD and evaluated the M2e specific CD4 Tc response on day 30 (day 10 after last vaccination) (Fig S3G). Like the response observed after infection, we found that I-A^d^ M2e_2-17_ tetramer^+^ CD4 Tc robustly formed in the NT (Fig S3H-I). Furthermore, I-A^d^ M2e_2-17_ tetramer^+^ CD4 TRM formed and persisted only in the NT and cLN on day 50 (day 30 after last vaccination) with almost none in the lungs and few in mediastinal lymph nodes (MLN) (Fig S3J-M). To ascribe a role to M2e-specific CD4 TRM, we immunized mice with CTA1-DD or CTA1-3M2e-DD and challenged them at day 50 with high dose of PR8 in low inoculum volume, to restrict initial viral replication to the URT (Gailleton et al., 2025) (Fig S3N). This immunization strategy mainly generates M2e-specific CD4 TRM with no CD8 TRM (Eliasson et al., 2018). After challenge, 45% of the mice that were immunized with the control succumbed to the disease, however, immunization with CTA1-3M2e-DD protected against mortality and reduced weight loss (Fig S3O-P). Surprisingly, we could not detect differences in viral titers in neither lungs nor NT between the groups suggesting that M2e-specific CD4 TRM rely on other mechanisms for protection than viral control (Fig S3Q), as previously observed (Omokanye et al., 2022).

Taken together, our findings revealed that NT CD4 TRM are important for protection during heterosubtypic infection.

### CD4 TRM in the lungs and NT are heterogeneous, but Th17 cells are prevalent in the NT

Having identified the dynamics and role of NT CD4 TRM, we wanted to investigate the transcriptional differences between total and antigen-specific CD4 TRM in NT *vs* lungs. Therefore, we performed scRNA-seq together with scTCR-seq at 30 dpi. Mice were injected with CD4 antibody i.v. 5 minutes before sacrifice and total CD4 TRM, I-A^b^ HA_91-107_ tetramer^+^ and I-A^b^ NP_306-322_ tetramer^+^ CD4 TRM were sorted from NT and lungs of mice (Fig. 4A). Furthermore, we also sequenced a small proportion of all live cells, which were CD4^-^. To maximize the number of cells and samples that could be loaded simultaneously, different samples were labeled with hashtag antibodies. Data from all mice and organs were integrated, processed, and visualized on a two-dimensional UMAP (Fig 4B). T cell subpopulations were assigned based on well-known markers and gene signatures (Fig 4C, Fig S4A-B and Table S1). The major T cell subpopulation was, as expected given the intracellular viral infection, Th1 cells (Fig S4C) defined by a well-established signature(Andreatta et al., 2022), including *Ifng* and *Cxcr3*. Cytotoxic CD4 T cells were present in both organs and defined by *Nkg7*, *Fasl*, *Ccl4*, *Ccl5* and a CD4 cytotoxic signature(Hashimoto et al., 2019). We were unable to properly define a third major cluster, C.3; this cluster harbored several mitochondrial genes, even though all mitochondrial genes were regressed out and did not contribute to the clustering. Therefore, we believe these cells to be either senescent/dying cells or highly proliferative cells. Tfh cells were generally more prevalent within lungs (Fig S4C), consistent with a possible role in supporting ectopic GC in iBALTs. Based on their transcriptional profile they were separated into two clusters, named Tfh.1 and Tfh.2. Tfh.1 were Tfh cells with strong expression of *Tnfsf8*, *Bcl6*, and a signature previously associated with Tfh (Andreatta et al., 2022). Nevertheless, Tfh.2 had higher expression of other Tfh-associated genes such as *Tox*, *Slamf6* and *Pdcd1*. C.5 cells expressed several transcription factors, *Cd40lg* but also had high expression of genes involved in regulation of NF-kB signaling (*Nfkbia*, *Nfkbid*, *Nfkbiz*). This cluster has been previously reported and suggested to include cells which are poised to rapidly respond to secondary infections by producing cytokines (Kurd et al., 2020; Woodring et al., 2022). Quiescent memory cells were slightly more prevalent within NT and characterized by the expression of several long non-coding RNAs (including *Malat1*)(Dey et al., 2023; Plasek and Valadkhan, 2021) and low ribosomal gene expression. A small proportion of our CD4 TRM was naïve-like T cells expressing *Ccr7*, *Sell* and *Klf2 (Wang et al., 2022)*. Furthermore, we identified Tregs by *Foxp3* expression and Treg-signature(Yang et al., 2021). Early activated cells were defined based on the expression of genes linked with migration within the tissue, such as *Gpr183(Li et al., 2016)*, and signaling receptors such as *Ms4a4b*(Yan et al., 2012) suggesting an overall movement within the tissue but also modulation of T cell activation. NKT cells and IFN responsive cells were clearly identified by their distinctive signatures: the former expressed *Klrd1*, *Klrk1*, *Klrb1c*(Cohen et al., 2013; Hu et al., 2022) and other cytotoxic genes (*Nkg7*, *Fasl*)(Ng et al., 2024; Tuomela et al., 2022) while the latter was enriched for Interferon stimulated genes (*Isg15*, *Ifit3*, *Ifit1*), a signature similar to one previously identified in human CD4(Szabo et al., 2019a). Strikingly, Th17 cells, expressing *Rorgc*, *Il17a*, *Tmem176a* and *Ccr6* were much more abundant within NT(Szabo et al., 2019a) (Fig S4C and Table S1). We were also able to identify several endothelial and epithelial airway cells clusters, from the live cell fraction devoid of CD4 T cells.

**Fig 4:**
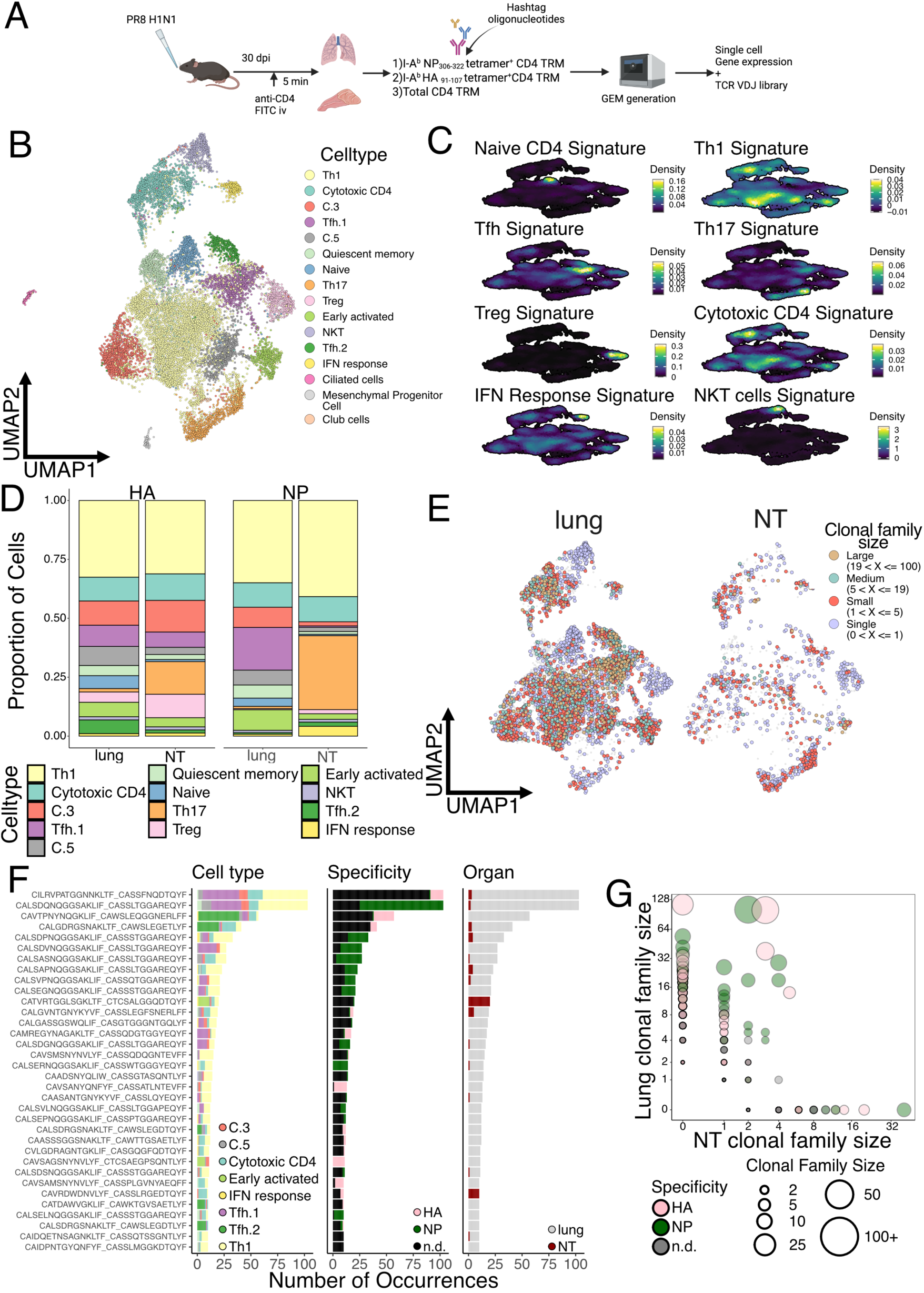
Single cell RNA and TCR sequencing of CD4 TRM reveals differential cluster distribution but clonal sharing between organs. A) Schematic diagram showing the preparation of antigen-specific cells from PR8 IAV infected mice for generation of GEM using 10X Chromium controller. B) UMAP plot of unsupervised clustering for 15301 CD4 TRM isolated from lungs and NT of naïve and PR8 IAV infected mice. C) Density plot showing the expression of selected gene signatures, projected on the same UMAP as in B. Naïve CD4 T signature(Wang et al., 2022), Th1 signature(Andreatta et al., 2022), Tfh signature(Andreatta et al., 2022), Th17 signature(Szabo et al., 2019a), Treg signature(Yang et al., 2021), Cytotoxic CD4 signature(Hashimoto et al., 2019), IFN response signature(Szabo et al., 2019a), NKT cells signature(Cohen et al., 2013; Hu et al., 2022). D) Bar graph showing proportion of each UMAP cluster divided by organ and antigen specificity. Only antigen specific CD4 TRM from infected mice were included in the analysis. E) UMAP plot split by tissue (lung or NT) and colored according to clone size for CD4 TRM in IAV infected mice. Clones were binned into single clones (1 member; light blue), small (2 to 5 members; red), medium (6 to 19 members; green), large (over 20 members; gold). Light grey dots represent cells where TCR clonotype could not be assigned. F) Bar plot characterizing the top 34 most expanded clones. The y axis reports CDR3 sequences of TCRαβ while the bar graphs show the number of cells divided according to cluster distribution, I-A^b^ NP_306-322_ tetramer^+^ or I-A^b^ HA_91-107_ tetramer^+^ and organ specificity. n.d. = not determined. G) Graph showing clonal relationship between lungs and NT from PR8 IAV infected mice. Each circle indicates one clonal family and size of the circle defines the size of the clonal family. Color of the circle denotes the antigenic specificity against I-A^b^ NP_306-322_ tetramer and against I-A^b^ HA_91-107_ tetramer. n.d. = not determined.

Overall, our scRNASeq analysis highlighted the diversity and variety of CD4 TRM subpopulations in both lungs and NT. While Th1 and cytotoxic CD4 T cells made up almost 50% of the total TRM in both lungs and NT, consistent with an intracellular viral infection, their relative proportions were similar between organs (Fig. S4C). Interestingly, IAV infection did not appear to majorly reshape the CD4 TRM landscape in terms of the abundance of cell type (Fig S4D). Therefore, to confirm our results and limit our analysis to influenza-specific cells, we subsetted the CD4 cells to only include the tetramer^+^ cells. Indeed, this did not change the proportion of CD4 TRM, confirming an increased abundance of both HA^+^ and NP^+^ Th17 cells within the NT, while these cells were almost absent in lungs (Fig 4D and Fig S4E). By visualizing the distribution of HA^+^ and NP^+^ cells as density plots we established that antigen-specific cells differentiated into almost all CD4 TRM subsets, except for Th17 in the lungs. However, the density distribution in NT was dramatically different between the two proteins, with NP^+^ cells being mostly Th17 and Th1 while HA^+^ cells also included many cytotoxic and Tfh CD4 TRM cells (Fig 4D and Fig S4E).

### CD4 TRM within the NT exhibit a distinct transcriptional profile, compared to lungs

The power of the single cell approach is that it allows transcriptional differences to be compared between the same CD4 TRM subtypes, depending on the organ. Therefore, we performed differential gene expression (DEG) analysis for each cell subpopulation and visualized the results with both heatmap and volcano plots (Fig S4F-G and Table S1). As expected, the majority of CD4 TRM subpopulations exhibited similar gene expression between organs, however Th1, Cytotoxic CD4, Tfh.1 and Th17 clusters were the ones with most organ-specific differences. For Th1, lung cells had higher expression of *Ccr2*, a chemokine receptor previously linked with pulmonary homing(Brownlie et al., 2022; Hoft et al., 2019), *Ctla4* but also *Ifng* and *Nfkbia,* suggesting a more active and productive phenotype, compared to that of their NT counterpart, in line with what we observed above (Fig 2A-B). On the other hand, NT Th1 were enriched for other essential transcription factors such as *Crem* and *Taf7* which are involved in cytokine production(Devaiah et al., 2010; Rauen et al., 2013) and *Xcl1*, which promotes Treg differentiation(Nguyen et al., 2008), suggesting an overall more immune-suppressive state. Cytotoxic CD4 cells had similar pattern with *Ccr2* and genes associated with a more active phenotype expressed in the lungs. Interestingly, NT CD4 TRM upregulated several chemokine receptors, including *Cxcr6* and *Ccr8,* and many other factors previously associated with T cells at mucosal sites (such as *Spp1* and *Penk*)(Kourepini et al., 2014; Mas-Orea et al., 2023). Remarkably, *Cxcr6* was also significantly upregulated in NT Tfh.1 and Th17 clusters. In addition, Tfh.1 NT cells also shared *Tff1* and *Xcl1* upregulation with other clusters, while NT Th17 exhibited increased *Crem* and *Nr4a2* expression.

In summary, several CD4 TRM subpopulations displayed differential gene expression between lungs and NT, with the lung cells showing a more activated phenotype and higher *Ccr2* expression, consistent with their localization. Conversely NT CD4 TRM expressed genes commonly associated with T cells at mucosal sites and with an immune-dampening phenotype, with several clusters also expressing the chemokine receptor *Cxcr6*.

### Expanded clonotypes have multiple fates and populate both lungs and NT

Having identified differences between CD4 TRM residing in different organs we wanted to investigate whether T cell clones were compartmentalized or shared between them. A caveat of our experimental procedure is that we had to pool several mice to obtain sufficient numbers of antigen-specific cells and, therefore, we were not able to define clonality at the single mouse level. However, a previous study suggested that CD4^+^ T cell responses are largely private even for a single viral-peptide specificity(Andreatta et al., 2022). Consistent with the notion of a largely private response we did not detect any exaggeratedly expanded T cell clones, with only 5 families with more than 30 members. Expanded clones were distributed across the majority of CD4 TRM subpopulations within the lungs, except for the naïve, Treg, NKT and Th17 which were composed mainly of single clones (Fig 4E). Conversely, NT exhibited a higher clonal expansion within Th17. When focusing only on antigen-specific clones, we detected a very similar pattern of clonal expansion (Fig S5A). To visualize the specificity, organ and subpopulation distributions, we plotted the top 34 expanded clones (Fig 4F). This confirmed the pattern of expansion and that most of expanded clones were shared among several subpopulations but also between organs. Validating our tetramer sorting approach, clones were either HA or NP specific, however, in many cases, we did not sort all clones within the same family, as also expected. Substantiating the private nature of TCR responses was the fact that many of the NP expanded clones had similar, but not identical, CDR3 sequences (Fig 4F). This suggests that certain TCRs had higher affinity for NP peptide but stochastic variations, not altering binding, were present in different mice. Indeed, analysis of paired TCRa-TCRb usage demonstrated a strong bias in NP responses for TRAV6-7_TRBV3 for both lungs and NT (Fig S5B). By visualizing with a bubble plot, in which the size of the bubble is proportional to the clone size, we revealed that many of the more expanded, antigen-specific clones in the lungs were also shared with the NT (Fig 4G). This points toward a common origin within the same lymph nodes and subsequent migration to tissues of residency. Finally, to determine whether certain clonotypes were more prone to direct CD4 TRM toward a certain fate we utilized the “clonotype bias” measure introduced by Andreatta et al.(Andreatta et al., 2022) and implemented within the scRepertoire package(Borcherding et al., 2020). The analysis revealed that the majority of clonotypes were largely unbiased with few exceptions (Fig S5C). Interestingly, clonotype bias strongly correlated with being organ-specific (Fig S5C-D). For instance, the third largest clone was present only in the lungs and over 60% of its members belonged to the Tfh.2 cluster (Fig S5C-D). Likewise, the only large clone found exclusively in NT had just shy of 50% of its members belonging to the early activated CD4 TRM group (Fig S5D). However, the most expanded clones were generally not biased, despite having a strong prevalence of Th1 and Tfh.1 cells (Fig S5E). The tendency of the few biased clones to be organ specific could also suggest local proliferation and expansion.

Combined TCR analyses unveiled how most CD4 TRM clonotypes can differentiate into multiple fates and then reside in both lungs and NT. However, there were few clones that were tissue-specific, and they tended to be biased toward one CD4 T cell subtype.

### CXCR6-CXCL16 axis promotes CD4 TRM recruitment to the NT

Given that *Cxcr6* was highly expressed in several NT clusters (Fig S4F-G) we further explored the mechanistic involvement of this chemokine receptor in the migration of effector CD4 T cells to the NT. First, we confirmed *Cxcr6* to be the most interesting candidate as it was more highly expressed in NT CD4 TRM than in lung CD4 TRM (Fig 5A-C). In particular, the clusters of cytotoxic CD4 T cells, Tfh.1, Th17, Treg, early activated and Tfh.2 displayed higher expression of *Cxcr6* in NT (Fig 5B). CXCR6 is known to be predominantly expressed in CD8 TRM rather than circulating CD8 T cells (Mabrouk et al., 2022). Similarly, we found high CXCR6 expression in CD4 TRM and low expression in CD4 effector memory T cells (TEM i.v.^+^) of IAV-infected mice (Fig 5D). While CXCR6 expression was greater in infected NT than lungs, its expression was similar to that of NT CD4 TRM in naïve mice (Fig 5D). This may seem surprising, however, by definition, TRM are not naïve cells and therefore their presence in NT has been triggered by a previous insult. Furthermore, we also found higher CXCR6 expression among antigen specific OT-II TRM in NT *vs* lungs (Fig 5E-F).

**Fig 5:**
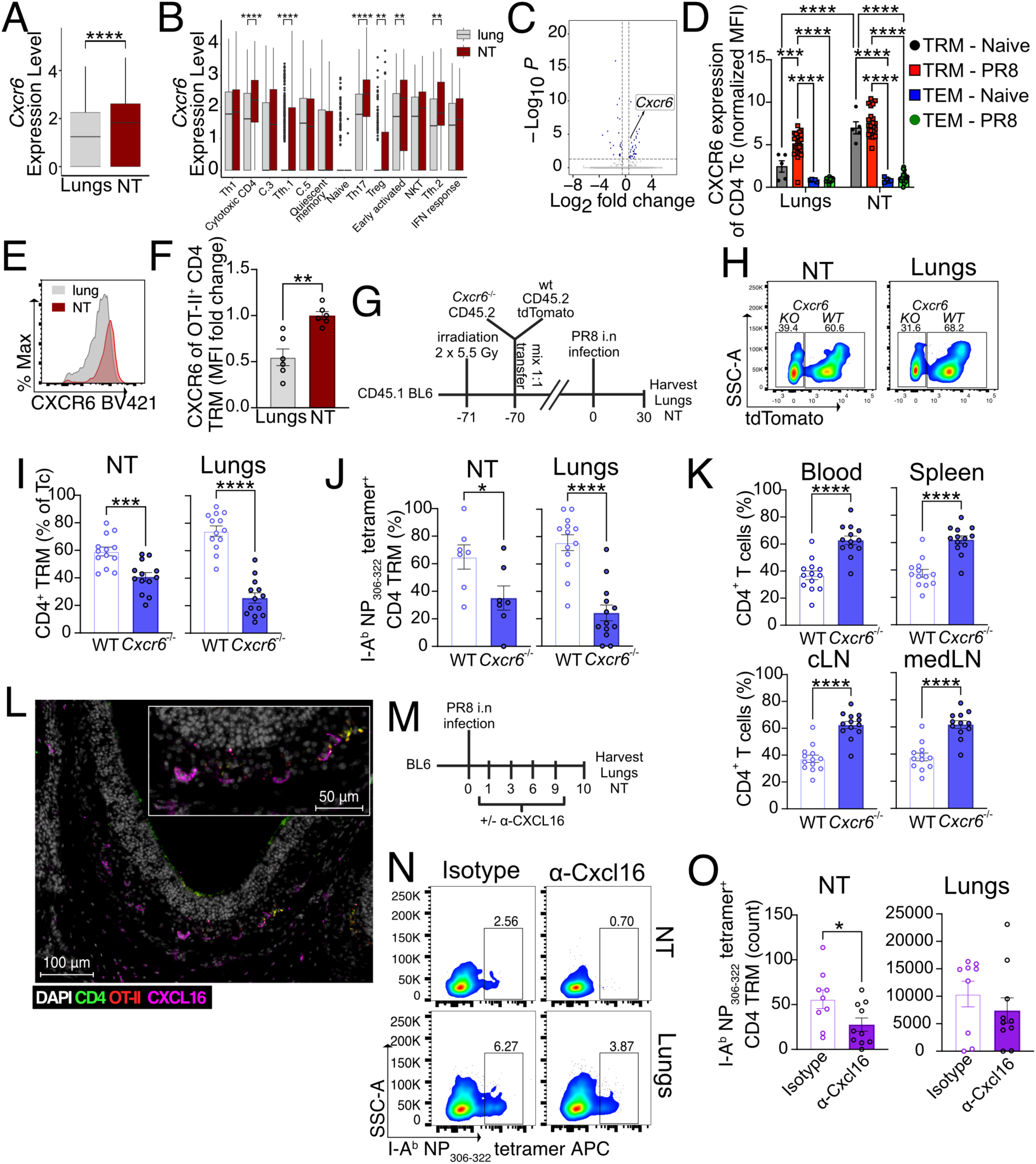
CXCR6-CXCL16 axis promotes NT CD4 TRM establishment. A) Box plot showing the expression of *Cxcr6* mRNA among CD4 TRM of the NT and lungs. Data are presented as median and interquartile range. NS, not significant, \*\*\*\**P*<0.0001; \*\*\**P*<0.001; \*\**P*<0.01; \**P*<0.05 by unpaired two-tailed t-test. B) Box plot showing the expression of *Cxcr6* mRNA among different cell clusters of lungs and NT. Data are presented as median and interquartile range. NS, not significant, \*\*\*\**P*<0.0001; \*\*\**P*<0.001; \*\**P*<0.01; \**P*<0.05 by Wilcoxon rank sum test with Benjamini-Hochberg correction C) Volcano plot for differential expressed genes in the NT in comparison to the lungs of the Th17 cluster. The dotted lines indicate fold change and adjusted p value cutoffs. D) Bar plot for the expression of CXCR6 (each Median Fluorescence Intensity (MFI) normalized to mean MFI from CD4 TEM iv^+^ of NT) in CD4 TRM and iv^+^ CD4 TEM from the lungs and NT of naïve mice and PR8 IAV infected mice (30 dpi) as indicated. The experiment was repeated thrice and the results (mean ± s.e.m.) are pooled. NS, not significant, \*\*\*\**P*<0.0001; \*\*\**P*<0.001; \*\**P*<0.01; \**P*<0.05 by two-way ANOVA, with Tukey’s multiple comparison test. E-F) Expression of CXCR6 on OT-II CD4 TRM of NT and lung on d30 following PR8-OVA infection as indicated. E) Representative histograms for CXCR6 expression from OT-II CD4 TRM of NT and lung are shown. F) Scatter plot showing the expression of CXCR6 (each Median Fluorescence Intensity (MFI) normalized to mean MFI of NT OT-II CD4 TRM). The experiment was repeated thrice and the results (mean ± s.e.m.) are pooled. NS, not significant, \*\*\*\**P*<0.0001; \*\*\**P*<0.001; \*\**P*<0.01; \**P*<0.05 by unpaired two-tailed t-test. G-K) Frequency of different cell populations within different organs derived from WT-*Cxcr6*^-^ _/-_ BM chimeric mice on day 30 post infection with PR8 IAV as indicated. G) Schematic representation of the experimental setup. H) Representative flow cytometry plots showing the percentage of WT and *Cxcr6*^-/-^ CD4 TRM in NT and lungs. I) Scatter plot for the percentage of WT and *Cxcr6*^-/-^ CD4 TRM in NT and lungs as indicated. J) Scatter plot for the percentage of I-A^b^ NP_306-322_ tetramer - specific WT and *Cxcr6*^-/-^ CD4 TRM in NT and lungs as indicated. Samples that had less than 7 OT-II CD4 TRM were excluded from the analysis. The experiment was repeated thrice and the results (mean ± s.e.m.) are pooled. NS, not significant, \*\*\*\**P*<0.0001; \*\*\**P*<0.001; \*\**P*<0.01; \**P*<0.05 by unpaired two-tailed t-test. K) Scatter plot showing the percentage of *Cxcr6*^-^ _/-_ and WT CD4 T cells in different organs of the BM chimera on day 30 post infection with PR8. The experiment was repeated thrice and the results (mean ± s.e.m.) were pooled. NS, not significant, \*\*\*\**P*<0.0001; \*\*\**P*<0.001; \*\**P*<0.01; \**P*<0.05 by unpaired two-tailed t-test. L) A representative microscopic image of CXCL16 expression (magenta) and OT-II CD4 T cells (red and green) in the olfactory epithelium of the murine NT. NT is isolated on day 30 post PR8-OVA infection from mice that received OT-II CD4 T cells. M-O) Antigen-specific CD4 TRM in the lungs and NT of mice treated with isotype control or anti-CXCL16 antibody on day 10 post PR8 IAV infection. M) Schematic representation of the experimental setup. N) Representative flow cytometry plots indicating the percentage of I-A^b^ NP_306-322_ tetramer - specific CD4 TRM are shown. O) Scatter plot indicating the absolute number of I-A^b^ NP_306-322_ tetramer - specific CD4 TRM. The experiment was performed twice and the results (mean ± s.e.m.) are pooled. NS, not significant, \*\*\*\**P*<0.0001; \*\*\**P*<0.001; \*\**P*<0.01; \**P*<0.05 by unpaired two-tailed t-test.

Within the lungs, CXCR6 is essential for the positioning of CD8 TRM in the airways but not in the lung interstitium after IAV infection(Wein et al., 2019). To determine whether CXCR6 is required for the establishment of CD4 TRM in the NT, we generated WT-*Cxcr6*^-/-^ bone marrow (BM) chimeras. To do this, we lethally irradiated congenic C57BL/6 (CD45.1^+^) (recipient) to deplete the host bone marrow and reconstituted them with equal numbers of tdTomato^+^ WT BM (CD45.2^+^) and tdTomato^-^ *Cxcr6*^-/-^ BM (CD45.2^+^) (Fig 5G). Blood chimerism was verified on day 67 post BM transfer, with almost 90% of the leukocytes being donor-derived. Interestingly, we detected a higher frequency of *Cxcr6*^-/-^ leukocytes in the blood (Fig S6A). On day 70 post BM transfer, we infected the mice and analyzed organs at day 30 post infection. CXCR6 expression helped in lodging total and I-A^b^ NP_306-322_ tetramer^+^ CD4 TRM in both NT and lungs, as we observed a reduced frequency of *Cxcr6*^-/-^ CD4 TRM in comparison to WT CD4 TRM in both compartments (Fig 5H-J). Surprisingly, we found that 6 out of 13 chimeric mice had too few I-A^b^ NP_306-322_ tetramer^+^ CD4 TRM in the NT for downstream analysis, from both WT and KO cells (Fig 5J). We speculate that this may be due to a generally low level of donor T cell chimerism achieved at the NT in comparison to that in the lungs (Fig S6B). The finding that NT CD4 TRM are not easily depleted or replenished, consolidate our data about longevity of these cells, i.e. that once they are formed and reside in the NT they will persist. Moreover, we detected an increased accumulation of *Cxcr6*^-/-^ CD4 T cells in the blood, spleen, lung-draining mediastinal lymph nodes (mLN) and URT-draining cervical lymph nodes (cLN), confirming the hypothesis that *Cxcr6*^-/-^ CD4 T cells have migratory defects and/or they are not retained at the tissue (Fig 5K).

CXCL16 is the only known ligand for CXCR6. It exists in both transmembrane and soluble forms and acts as an adhesion molecule and a chemoattractant for leukocytes(Koenen et al., 2017; Wilbanks et al., 2001). In murine lungs, CXCL16 colocalizes with Epcam and its expression is restricted to the large airways lining the lungs(Wein et al., 2019). We investigated CXCL16 expression in NT by microscopy and demonstrated that CXCL16 is expressed in the NT of PR8-OVA infected mice and some OT-II CD4 TRM localize near CXCL16 (Fig 5L, Fig S6C). Finally, to examine the contribution of CXCL16 to the early recruitment of effector CD4 T cells to the NT and lungs, we treated PR8-infected mice with α-CXCL16 antibody (Fig 5M). α-CXCL16 treatment significantly reduced the frequency and number of I-A^b^ NP_306-322_ tetramer^+^ CD4 TRM retained in the NT and, to a lesser extent, in the lungs (Fig 5N-O).

Altogether, we showed that CXCL16 production by airway cells and CXCR6 expression on CD4 T cells are promoting the recruitment of CD4 effector memory T cells to respiratory mucosal tissues and the subsequent establishment of TRM.

### Healthy humans harbor IAV-responsive CD4 TRM in the nasal mucosa

Given the importance of IAV-specific CD4 TRM in the NT of mice, we wanted to investigate whether a similar population exists in healthy human adults. As every individual is exposed to IAV by the age of three(Bodewes et al., 2011), healthy adults must have undergone several symptomatic and asymptomatic infections, and we hypothesized that they should maintain IAV-specific CD4 TRM in their NT. The recruited subjects did not have any respiratory infection or symptoms in the two weeks prior to the study, to avoid contamination from recently activated CD4 T cells. We collected nasopharyngeal swab (NT) and peripheral blood mononuclear cells (PBMC) and analyzed T cells by flow cytometry (Thome et al., 2014) (Fig S7A-B). While the frequency of naïve CD4 T cells (CD3^+^CD4^+^CD45RA^+^CCR7^+^) was higher in PBMCs, CD4 central memory T cells (TCM) (CD3^+^CD4^+^CD45RA^-^CCR7^+^) and CD4 terminally differentiated effector memory T cells re-expressing CD45RA (TEMRA) did not show any difference (Fig S7C). Interestingly, CD4 TEM (CD3^+^CD4^+^CD45RA^-^CCR7^-^) were higher in NT with their subpopulation TRM (CD3^+^CD4^+^CD45RA^-^CCR7^-^ CD69^+^) being exclusively present in NT of all subjects, albeit in varying numbers. When we analyzed both PBMCs and NT, we found that NT CD4 TRM included CD103 (18%) and CXCR6 expressing cells (23%) as previously reported(Ramirez et al., 2024) while PBMC CD4 TEM barely expressed them, similar to our findings in mice (Fig 6A-B).

**Fig 6:**
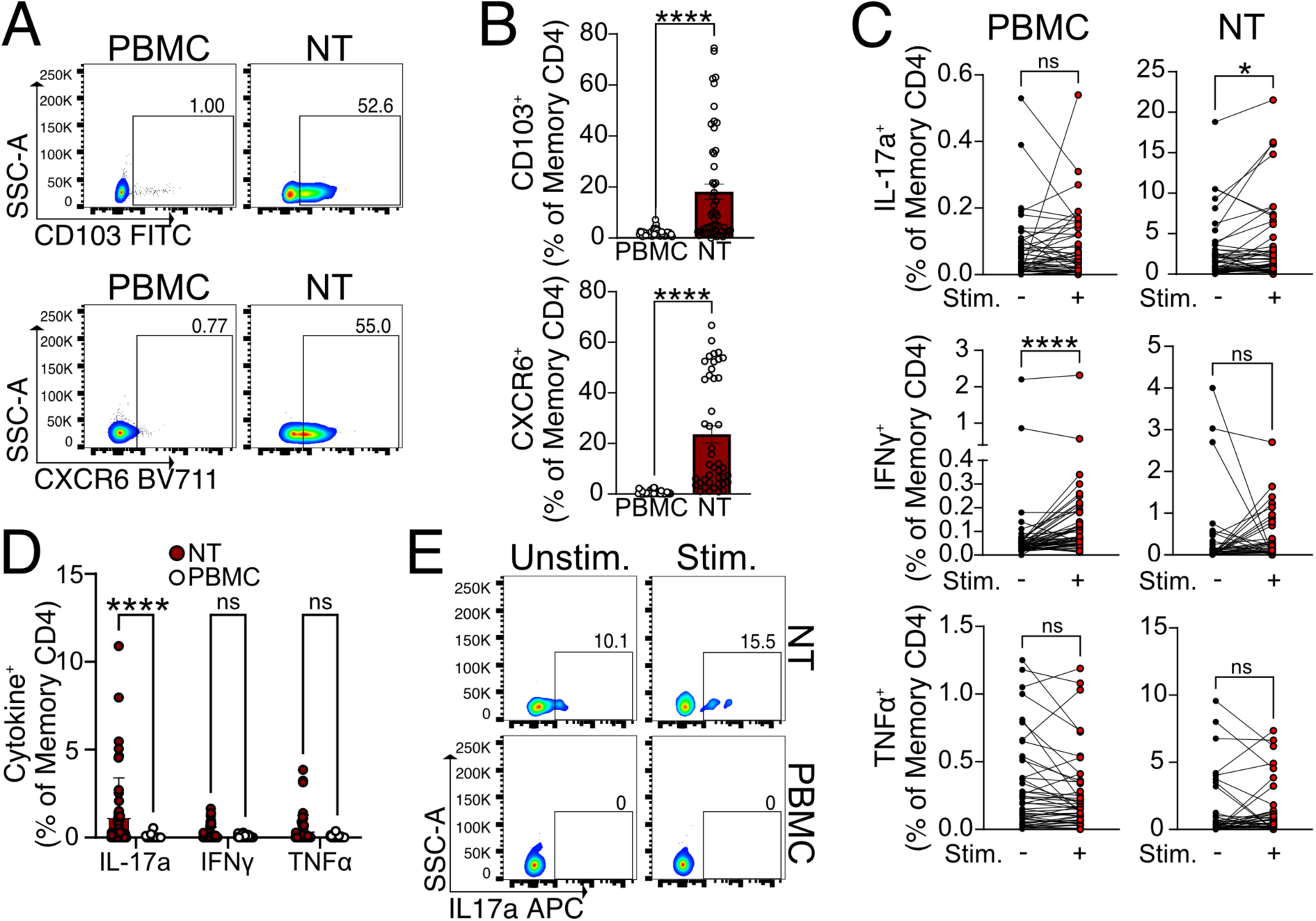
Functional IAV-specific CD4 TRM exist in the nasopharynx of healthy human subjects. A-B) Expression of CD103 and CXCR6 on NT CD4 TRM and CD4 TEM of PBMC derived from healthy human subjects. A) Representative flow cytometry plots indicating the percentage of CD103 and CXCR6 expression on CD4 TRM and CD4 TEM from the same subject. B) Scatter plot showing the percentage of CD103 and CXCR6 expression on CD4 TRM and CD4 TEM. Each data point indicates one subject. The experiment was performed five times and the results (mean ± s.e.m.) are pooled. NS, not significant, \*\*\*\**P*<0.0001; \*\*\**P*<0.001; \*\**P*<0.01; \**P*<0.05 by unpaired two-tailed t-test. C) Percentage of NT CD4 TRM and CD4 TEM from PBMC expressing IL-17a, IFN-γ and TNFα. The cytokine expression is indicated from unstimulated cells and IAV NP peptide pool and M1 peptide pool stimulated cells connected by a line. Each line on the graph indicates cytokine expression from the cells of each subject. NS, not significant, \*\*\*\**P*<0.0001; \*\*\**P*<0.001; \*\**P*<0.01; \**P*<0.05 by two-sided Wilcoxon matched-pairs signed-rank test. D) Scatter plot showing the percentage of NT CD4 TRM and CD4 TEM from PBMC expressing IL-17a, IFN-γ and TNF after stimulation with IAV NP peptide pool and M1 peptide pool (after subtraction of signals from the unstimulated control; negative values considered zero). The experiment was performed seven times and the results (mean ± s.e.m.) are pooled. NS, not significant, \*\*\*\**P*<0.0001; \*\*\**P*<0.001; \*\**P*<0.01; \**P*<0.05 by two-way ANOVA, with Tukey’s multiple comparison test. E) Representative flow cytometry plots showing the expression of IL-17a in NT CD4 TRM and CD4 TEM of PBMC which are stimulated with IAV NP peptide pool and M1 peptide pool or left unstimulated.

A longitudinal study on SARS-CoV-2 vaccinees who had breakthrough infections showed functional NT T cells, specific for SARS-CoV-2 proteins, that persisted for ≥140 days(Lim et al., 2022). Given the relatively long lifespan of NT CD4 T cells, we wanted to examine whether healthy adults retained IAV-specific NT CD4 TRM, acquired by encountering IAV at some timepoint in life. To do that, we stimulated PBMC and NT cells with IAV peptide pools derived from the internal proteins NP and M1 (Matix protein 1) and assessed cytokine expression. We found that PBMC CD4 TEM from 79.5% of the subjects responded to stimulation by secretion of IFN-γ (Fig 6C). Conversely, we did not detect an IFN-γ response in NT CD4 TRM after stimulation. Furthermore, we found no detectable TNF response from NT or PBMCs CD4 T cells (Fig 6C). As the scRNA-seq mouse experiment had revealed high frequencies of Th17 cells in NT (Fig 4E), we also tested if IL-17a expression was induced in stimulated cells. Intriguingly, in NT CD4 TRM the frequency of IL-17a expressing cells increased after stimulation in 63.2% of the individuals. Little to no IL-17a expression was detected in PBMCs before or after stimulation (Fig 6C-E).

Overall, our data show that human NT CD4 TRM share important features with mouse CD4 TRM, such as the expression of CD103 and CXCR6. Moreover, NT of healthy individuals is populated by IAV-specific Th17 CD4 TRM.

### NT Th17-CD4 TRM aid in viral clearance and dampen tissue damage

Given the abundance of Th17 CD4 TRM in NT of mice and human, we wanted to further delineate their functional role. First, we confirmed that PR8-infected mice carried IAV-specific Th17 CD4 TRM in NT by stimulating lung and NT cells with a single immunodominant NP peptide (NP_306-322_) and assessing intracellular expression of IL-17a as a measure of Th17 response. We used NP_306-322_ as this was the same peptide, loaded into tetramers, used to sort CD4 TRM for scRNA sequencing. Consistent with all our data, we found that only NT, but not lung, CD4 TRM from PR8 infected mice expressed IL-17a. Surprisingly, IL-17a production was stimulation-independent although infection-dependent (Fig 7A-B, Fig S8A).

**Fig 7:**
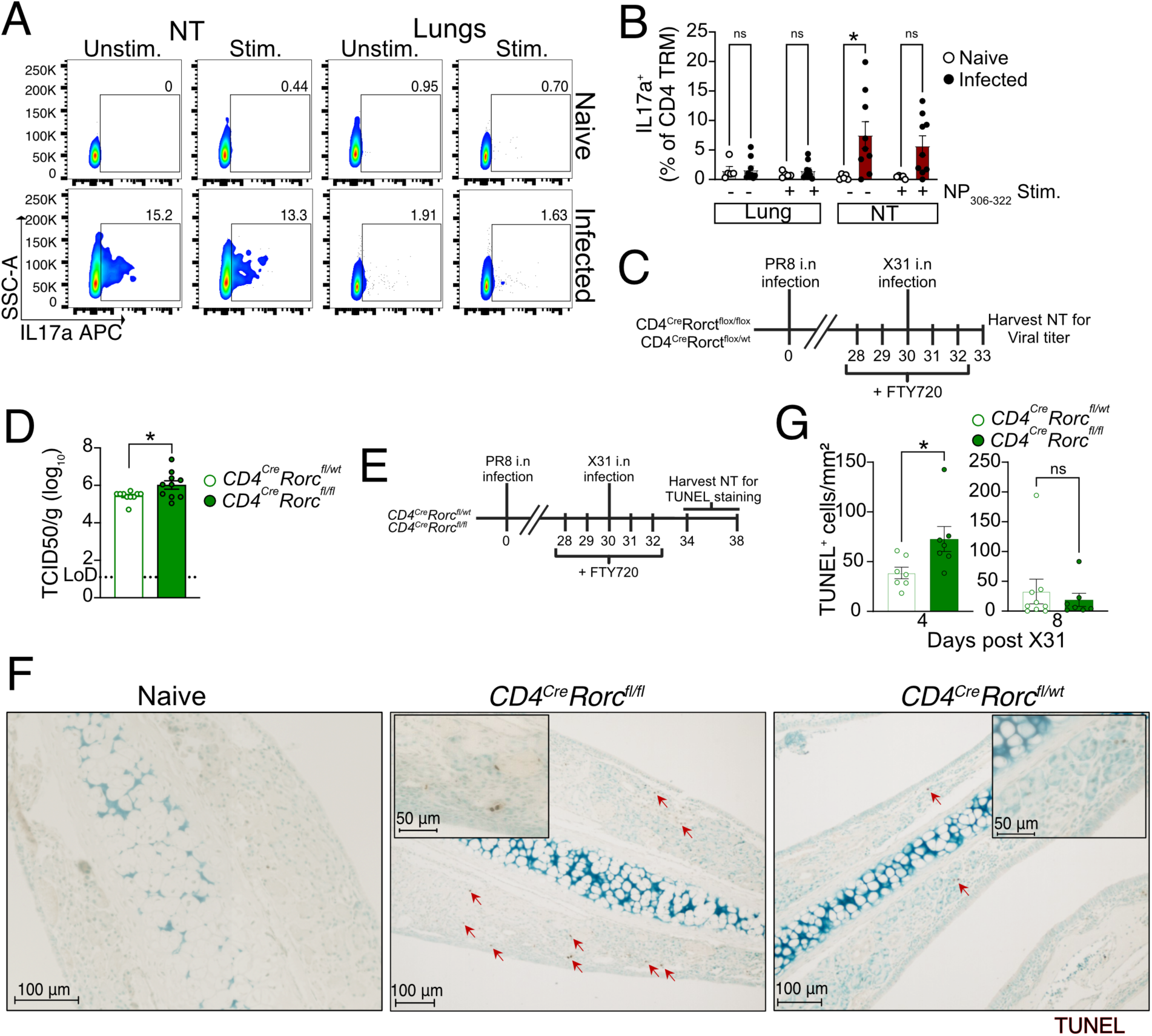
NT harbors Th17 CD4 TRM that reduce local pathology. A-B) Frequency of IL-17a^+^ cells among CD4 TRM of NT and lungs upon restimulation with IAV immunodominant NP_306-322_ peptide in comparison to unstimulated cells. The organs are isolated on day 30 following PR8 IAV infection of mice and from naïve mice. A) A representative flow cytometry plot showing the percentage of IL-17a in lungs and NT. B) Scatter plot indicating the frequency of IL-17a^+^ cells among all CD4 TRM. The experiment was repeated twice and the results (mean ± s.e.m.) are pooled. NS, not significant, \*\*\*\**P*<0.0001; \*\*\**P*<0.001; \*\**P*<0.01; \**P*<0.05 by two-way ANOVA, with Tukey’s multiple comparison test. C-D) Viral titers in the NT of *CD4^cre^Rorct^fl/fl^*and *CD4^cre^Rorc^fl/wt^* mice on day 3 following secondary infection with X31 IAV. The mice were infected with PR8 and reinfected with X31 IAV on day 30 post primary infection and treated with fingolimod ip as indicated. C) A schematic representation of the experimental setup. D) Scatter plot showing viral titers (TCID_50_/g) in the NT of the two groups. The experiment was repeated twice and the results (mean ± s.e.m.) are pooled. NS, not significant, \*\*\*\**P*<0.0001; \*\*\**P*<0.001; \*\**P*<0.01; \**P*<0.05 by unpaired two-tailed t-test. E-G) Microscopy for TUNEL^+^ cells in the nasal septum (respiratory region) derived from *CD4^cre^Rorct^fl/fl^*and *CD4^cre^Rorc^fl/wt^* mice. The mice were infected with PR8 and reinfected with X31 IAV on day 30 following PR8 IAV infection. The tissues were isolated on day 4 and day 8 following X31 IAV infection. E) A schematic representation of the experimental setup. F) Representative microscopic images for TUNEL^+^ cells (brown) in the nasal septum from mice on day 4 post X31 IAV infection are shown. Scale bar: 100µm. G) Scatter plot showing the number of TUNEL^+^ cells per mm2 in the respiratory region of the nasal septum. The experiment was repeated twice and the results (mean ± s.e.m.) are pooled. NS, not significant, \*\*\*\**P*<0.0001; \*\*\**P*<0.001; \*\**P*<0.01; \**P*<0.05 by two-way ANOVA, with Tukey’s multiple comparison test.

Th17 cells recruit neutrophils, B cells, promote transport of secretory IgA to the mucosal sites and protect the hosts in an antibody independent manner during IAV infection(Eliasson et al., 2018; Jaffar et al., 2009; Laan et al., 1999; Wang et al., 2011). On the other hand, detrimental effect of Th17 was also reported as an intranasal immunization triggered Th17 population that caused increased morbidity in mice(Maroof et al., 2014). To examine whether NT Th17 CD4 TRM had a beneficial role, we utilized *CD4^Cre^Rorc^fl/fl^*(CD4-specific Th17 KO) and *CD4^Cre^Rorc^fl/wt^* (WT) mice(Gribonika et al., 2022) (Fig S8B). Mice deficient in *Rorc* gene lack RORγt, a Th17 lineage-specific transcription factor that controls the activation of IL-17A and IL-17F genes, and do not produce Th17 CD4 T cells(Ivanov et al., 2006). To verify the specific role of Th17 CD4 TRM, we infected *CD4^Cre^Rorc^fl/fl^* and *CD4^Cre^Rorc^fl/wt^* mice with PR8 and challenged them at day 30 post primary infection (Fig 7C). FTY720 treatment ruled out the contribution of circulating lymphocytes. CD4 Th17 KO mice had slightly higher viral titer in the NT than did WT mice on day 3 post-secondary infection (Fig. 7D). Furthermore, Th17 CD4 cells have been suggested to play a role in tissue repair and, by secreting anti-apoptotic cytokines that dampen tissue damage(Dutta et al., 2023; Konieczny et al., 2022). We performed a reinfection experiment in a similar way and enumerated apoptotic cells (TUNEL^+^) in the NT upon X31 challenge (Fig 7E). Indeed, *CD4^Cre^Rorc^fl/fl^* mice exhibited an increased number of apoptotic cells in the NT (Fig 7 F-G). However, depletion of IL-17 did not affect viral titer or apoptotic cell clearance in NT (Fig S8C-D) suggesting IL-17-independent mechanisms, mediated by Th17 in the NT.

Altogether, we confirmed that Th17 CD4 TRM ameliorate IAV-related pathology by reducing viral replication and tissue damage.

## DISCUSSION

The current understanding of CD4 TRM during IAV infection relies on findings from lung CD4 TRM. Despite the URT being the first location where IAV encounters the immune system, little is known about CD4 TRM at this crucial immunological site. Here, we defined the phenotype and functional role of CD4 TRM residing in the nasal tissue region of the URT. We demonstrated that antigen-specific CD4 TRM vigorously respond to heterologous IAV challenge, promote direct viral control and provide tissue protection in the URT.

Notably, we found that endogenous antigen-specific CD4 TRM, especially HA-specific ones, are long lived. The HA^+^ CD4 TRM in the NT decayed at a slower rate than CD4 TRMs in the lungs, even though they were less abundant when quantified in absolute numbers. Furthermore, a small fraction of both NP^+^ and HA^+^ CD4 TRM expressed Ki-67 on day 60 suggesting that they may be maintained by homeostatic proliferation like lung CD4 and CD8 TRM(Takamura et al., 2016; Turner et al., 2014). We hypothesize that NALT(Bates, 2022) or ectopic germinal center (GC) in the NT(Gailleton et al., 2025) could be the local sites for naïve CD4 T cell priming and expansion. Indeed, one recent study showed that i.n. vaccination established a CD4 T cell response within the interfollicular regions of the NALT, in turn promoting the homing of antibody-secreting plasma cells to the NT(Liu et al., 2024). Similarly, our recent work indicated that IAV infection and i.n. vaccination induce ectopic GC structures within the NT itself(Gailleton et al., 2025). Furthermore, it should be noted that, while expanded lung clonal families are also found within the NT, the opposite is not true, with all clonal families in the NT with >8 members being organ specific (Fig 4G). However, we could not exclude the possibility of some degree of replenishment from local tertiary lymphoid structures or lymphoid organs via the blood. Overall, our data suggests that few cells traffic from lung draining lymph nodes to NT while the majority instead expand *in situ*.

Even though effector T cells traffic to the infection site due to inflammatory cues, their establishment as TRM may or may not require the presence of antigen depending on the tissue. Antigen-dependent residency has been described for the lungs and brain(Khan et al., 2016; McMaster et al., 2018; Wakim et al., 2010; Zens et al., 2016), but not genital tract or intestinal CD8 TRM, where the inflammatory milieu and chemokines are sufficient for their formation(Bergsbaken and Bevan, 2015; Sato et al., 2014). Even in lungs, antigen-dependency can be bypassed by the introduction of adjuvants, such as zymosan, which evoke a specific inflammatory milieu(Caminschi et al., 2019). Here, we demonstrated that CD4 TRM in NT require the presence of cognate antigen to be formed after IAV infection, which is the opposite of CD8 TRM(Pizzolla et al., 2017a). Furthermore, we showed that the same held true for lung CD4 TRM. The observation in the NT highlights an interesting discrepancy regarding the minimal cues needed by different T cell types to establish residency, even within the same tissue.

Generally, CXCR6 is a chemokine receptor that is more highly expressed on CD8 T cells than on CD4 T cells(Di Pilato et al., 2021; Mabrouk et al., 2022) and is well described as an essential factor for CD8 TRM seeding to the lung airways but not the lung interstitium, after IAV infection (Wein et al., 2019). Here, using bone marrow chimeras and blocking experiments we revealed that the CXCR6-CXCL16 axis is important for the formation of antigen-specific CD4 TRM in NT and lungs. Another study did not find a role for CXCR6 in the recruitment of antigen specific CD4 T cells to the lungs of *M. tuberculosis* infected mice (Ashhurst et al., 2019), however the establishment of TRM was not verified. Nevertheless, a recent sequencing-based study suggested the presence of CXCR6-CXCL16 interactions in the nasal mucosa of IAV-infected mice, using cell-cell communication analysis (Kazer et al., 2024). The authors showed CXCL16 to be expressed by monocyte derived macrophages and KNIIFE cells (Krt13+ nasal immune-interacting floor epithelial) cells. Similarly, we found that cells in the NT express CXCL16 however we did not pinpoint the specific cell population. We suggest two possible mechanisms by which CXCL16 could help recruit CD4 TEM to the NT; either as soluble chemokine, by attracting CD4 TEM or as membrane bound, by retaining the CD4 TRM after migration (Koenen et al., 2017; Wilbanks et al., 2001). Longer depletion experiments or depletion at specific time windows will help cement the role of this chemokine. Still, given the homing of some CXCR6^-/-^ CD4 to the NT, it is plausible that other factors may also be involved in the recruitment and maintenance of TRM.

Functionally, NT CD4 TRM produce cytokines such as IL-2 and IFN-γ upon antigen stimulation. IFN-γ is a cytokine that protects the host by inhibiting viral entry, blocking attachment and replication and stimulating NK cells proliferation during the early phase of infection(Brass et al., 2009; Feeley et al., 2011; Fong et al., 2022; Weiss et al., 2010). Presence of TRM responding to environmental antigens, may explain cytokine production in unstimulated cells. In addition, our scRNA-seq data highlighted the presence of diverse CD4 TRM subtypes, such as cytotoxic CD4 which has had also been suggested to play a role in antiviral defense(Brown et al., 2012; McKinstry et al., 2012). In line with these mechanisms, we found that CD4 TRM are essential, while circulating memory CD4 cells are dispensable, for local protection in the NT. The role of NT CD4 TRM in limiting viral growth was further validated by our i.t. *vs* i.n. priming model that established CD4 TRM in lungs only or in both lungs and NT, respectively. Our results agree with previous work demonstrating that mucosal lung CD4 TRM are better at producing IFN-γ and at protecting against IAV infection than are splenic memory CD4 T cells(Teijaro et al., 2011). CTA1-3M2e-DD-vaccination provided protection against viral challenge; however, the effect size was limited as the URT-restricted infection was only lethal in 43% (6 of 14) of mice. In this context, it was hard to definitively ascribe an effect to CD4 T cells, even if CD4 depletion resulted in a trend towards decreased survival (4 of 15 mice died in CD4 depleted *vs* 1 of 15 in CTA1-3M2e-DD vaccinated). Overall, we demonstrated that NT CD4 TRM have coordinated antiviral and tissue repair functions, depending on antigen specificity. However, it remains to be determined how these distinct protective mechanisms are integrated during a physiological, polyclonal response to infection.

Interestingly, we detected a greater proportion of Th17 CD4 TRM in the NT as compared to lungs. Th17 enrichment was independent of both antigen specificity and infection status. In general, NT harbored a substantial proportion of CD4 memory T cells and very few naïve CD4 T cells. Th17 cells present at steady state could have formed in response to commensal microbiota of naïve mice; it is well known that commensal bacteria play an important role in mucosal immunity and tolerance induction (Liu et al., 2024; Pandiyan et al., 2019). We speculate that the Th17 CD4 TRM in the naïve mice might have an immunoregulatory function similar to that of commensal-specific Th17 in the intestine (Brockmann et al., 2023) or that they may have entered earlier in life, in response to other external stimuli. Importantly, lung Th17 cells have been previously reported to protect from IAV infection in an antibody independent manner(Eliasson et al., 2018; Jaffar et al., 2009; Laan et al., 1999; Lin et al., 2009; McGeachy, 2011; Mills, 2023; Wang et al., 2011). In addition, NT Th17 cells have been shown to form during and protect against several intranasal bacterial infections (Borkner et al., 2021; Solans et al., 2018). All these studies took advantage of *Il17a*-/- mice, which totally lack IL-17a production, and thus have not been able to determine a definitive role for CD4 Th17 cells specifically. Here, we used a recently described conditional knockout, *CD4^Cre^Rorc^fl/fl^* (Gribonika et al., 2022), which allowed us to exclusively dissect the contribution of Th17 CD4 T cells. We demonstrated that NT Th17 CD4 TRM play a role in reducing apoptosis and somewhat in viral clearance upon heterologous IAV infection, independently of IL-17. Th17 CD4 T cells are also well known to produce IL-22 (Liang et al., 2006), which in turn can stimulate the production of anti-apoptotic factors within airways epithelial cells and maintain a healthy tissue (Omokanye et al., 2022; Sonnenberg et al., 2010). In addition, we detected IAV-specific Th17 CD4 TRM in healthy human subjects, suggesting that these specific cells are formed and persist in the NT of healthy adults for years and can be reactivated upon reinfection, similar to what was shown for SARS-CoV-2 and *B. pertussis* (Lim et al., 2022; McCarthy et al., 2024).

Finally, given that nasal respiratory epithelium plays a major role in the transmission of airborne viruses, it is essential to focus on vaccinations that can induce memory cells in the NT(Richard et al., 2020). Although the current intramuscular IAV vaccines are largely effective against IAV infection, they are poor at inducing mucosal immune responses and at protecting against heterologous viruses. Our findings that i.n. CTA1-3M2e-DD immunization can induce M2e-specific CD4 TRM, along with another recent study showing that i.n. Spike vaccination generates antigen specific CD4 TRM in the NT support i.n. immunization as an effective tool for generating functional CD4 TRM in the NT(Diallo et al., 2023b). Nevertheless, immunization strategies to overcome their gradual decay need to be developed.

Collectively, our data provide a comprehensive understanding of NT CD4 TRM that could be leveraged to design mucosal vaccines aimed at generating NT CD4 TRM providing heterosubtypic protection.

## MATERIALS AND METHODS

### Mice

We conducted the experiments according to protocols approved by regional animal ethics committee in Gothenburg (ethical permit numbers: 1666/19, 2230/19 3307/20 and 38/23). C57BL/6N and BALB/c mice were purchased from Janvier labs, France and Taconic Biosciences, Denmark. They were housed in the specific pathogen free animal facility of Experimental Biomedicine Unit (EBM) at the University of Gothenburg. B6.SJL- *Ptprc^a^ Pepc^b^*/BoyJ (CD45.1^+^) and B6.129P2-*Cxcr6^tm1Litt^*/J (*Cxcr6*^-/-^) mice were purchased from the Jackson Laboratory, USA and were bred at EBM. Ovalbumin-specific CD45.1^+^ OT-II, tdtomato^+^ OT-II mice TCR transgenic, *CD4^Cre^Rorc^fl/wt^* and *CD4^Cre^Rorc^fl/fl^*(Gribonika et al., 2022) were bred at EBM at the University of Gothenburg. We used both female and male mice which were eight to twelve weeks old in the experiments.

### Sampling of human subjects

Written consents were obtained from the study participants before obtaining the samples. Ethical permit (ethical permit number: 2023-07055-01) was obtained by the Swedish Ethical Review Authority to collect paired venous blood and nasopharyngeal swabs from healthy subjects aged between 18-60 years old. Both male and female volunteers were included. We excluded the subjects who fulfilled any of the following criteria: respiratory infection during the last two weeks, allergic rhinitis, chronic obstructive pulmonary disease, asthma, regular nose bleeding, immunodeficiency and IAV infection in the last two years (verified by PCR).

8 ml of venous blood was collected into lithium heparin coated tubes and was subjected to density gradient centrifugation to obtain peripheral blood mononuclear cells (PBMC) using lymphoprep (Stemcell Technnologies) according to manufacturer’s instructions.

Nasopharyngeal cells (NT) were collected into 3 ml RPMI containing 2% fetal calf serum (FCS) and 1.5 mM DTT as previously described(Lim et al., 2022). For swab collection the volunteers were seated and sample collected from both nostrils using the same Eswabs (Copan, #482C). Swabs were inserted along the floor of the nasal cavity and rotated in circular motion. Either posterior nasopharynx or inferior nasal turbinates were sampled from different individuals. The cells were vortexed for 1 min and were incubated at 37°C for 30 min. The cells were spun at 700xg for 8 min at 4°C and the cell pellets were suspended in FACS buffer.

In total 60 paired samples were analyzed but data from 10 of them was excluded for the cytokine, CXCR6 and CD103 analysis because of low recovery of CD4 TRM from NT or PBMC.

### Infection of mice

Mouse adapted Influenza A/Puerto Rico/8/34 (PR8) (molecular clone; H1N1) and Influenza A/HK/x31 (HKx31)) (molecular clone; H3N2) were grown in 10 day old embryonated chicken eggs. The allantoic fluid was harvested, and the viral titer was determined using TCID_50_ assay. PR8 containing OVA_323-339_ (PR8-OVA)(Thomas et al., 2006) was kindly gifted by Paul G. Thomas, St. Jude Children’s Research Hospital, USA. For sublethal total respiratory tract infection (TRT infection), mice were briefly anesthetized using isoflurane and inoculated intranasally with a sublethal dose (50 TCID_50_) of mouse adapted Influenza A/Puerto Rico/8/34 (PR8) (Molecular clone; H1N1) diluted in HBSS containing 0.1% BSA (total volume of 25 µl). To enhance the amount of CD4 TRM in the NT, mice were infected intranasally with a high dose (10^5^ TCID_50_) of PR8 or PR8-OVA in a volume of 5 µl (URT restricted infection) 30 minutes after the TRT infection.

For rechallenge experiment after PR8 infection, a lethal dose of 5000 TCID_50_ of X31 strain was used. For rechallenge experiment after CTA1-3M2e-DD immunization, mice were infected intranasally with a high dose (10^5^ TCID_50_) of PR8 in a volume of 10 ul in a URT restricted manner.

For intratracheal infection, 500 TCID_50_ of PR8 diluted in HBSS containing 0.1% BSA (total volume of 25 µl) is inoculated intratracheally after anesthetizing the mice.

### Immunization of mice

5µg CTA1-3M2e-DD fusion protein was diluted in 1X PBS to an end volume of 5µl. BALB/c mice were immunized with CTA1-3M2e-DD intranasally and boosted twice with a ten-day interval. Control group was immunized with 5µg CTA1-DD adjuvant similarly.

### Blocking of CXCL16, IL17A/F and depletion of T cells *in vivo*

Mouse CXCL16 antibody (MAB503) was purchased from R&D systems and was dissolved in 1X PBS. The antibody was administered intraperitoneally at a dosage of 2.5µg/g weight of mouse on day 1, 3, 6 and 9 post infection with PR8. Mice treated with equal amount of *InVivo*MAb rat IgG2a isotype control, anti-trinitrophenol (BE0089, Bioxcell) were used as controls. For depletion of IL-17A and IL-17 F, *InVivo*MAb anti-mouse IL-17A (BE0173) and *InVivo*MAb anti-mouse IL-17F (BE0303) were purchased from Bioxcell and administered intraperitoneally. The mice received 400 ug of each antibody on day 29 and 200ug of each antibody from day 30-32/33 post PR8 infection. The control group received equal amount of *InVivo*MAb mouse IgG1 isotype control, unknown specificity (BE0083) (Bioxcell).

*InVivo*MAb anti-mouse CD8α (BE0117), *InVivo*MAb anti-mouse CD4 (BE0003-1) and *InVivo*MAb rat IgG2b isotype control (BE0090) were purchased from Bioxcell and were administered intraperitoneally at a daily dosage of 10µg/g of mouse from day 28 to day 32 following PR8 infection.

### Fingolimod treatment

10mg/ml stock of fingolimod (Cayman Chemicals) was prepared in DMSO. We administrated fingolimod at a dosage of 2.5 µg/g weight of mouse dissolved in sterile physiological saline solution by intraperitoneal injection at indicated days.

### Adoptive transfer of antigen specific T cells

For adoptive transfer of OT-II^+^ CD4 T cells, CD45.1^+^ OT-II or tdTomato^+^ OT-II mice were euthanized, and spleens were collected. Spleens were mashed and made into single cell suspension by passing through a 70µm strainer. Total CD4 T cells were isolated using EasySep mouse CD4^+^ T cell isolation kit (Stemcell technologies) according to manufacturer’s instruction. A total of 2.5-5x10^6^ OT-II^+^ CD4 T cells suspended in 1X PBS were administrated by intravenous injection four hours prior to infection with PR8-OVA.

For generation of effector OT-II T cells, we isolated CD4 T cells from splenocytes of tdtomato^+^ OT-II mice. OT-II T cells were stimulated by culturing with splenocytes pulsed with 1uM OVA peptide ISQAVHAAHAEINEAGR in 5:1 ratio respectively in RPMI media containing 10% FBS, 10µg/ml gentamycin, 55µM β-mercaptoethanol (Gibco) and 20U/ml murine IL-2 (Peprotech) for 60h in 5% CO_2_ cell culture incubator. The cells were washed and incubated with fluorochrome conjugated antibodies against surface antigens-CD3 PerCPCy5.5 (17A2), CD4 APC-Cy7 (RM4-4) and CD69 BV605 (H1.2F3) for 20 min at 4°C. To exclude dead cells, the cells were washed and stained with LIVE/DEAD™ Fixable Aqua Dead Cell Stain (cat. no: L34957, Invitrogen) according to manufacturer’s instruction. Cells were suspended in FACS buffer and live CD3^+^CD4^+^CD69^-^ dtomato^+^ OT-II effector T cells were collected after sorting in a BD FACSAria fusion (BD Biosciences). A total of 5x10^5^ effector OT-II^+^ CD4 T cells suspended in 1X PBS were administrated by intravenous injection 60 h post infection with PR8 or PR8-OVA.

### Bone marrow chimera

For myeloablation, CD45.1^+^ B6.SJL-*Ptprc^a^ Pepc^b^*/BoyJ mice were irradiated with 11 Gy in equally split doses with a 4-hour gap in a RS 2000 X ray irradiator machine. The mice were rested for a day and transplanted with a mix of 2.5 x 10^6^ tdtomato^+^ WT and 2.5 x 10^6^ CD45.2^+^ *Cxcr6*-/- bone marrow cells (BM) by intravenous administration. The recipients with newly reconstituted BM (BM chimera) were infected with 50 TCID_50_ PR8 intravenously 70 days after transplantation.

### Isolation and processing of blood, lungs, NALT, nasal tissue and Peyer’s patches

For flow cytometry experiments with cells from blood, lungs, NALT and nasal tissue, mice were injected with CD4 FITC or CD4 APC-Cy7 (RM4-4) antibody intravenously to label the circulating CD4 T cells. The mice were completely anesthetized, and blood was withdrawn into EDTA coated tubes from the heart after five minutes of *in vivo* antibody labelling. The lung was perfused with 1X PBS and processed into single cell suspension using the mouse lung dissociation kit (Miltenyi Biotec). Blood was treated with RBC lysis buffer for five minutes, washed and resuspended in FACS buffer. Nasal tissue and NALT were isolated and processed immediately as previously described(Pizzolla et al., 2017a). For isolation of nasal tissue and NALT, the mouse head was collected and deskinned. Briefly, NALT was obtained by peeling the upper palate using a forceps and the cells were extracted by teasing the NALT between two frosted slides. The rest of the head was cut coronally to remove the brain and the incisors were removed by scissors. The nasal septum, nasoturbinates, ethmoid turbinates, vomeronasal organs, olfactory recess, maxilloturbinates including cartilages and bones were collected which is referred here as nasal tissue (NT). Nasal tissue was digested in 3 ml RPMI 1640 media supplemented with 2% FBS, 6 mg collagenase type 3 (Worthington Biochemical corporation) and 600 µg DNase I, grade II from bovine pancreas (Sigma Aldrich) in a shaking incubator at 37°C for one hour. The cells were made into single cell suspension by straining through a 70 µm nylon mesh. The cells were washed and resuspended in FACS buffer or supplemented media for downstream processing.

For processing of cells from Peyer’s patches (PPs), around 3-5 PPs were isolated from the small intestine and collected in HBSS media. The PPs were washed twice with HBSS supplemented with 1% FBS, 15 mM HEPES and 5 mM EDTA by incubation at 37°C for 15 min with occasional shaking. The PPs were mashed, and the cells were made into single cell suspension by straining through a 70 µm nylon mesh. The cells were washed and resuspended in FACS buffer for staining.

### Stimulation of mouse cells with viral peptides

The cells obtained from the organs of naïve and infected mice were stained for extracellular markers prior to stimulation with peptides. The following IAV peptide array were obtained from obtained from BEI resources, NIAID, NIH: Influenza Virus A/New York/348/2003 (H1N1) Nucleoprotein (NP) (NR-50714), A/New York/444/2001 (H1N1) Nonstructural protein 1 (NS1), Influenza Virus A/Puerto Rico/8/1934 (H1N1) Hemagglutinin (HA) (NR-18973), Influenza Virus A/Puerto Rico/8/1934 (H1N1) Neuraminidase (NA) (NR-19257) and SARS-Related Coronavirus 2 Nucleocapsid (N) (NR-52404). 1µg each of all peptides from the peptide array were mixed to obtain the respective peptide pools.10 million cells derived from lungs and nasal tissue were stimulated with IAV NP, NS1, HA and NA peptide pools containing 1 µg of each peptide pool in RPMI 1640 supplemented with 10% FBS, 1% Penicillin/Streptomycin and Protein Transport Inhibitor (Containing Brefeldin A) (diluted 1:1000) (BD, 555029). The cells were incubated for four hours at 37°C and further washed with 1x PBS to proceed for intracellular cytokine staining.

For comparison of intracellular cytokines produced by CD4 TRM that are stimulated by IAV NP or SARS-Related Coronavirus 2 Nucleocapsid (N) Protein (SARS-CoV-2 N), live CD3^+^CD4iv^-^CD4 tissue^+^ CD44^+^CD62L^-^CD69^+^ CD4 TRM cells were sorted from lungs and NT 30- and 60-days post infection. The CD4 TRM (CD45.2^+^) were co-cultured with congenic CD45.1^+^ splenocytes pulsed with IAV NP peptide pools containing 1ug of each peptide from peptide array (NR-50714, BEI Resources) or with SARS-CoV-2 N peptide pools containing 1ug of each peptide from peptide array (NR-52404, BEI Resources) for 18 hours at 37 deg in RPMI 1640 supplemented with 10% FBS, 1% Penicillin/Streptomycin. The culture was supplemented with Protein Transport Inhibitor (Containing Brefeldin A) (diluted 1:1000) at the 12^th^ hour. The cells were further washed with 1x PBS to proceed for intracellular cytokine staining.

### Stimulation and flow cytometry of human PBMC and nasopharyngeal cells

PBMC and NT cells were incubated with Human Trustain FcX (Biolegend, 422302) for 10 min at 4°C. The cells were then incubated with anti-human CD69 BV421 antibody (clone: FNXO) for 20 min at 4°C and washed with FACS buffer. The cells were finally resuspended in T cell stimulation media (RPMI 1640 supplemented with 10% FBS, 1% Penicillin/Streptomycin, Protein Transport Inhibitor (Containing Brefeldin A) (diluted 1:1000) (BD, 555029), Monensin (Biolegend, 420701) and anti-human CD28/CD49d (BD, 347690)(Sekine et al., 2020). The cells were stimulated with peptide pools containing 2ug each of all peptides from Influenza Virus A/New York/348/2003 (H1N1) Nucleoprotein (NP) (NR-50714) peptide array and Influenza Virus A/New York/348/2003 (H1N1) Matrix protein 1 (M1) (NR-2613) peptide array for 6 hours at 37°C. Cells without the peptide pools (peptide diluent) were kept as unstimulated controls. After washing with FACS buffer, the cells were stained for extracellular antigens with the following fluorochrome conjugated antibodies (clone used indicated in bracket): anti-CD3 BUV737 (UCHT1), anti-CD4 PE-Dazzle594 (OKT4), anti-CD45RA BV605 (HI100), anti-CCR7 APC-Cy7 (G043H7), anti-CD103 FITC (Ber-ACT8) and anti-CXCR6 (13B1E5). After incubation at 4°C for 20 min, the cells were washed and stained with LIVE/DEAD™ Fixable Aqua Dead Cell Stain (cat. no: L34957, Invitrogen) according to manufacturer’s instruction to exclude dead cells. To detect intracellular cytokines, the cells were fixed with 4% paraformaldehyde for 20 min at RT, washed with ice cold 1X PBS and permeabilized with permeabilization buffer (1X PBS containing 0.5% BSA and 0.5% saponin) for 10 min at RT. The cells were washed and resuspended in permeabilization buffer with the following fluorochrome conjugated antibody cocktail: anti-IFN-γ PE (25723.11), anti-TNF BV650 (MAb11) and anti-IL-17a APC (BL168). After incubation at 4°C overnight, the cells were washed and suspended in FACS buffer for data acquisition at BD LSR Fortessa X-20 or Sony spectral analyzer.

### Flow cytometry of murine cells

All fluorochrome-conjugated antibodies are titrated to determine the optimal concentration for flow cytometry. Single cell suspension from organs were prepared and resuspended in FACS buffer as mentioned before. The following antibodies are used for mouse extracellular antigen staining: anti-CD3 PerCPCy5.5 (17A2), anti-CD4 FITC (RM4-4), anti-CD4 APC-Cy7 and BUV496 (RM4-5), anti-CD8 PE-Cy7 (53-6.7) anti-CD44 BV786 (IM7), anti-CD44 PE-Dazzle 594 (IM7), anti-CD62L BV711 and APC (MEL-14), anti-CD69 BUV737 (H1.2F3), anti-CD11a PE-Cy7 (H155-78), anti-CD103 PE-Cy7, BV421 and BV711 (2E7), anti-CD45.1 APC (A20), anti-CD45.2 BUV395 (104), anti-CXCR6 BV421 (SA051D1), anti-ICOS BV421 (7E.17G9), anti-PD1 PE (J43), anti-CD19-Alexa Flour 700 (6D5), anti-CD11b-Alexa Flour 700 (M1/70). The extracellular staining was performed by incubating the cells with the antibodies for 20 minutes at 4°C. For labelling antigen specific cells with fluorochrome conjugated tetramers, cells were incubated with the tetramers for 2h 40 min at 4°C (I-A^b^ NP_306-322_ APC and I-A^b^ HA_91-107_ PE/BV421 tetramers) or 1 h at room temperature (I-A^d^ M2e_2-17_ PE) and the extracellular antibody mix was added onto the top and incubated for 20 minutes. The following tetramers kindly provided by NIH tetramer core facility at Emory University were used: I-A^b^ NP_306-322_ APC, I-A^d^ M2e_2-17_ PE, I-Ab HA_91-107_ PE and I-Ab HA_91-107_ BV421. I-A^b^ human CLIP_87-101_ conjugated with APC, PE or BV421 and I-A^d^ human CLIP _87-101_ conjugated with PE were used as negative controls. To exclude dead cells, the cells were washed and stained with LIVE/DEAD™ Fixable Aqua Dead Cell Stain (cat. no: L34957, Invitrogen) according to manufacturer’s instruction. The cells were fixed with IC fixation buffer (eBioscience, 00-8222) for intracellular cytokine staining, or fixed with Foxp3 Transcription Factor Fixation/Permeabilization buffer (eBioscience 00-5523) for transcription factor staining for 60 minutes at room temperature. The cells were washed and resuspended in 1X permeabilization buffer (eBioscience, 00-8333) and incubated with the following antibodies: anti-IL-2 BV785 (JES6-5H4), anti-TNF BV421 (MP6-XT22), anti-IFN-γ BV605 (XMG1.2), anti-IL-17a PE or FITC (TC11-18H10), anti-RORγt PE-CF594 (Q31-378) and anti-Ki-67 V450 (B56). The cells were washed with 1X permeabilization buffer and resuspended in FACS buffer. The labelled cells were run, and the data was acquired on the BD LSR Fortessa X-20 or BD LSR II (BD Biosciences) flow cytometer or Sony ID7000 Spectral cell analyzer and was analyzed using Flow-Jo software (Tree Star).

### Immunofluorescence staining

Mice were euthanized and upper part of mice heads were collected after removing the skin and eyeballs. The heads were fixed with 1.5% Formaldehyde Solution (w/v) (ThermoScientific) solution at 4°C for 24 hours. After removing the teeth and remaining muscle tissues, they were transferred into 0.5M EDTA solution for decalcification for 7 days at 4°C. The samples were embedded into sequential dilutions of OCT paramount (Histolab) and were snap frozen for sectioning. 10-14 μm thick frontal sections thick sections were cut and SuperFrost microscopy slides and stored at -80°C or were processed for staining. In the latter case, slides were baked for 1 hour at 60°C. Antigen retrieval was performed on nasal tissue sections mounted on glass slides to enhance epitope accessibility for immunofluorescence staining. Slides were submerged in a sodium citrate retrieval buffer consisting of 10 mM sodium citrate and 0.05% Tween-20, adjusted to pH 6.0. Heat-induced epitope retrieval was carried out by placing the slides in the buffer and heating them in a microwave until the solution reached 100 °C. Once the target temperature was reached, the slides were removed from heat and allowed to cool gradually to room temperature for approximately 1 hour. Following cooling, slides were fixed with ice-cold acetone until the samples were dried. The tissue was blocked with 1X PBS containing 10% FBS and 5% Normal Rat Serum for 15 min at RT. After washing with 1X PBS, the tissues were stained with antibody cocktail overnight at 4°C in the dark. The following antibodies were used: anti-CXCL16 (Uncoupled Polyclonal, #bs-1441R-TR Biosciences), anti-CD4 (FITC RM4-4, #116004 Biolegend), anti-CD44 (APC IM7, #103012 Biolegend), anti-CD45.1 (PE A20, #110707 Biolegend), Goat anti-Rabbit IgG (Texas Red, #sc-2780 Santa Cruz Biotech).The sections were counterstained by incubating with Hoechst 33342 solution (ThermoFischer Scientific) for 15 min at RT in the dark. The sections were mounted with a fluorescent mounting medium (Dako), acquired on an Axio Imager. Z2 (Zeiss) and analysed on TissueFACS Analyser (Version 7.1.120 Xylis).

### TUNEL assay

Mice were euthanized and upper part of mice heads were collected after removing the skin and eyeballs. The heads were fixed with 4% Formaldehyde Solution (w/v) (ThermoScientific) solution at 4°C for 24 hours. After removing the teeth and remaining muscle tissues, they were transferred into 14% EDTA solution (w/v) for decalcification for 14 days at 4°C. The tissues were sequentially dehydrated and embedded into paraffin wax in a cassette. 4 μm thick sections were cut from the paraffinized tissue blocks. TUNEL assay kit -HRP-DAB (Abcam, ab206386) was used to detect apoptotic cells according to manufactureŕs instructions. The images were Axio Imager. Z2 (Zeiss) and analysed on TissueFACS Analyser (Version 7.1.120 Xylis) and QuPath (version 0.5.1) software.

### Neutralization assay

200000 MDCK cells were plated in a 24 well plate and was incubated overnight at 37°C. Serum from IAV infected mice or mice immunized with irrelevant protein (streptavidin or SARS-CoV-2 spike protein) was serially diluted (two-fold) and incubated with 10^6^ TCID_50_ X31 (MOI=5) in 500µl DMEM supplemented with 1mM HEPES, 1µg/ml TPCK trypsin and 5µg/ml gentamycin for 1 hour at 37°C. The plate containing MDCK cells was washed twice with 1x PBS and the virus/serum mixture was added onto the cells. The plate was incubated for 18 hours and washed twice with 1x PBS. The cells were incubated with 5x trypsin for cell detachment and the washed cells were fixed with fixation/permeabilization solution (eBioscience™ Foxp3 / Transcription Factor Staining Buffer Set, 00-5523-00) overnight at 4°C. The cells were washed twice with FACS buffer containing 0.1% saponin. The infected cells were labelled by incubating cells with InVivoMAb anti-Influenza A virus NP (clone H16-L10-4R5 (HB65)) (Bioxcell) labelled with alexa fluor 488 (Protein Labeling Kits for alexafluor 488, Invitrogen, A10235) for 30 minutes at 4°C. The cells were washed twice with FACS buffer containing 0.1% saponin and resuspended in FACS buffer for flow cytometry. The average signals from cells incubated with virus in the absence of serum was considered as 100% infectivity and is used to calculate the relative infectivity of cells in presence of serum.

### Viral titer determination

Lung and URT (nasal tissue and NALT) are harvested and suspended in 1ml 1x PBS per 0.5g of organ. The tissues were homogenized using a tissue homogenizer and spun at 1200xg for 10 min at 4°C. The supernatant was used for determining the viral titer using the standard TCID_50_ assay.

### Single cell RNA sequencing

Mice were injected with CD4 FITC (RM4-4) antibody intravenously to label the circulating CD4 T cells (referred as CD4 iv). The mice were completely anesthetized, and the lung was perfused with 1X PBS. Nasal tissue and lungs were isolated, digested and processed into single cells as described before. The samples were enriched for CD4 T cells using the EasySep™ mouse CD4+ T Cell isolation kit (Stemcell technologies) and the dead cells were removed using EasySep™ Dead Cell Removal (Annexin V) Kit (Stemcell technologies) according to manufacturer’s instruction. Cells from PR8 infected mice were labelled with the following fluorochrome-conjugated MHC-II tetramers: I-Ab NP_306-322_ APC and I-Ab HA_91-107_ BV421 as described before. After washing with FACS buffer, cells were labelled with TotalSeq™-C0987 anti-APC antibody (408007, Biolegend) to identify NP-specific CD4 TRM. Cells were washed and labelled with antibodies. The following fluorochrome labelled antibodies were used: CD3 PerCPCy5.5 (17A2), CD4 APC-Cy7 (RM4-4) (for labelling tissue resident CD4 Tc-referred as CD4 tissue) and CD69 BV605 (H1.2F3) and CD44 PE-Dazzle 594 (IM7) and CD62L BV711 (MEL-14). Additionally, totalSeq-C0595 anti-mouse CD11a antibody (101131, Biolegend) and the following barcoded hashtag antibodies were used: hashtag antibodies 3,5-9 (155865, 155869, 155871, 155873, 155875, 155877; Biolegend) for demultiplexing of samples during data processing and to detect cells expressing CD11a protein on the cell surface. Live total CD4 TRM (CD4 iv^-^ CD4 tissue^+^CD44^+^CD62L^-^CD69^+^) were sorted from PR8 infected and naïve mice while NP-specific CD4 TRM (NP_306-322+_ CD4 iv^-^ CD4 tissue^+^CD44^+^CD62L^-^CD69^+^) and HA-specific CD4 TRM (HA ^+^ CD4 iv^-^ CD4 tissue^+^CD44^+^CD62L^-^CD69^+^) were sorted from PR8 infected mice. In addition to CD4 T cells, we sorted total live cells devoid of CD4 T cells from the NT of naïve and PR8 infected mice. The sorted cells were collected in a BD FACSAria fusion (BD Biosciences) cell sorter and processed immediately. The total number of cells sorted from each group are the following. Lung total CD4 TRM from naïve mice: 8000 cells, NT total CD4 TRM from naïve mice: 3095 cells, lung NP-specific CD4 TRM: 4002 cells, NT NP-specific CD4 TRM: 1172 cells, lung HA-specific CD4 TRM: 4073 cells, NT HA-specific CD4 TRM: 1314 cells, lung total CD4 TRM from infected mice: 8000 cells, NT total CD4 TRM from infected mice: 3000 cells, live cells except CD4 T cells from NT of naïve and infected mice: 10000 cells each. The sorted cells were pooled and washed in the Laminar Wash Mini system (Curiox Biosystems).

A maximum of 20,000 cells were processed into single cells in a chromium controller (10X Genomics). The single cell gene expression, TCR VDJ and cell surface protein (CSP) libraries were prepared using Chromium Next GEM Single Cell 5’ Reagent Kits v2 (Dual Index) kit (PN-1000265), library construction kit (PN-1000190), 5’ Feature Barcode Kit (PN-1000256), Chromium Single Cell Mouse TCR Amplification Kit (PN-1000254), Chromium Next GEM Chip K Single Cell Kit (1000286), Dual Index Kit TT Set A (PN-1000215) and Dual Index Kit TN Set A (PN-1000250). The prepared cDNA libraries were quantified using Qubit Fluorometer (Invitrogen) and quality was assessed using Agilent Tapestation system. The libraries were sequenced on Illumina NovaSeq 6000 by SNP&SEQ Technology Platform, Science for Life Laboratory (Uppsala Biomedical Centre, Uppsala University, Sweden) according to sequencing instructions provided by 10x Genomics.

### Single cell RNA seq data processing

Raw fastq files were processed through the 10X cellranger pipeline using the multi command and default parameters with reference genome GRCm38-mm10. Raw UMI count matrices generated from the cellranger 10X pipeline were loaded and merged into a single Seurat object (Seurat version 4)(Hao et al., 2021; Hao et al., 2024; Satija et al., 2015). Cells were discarded if they met any of the following criteria: percentage of mitochondrial counts > 15%, number of unique features or total counts was in the bottom or top 0.5% of all cells, and number of unique features < 250. Gene counts were normalized to the same total counts per cell and were natural log transformed (after the addition of a pseudocount of 1). The normalized counts in each cell were mean-centered and scaled by their SD, and the following variables were regressed out: percentage of mitochondrial counts, percentage of ribosomal counts, G2M and S phase scores, TCR gene expression scores. Hashtags and APC-surface antibody were normalized by a centered log-ratio (CLR) normalization and binding assigned using HTODemux function. Data integration across cells originating from different samples was done using the Anchor method within Seurat. Selection of the number of components for the nearest-neighbor network computation was based on their visualization in an elbow plot. UMAP(Becht et al., 2018) was performed for the spatial visualization of the single-cell data set after features and cells were clustered using the Louvain algorithm. TCR data was processed using scRepertoire package(Borcherding et al., 2020) with standard setting. T cell clonality was based on identical nucleotide sequence in the CDR3 region. All TCR information were integrated with the scRNA-Seq data by aligning and merging the data with the metadata slot in the processed RNA-Seq Seurat object. Differentially expressed genes between different clusters were identified using the FindAllMarkers function from Seurat using default settings (Wilcoxon test and Bonferroni P value correction). Significant genes with average log fold change > 0.25 and expressed in > 25% of cells in that group were ranked according to fold change and are reported in table S1. For comparison between 2 groups, the FindMarkers function was used, and either top and bottom 20 genes were represented by Heatmap using the DoHeatmap function of Seurat or all genes were represented on a VolcanoPlot using the EnhancedVolcano R package(Blighe K, 2024). Data underlying the volcano plot is in Table S1. Gene signatures were added to the Seurat object using the AddModuleScore function.

### Statistical analysis

The data was analyzed using GraphPad prism software. The experiments were performed and analyzed in a non-blinded fashion. Both male and female mice were included in the experiments. For statistical analysis, unpaired two-tailed *t-*test, one-way ANOVA, with Dunnet’s multiple comparison test, two-way ANOVA, Tukey’s multiple comparison test or two-sided Wilcoxon matched-pairs signed-rank test and is indicated in the figure legends. Normality and lognormality was tested using GraphPad prism. Differences were considered significant when the P-value was <0.05. Outliers were identified and removed using Grubbs method (alpha=0.05) when indicated in the figure legend. For weight loss analysis, we used a linear mixed effects analysis of the body weight data (normalized to the initial weight of each individual animal) in R (www.r-project.org). As fixed effects, we used the treatment and the day (with an interaction term between those fixed effects). As random effects we had intercepts for the individual animals. Thereafter, we compared the weight curves across treatment groups (averaged over time) using estimated marginal means and pairwise comparisons with Tukey’s Honest Significant Difference (HSD) adjustment. To assess treatment group differences at each time point, we used the same linear mixed-effects model described above. Estimated marginal means were computed for each treatment group at each individual day (0–14). Pairwise comparisons between treatment groups were performed at each time point, and p-values were adjusted for multiple testing using Holm’s method.

## Acknowledgments

We would like to thank the staff at the Experimental Biomedicine (EBM) core facility at the University of Gothenburg for animal management; Paul G. Thomas, St. Jude Children’s hospital for PR8-OVA virus; Linda M Wakim, Doherty Institute for providing protocol for processing of nasal tissue; Carina Karlsson, University of Gothenburg for providing access to microtome; Alvar Almstedt, Bioinformatics and Data Centre (BDC), Core Facilities, University of Gothenburg, for data pre-processing; Genomics core facility at Sahlgrenska Hospital for providing access to Qubit fluorometer and tapestation and study participants for providing samples. MHC-II tetramers were obtained from the NIH tetramer core facility. The following reagents were obtained through BEI Resources, NIAID, NIH: Peptide Array, Influenza Virus A/New York/348/2003 (H1N1) Nucleocapsid Protein, NR-5071, Peptide Array, Influenza Virus A/New York/348/2003 (H1N1) Matrix protein 1 (M1) (NR-2613) and Peptide Array, SARS-Related Coronavirus 2 Nucleocapsid (N) Protein (NR-52404). Sequencing was performed by the SNP&SEQ Technology Platform in Uppsala. The facility is part of the National Genomics Infrastructure (NGI) Sweden and Science for Life Laboratory. The SNP&SEQ Platform is also supported by the Swedish Research Council and the Knut and Alice Wallenberg Foundation. The experiment schemes are created with Biorender.com.

## Author contributions

Conceptualization: NRM, DA. Methodology: NRM, RG, LS, KS, AS, MB, KT, DA. Formal analysis: NRM, RG, LS, KS, JE, HA, AS, NL, MB, KT, DA. Investigation: NRM, RG, LS, KS, JE, HA, MB, KT, DA. Data curation: DA, NRM. Software: DA, NRM. Resources: NL, MB. Project Administration: NRM, DA. Supervision: MB, NL, KT, DA. Funding Acquisition: DA, NRM. Writing – original draft: NRM, DA. Writing – review & editing: NRM, RG, LS, KS, JE, HA, AS, MB, KT, DA.

## Data availability

Single-cell RNA-seq data have been deposited in the NCBI GEO database under accession number []. All other data are present in the article and Supplementary Information.

## REFERENCES

Andreatta, M., A. Tjitropranoto, Z. Sherman, M.C. Kelly, T. Ciucci, and S.J. Carmona. 2022. A CD4(+) T cell reference map delineates subtype-specific adaptation during acute and chronic viral infections. eLife 11:

Ashhurst, A.S., M. Flórido, L.C.W. Lin, D. Quan, E. Armitage, S.A. Stifter, J. Stambas, and W.J. Britton. 2019. CXCR6-Deficiency Improves the Control of Pulmonary Mycobacterium tuberculosis and Influenza Infection Independent of T-Lymphocyte Recruitment to the Lungs. Frontiers in immunology 10:339.

Bates, J.T. 2022. Naïve CD4(+) T Cell Activation in the Nasal-Associated Lymphoid Tissue following Intranasal Immunization with a Flagellin-Based Subunit Vaccine. International journal of molecular sciences 23:

Becht, E., L. McInnes, J. Healy, C.A. Dutertre, I.W.H. Kwok, L.G. Ng, F. Ginhoux, and E.W. Newell. 2018. Dimensionality reduction for visualizing single-cell data using UMAP. Nature biotechnology

Bergsbaken, T., and M.J. Bevan. 2015. Proinflammatory microenvironments within the intestine regulate the differentiation of tissue-resident CD8⁺ T cells responding to infection. Nature immunology 16:406–414.

Besavilla, D.F., L. Reusch, J. Enriquez, K. Schon, and D. Angeletti. 2023. Pre-existing CD4 T cell help boosts antibody responses but has limited impact on germinal center, antigen-specific B cell frequencies after influenza infection. Front Immunol 14:1243164.

Beura, L.K., N.J. Fares-Frederickson, E.M. Steinert, M.C. Scott, E.A. Thompson, K.A. Fraser, J.M. Schenkel, V. Vezys, and D. Masopust. 2019. CD4(+) resident memory T cells dominate immunosurveillance and orchestrate local recall responses. The Journal of experimental medicine 216:1214–1229.

Blighe K, R.S., Lewis M 2024. EnhancedVolcano: Publication-ready volcano plots with enhanced colouring and labeling, R package version 1.22.0,https://github.com/kevinblighe/EnhancedVolcano.

Bodewes, R., G. de Mutsert, F.R. van der Klis, M. Ventresca, S. Wilks, D.J. Smith, M. Koopmans, R.A. Fouchier, A.D. Osterhaus, and G.F. Rimmelzwaan. 2011. Prevalence of antibodies against seasonal influenza A and B viruses in children in Netherlands. Clin Vaccine Immunol 18:469–476.

Borcherding, N., N.L. Bormann, and G. Kraus. 2020. scRepertoire: An R-based toolkit for single-cell immune receptor analysis. F1000Research 9:47.

Borkner, L., L.M. Curham, M.M. Wilk, B. Moran, and K.H.G. Mills. 2021. IL-17 mediates protective immunity against nasal infection with Bordetella pertussis by mobilizing neutrophils, especially Siglec-F(+) neutrophils. Mucosal immunology 14:1183–1202.

Brass, A.L., I.C. Huang, Y. Benita, S.P. John, M.N. Krishnan, E.M. Feeley, B.J. Ryan, J.L. Weyer, L. van der Weyden, E. Fikrig, D.J. Adams, R.J. Xavier, M. Farzan, and S.J. Elledge. 2009. The IFITM proteins mediate cellular resistance to influenza A H1N1 virus, West Nile virus, and dengue virus. Cell 139:1243–1254.

Brinkmann, V., M.D. Davis, C.E. Heise, R. Albert, S. Cottens, R. Hof, C. Bruns, E. Prieschl, T. Baumruker, P. Hiestand, C.A. Foster, M. Zollinger, and K.R. Lynch. 2002. The immune modulator FTY720 targets sphingosine 1-phosphate receptors. The Journal of biological chemistry 277:21453–21457.

Brockmann, L., A. Tran, Y. Huang, M. Edwards, C. Ronda, H.H. Wang, and Ivanov, II. 2023. Intestinal microbiota-specific Th17 cells possess regulatory properties and suppress effector T cells via c-MAF and IL-10. Immunity 56:2719–2735.e2717.

Brown, D.M., S. Lee, L. Garcia-Hernandez Mde, and S.L. Swain. 2012. Multifunctional CD4 cells expressing gamma interferon and perforin mediate protection against lethal influenza virus infection. Journal of virology 86:6792–6803.

Brownlie, D., I. Rødahl, R. Varnaite, H. Asgeirsson, H. Glans, S. Falck-Jones, S. Vangeti, M. Buggert, H.G. Ljunggren, J. Michaëlsson, S. Gredmark-Russ, A. Smed-Sörensen, and N. Marquardt. 2022. Comparison of Lung-Homing Receptor Expression and Activation Profiles on NK Cell and T Cell Subsets in COVID-19 and Influenza. Frontiers in immunology 13:834862.

Caminschi, I., M.H. Lahoud, A. Pizzolla, and L.M. Wakim. 2019. Zymosan by-passes the requirement for pulmonary antigen encounter in lung tissue-resident memory CD8(+) T cell development. Mucosal immunology 12:403–412.

Campbell, A.P., C. Ogokeh, J.Y. Lively, M.A. Staat, R. Selvarangan, N.B. Halasa, J.A. Englund, J.A. Boom, G.A. Weinberg, J.V. Williams, M. McNeal, C.J. Harrison, L.S. Stewart, E.J. Klein, L.C. Sahni, P.G. Szilagyi, M.G. Michaels, R.W. Hickey, M.E. Moffat, B.A. Pahud, J.E. Schuster, G.M. Weddle, B. Rha, A.M. Fry, and M. Patel. 2020. Vaccine Effectiveness Against Pediatric Influenza Hospitalizations and Emergency Visits. Pediatrics 146:

Cohen, N.R., P.J. Brennan, T. Shay, G.F. Watts, M. Brigl, J. Kang, and M.B. Brenner. 2013. Shared and distinct transcriptional programs underlie the hybrid nature of iNKT cells. Nature immunology 14:90–99.

Devaiah, B.N., H. Lu, A. Gegonne, Z. Sercan, H. Zhang, R.J. Clifford, M.P. Lee, and D.S. Singer. 2010. Novel functions for TAF7, a regulator of TAF1-independent transcription. The Journal of biological chemistry 285:38772–38780.

Dey, S., H. Ashwin, L. Milross, B. Hunter, J. Majo, A.J. Filby, A.J. Fisher, P.M. Kaye, and D. Lagos. 2023. Downregulation of MALAT1 is a hallmark of tissue and peripheral proliferative T cells in COVID-19. Clinical and experimental immunology 212:262–275.

Di Pilato, M., R. Kfuri-Rubens, J.N. Pruessmann, A.J. Ozga, M. Messemaker, B.L. Cadilha, R. Sivakumar, C. Cianciaruso, R.D. Warner, F. Marangoni, E. Carrizosa, S. Lesch, J. Billingsley, D. Perez-Ramos, F. Zavala, E. Rheinbay, A.D. Luster, M.Y. Gerner, S. Kobold, M.J. Pittet, and T.R. Mempel. 2021. CXCR6 positions cytotoxic T cells to receive critical survival signals in the tumor microenvironment. Cell 184:4512–4530.e4522.

Diallo, B.K., N.C. C, T.Y. Wong, P. Schmitt, K.S. Lee, K. Weaver, O. Miller, M. Cooper, S.D. Jazayeri, F.H. Damron, and K.H.G. Mills. 2023a. Intranasal COVID-19 vaccine induces respiratory memory T cells and protects K18-hACE mice against SARS-CoV-2 infection. NPJ vaccines 8:68.

Diallo, B.K., C. Ní Chasaide, T.Y. Wong, P. Schmitt, K.S. Lee, K. Weaver, O. Miller, M. Cooper, S.D. Jazayeri, F.H. Damron, and K.H.G. Mills. 2023b. Intranasal COVID-19 vaccine induces respiratory memory T cells and protects K18-hACE mice against SARS-CoV-2 infection. NPJ vaccines 8:68.

Dutta, A., C.Y. Hung, T.C. Chen, S.H. Hsiao, C.S. Chang, Y.C. Lin, C.Y. Lin, and C.T. Huang. 2023. An IL-17-EGFR-TRAF4 axis contributes to the alleviation of lung inflammation in severe influenza. Communications biology 6:600.

Eliasson, D.G., K. El Bakkouri, K. Schön, A. Ramne, E. Festjens, B. Löwenadler, W. Fiers, X. Saelens, and N. Lycke. 2008. CTA1-M2e-DD: a novel mucosal adjuvant targeted influenza vaccine. Vaccine 26:1243–1252.

Eliasson, D.G., A. Omokanye, K. Schon, U.A. Wenzel, V. Bernasconi, M. Bemark, A. Kolpe, K. El Bakkouri, T. Ysenbaert, L. Deng, W. Fiers, X. Saelens, and N. Lycke. 2018. M2e-tetramer-specific memory CD4 T cells are broadly protective against influenza infection. Mucosal immunology 11:273–289.

Feeley, E.M., J.S. Sims, S.P. John, C.R. Chin, T. Pertel, L.M. Chen, G.D. Gaiha, B.J. Ryan, R.O. Donis, S.J. Elledge, and A.L. Brass. 2011. IFITM3 inhibits influenza A virus infection by preventing cytosolic entry. PLoS pathogens 7:e1002337.

Fong, C.H., L. Lu, L.L. Chen, M.L. Yeung, A.J. Zhang, H. Zhao, K.Y. Yuen, and K.K. To. 2022. Interferon-gamma inhibits influenza A virus cellular attachment by reducing sialic acid cluster size. iScience 25:104037.

Gailleton, R., N.R. Mathew, L. Reusch, K. Schon, L. Scharf, A. Stromberg, A. Cvjetkovic, L. Aziz, J. Hellgren, K.W. Tang, M. Bemark, and D. Angeletti. 2025. Ectopic germinal centers in the nasal turbinates contribute to B cell immunity to intranasal viral infection and vaccination. Proc Natl Acad Sci U S A 122:e2421724122.

Goplen, N.P., Y. Wu, Y.M. Son, C. Li, Z. Wang, I.S. Cheon, L. Jiang, B. Zhu, K. Ayasoufi, E.N. Chini, A.J. Johnson, R. Vassallo, A.H. Limper, N. Zhang, and J. Sun. 2020. Tissue-resident CD8(+) T cells drive age-associated chronic lung sequelae after viral pneumonia. Science immunology 5:

Gribonika, I., A. Stromberg, C. Lebrero-Fernandez, K. Schon, J. Moon, M. Bemark, and N. Lycke. 2022. Peyer’s patch T(H)17 cells are dispensable for gut IgA responses to oral immunization. Sci Immunol 7:eabc5500.

Hao, Y., S. Hao, E. Andersen-Nissen, W.M. Mauck, 3rd, S. Zheng, A. Butler, M.J. Lee, A.J. Wilk, C. Darby, M. Zager, P. Hoffman, M. Stoeckius, E. Papalexi, E.P. Mimitou, J. Jain, A. Srivastava, T. Stuart, L.M. Fleming, B. Yeung, A.J. Rogers, J.M. McElrath, C.A. Blish, R. Gottardo, P. Smibert, and R. Satija. 2021. Integrated analysis of multimodal single-cell data. Cell 184:3573–3587.e3529.

Hao, Y., T. Stuart, M.H. Kowalski, S. Choudhary, P. Hoffman, A. Hartman, A. Srivastava, G. Molla, S. Madad, C. Fernandez-Granda, and R. Satija. 2024. Dictionary learning for integrative, multimodal and scalable single-cell analysis. Nature biotechnology 42:293–304.

Hashimoto, K., T. Kouno, T. Ikawa, N. Hayatsu, Y. Miyajima, H. Yabukami, T. Terooatea, T. Sasaki, T. Suzuki, M. Valentine, G. Pascarella, Y. Okazaki, H. Suzuki, J.W. Shin, A. Minoda, I. Taniuchi, H. Okano, Y. Arai, N. Hirose, and P. Carninci. 2019. Single-cell transcriptomics reveals expansion of cytotoxic CD4 T cells in supercentenarians. Proc Natl Acad Sci U S A 116:24242–24251.

Hoft, S.G., M.A. Sallin, K.D. Kauffman, S. Sakai, V.V. Ganusov, and D.L. Barber. 2019. The Rate of CD4 T Cell Entry into the Lungs during Mycobacterium tuberculosis Infection Is Determined by Partial and Opposing Effects of Multiple Chemokine Receptors. Infection and immunity 87:

Hu, X., M. Wu, T. Ma, Y. Zhang, C. Zou, R. Wang, Y. Zhang, Y. Ren, Q. Li, H. Liu, H. Li, T. Wang, X. Sun, Y. Yang, M. Tang, X. Li, J. Li, X. Gao, T. Li, and X. Zhou. 2022. Single-cell transcriptomics reveals distinct cell response between acute and chronic pulmonary infection of Pseudomonas aeruginosa. MedComm 3:e193.

Ivanov, II, B.S. McKenzie, L. Zhou, C.E. Tadokoro, A. Lepelley, J.J. Lafaille, D.J. Cua, and D.R. Littman. 2006. The orphan nuclear receptor RORgammat directs the differentiation program of proinflammatory IL-17+ T helper cells. Cell 126:1121–1133.

Jaffar, Z., M.E. Ferrini, L.A. Herritt, and K. Roberts. 2009. Cutting edge: lung mucosal Th17-mediated responses induce polymeric Ig receptor expression by the airway epithelium and elevate secretory IgA levels. Journal of immunology (Baltimore, Md. : 1950) 182:4507–4511.

Kazer, S.W., C.M. Match, E.M. Langan, M.-A. Messou, T.J. LaSalle, E. O’Leary, J. Marbourg, K. Naughton, U.H. von Andrian, and J. Ordovas-Montanes. 2024. Primary nasal influenza infection rewires the tissue-scale memory response dynamics. Immunity

Khan, T.N., J.L. Mooster, A.M. Kilgore, J.F. Osborn, and J.C. Nolz. 2016. Local antigen in nonlymphoid tissue promotes resident memory CD8+ T cell formation during viral infection. The Journal of experimental medicine 213:951–966.

Kinnear, E., L. Lambert, J.U. McDonald, H.M. Cheeseman, L.J. Caproni, and J.S. Tregoning. 2018. Airway T cells protect against RSV infection in the absence of antibody. Mucosal immunology 11:249–256.

Koenen, A., A. Babendreyer, J. Schumacher, T. Pasqualon, N. Schwarz, A. Seifert, X. Deupi, A. Ludwig, and D. Dreymueller. 2017. The DRF motif of CXCR6 as chemokine receptor adaptation to adhesion. PloS one 12:e0173486.

Konieczny, P., Y. Xing, I. Sidhu, I. Subudhi, K.P. Mansfield, B. Hsieh, D.E. Biancur, S.B. Larsen, M. Cammer, D. Li, N.X. Landén, C. Loomis, A. Heguy, A.N. Tikhonova, A. Tsirigos, and S. Naik. 2022. Interleukin-17 governs hypoxic adaptation of injured epithelium. Science (New York, N.Y.) 377:eabg9302.

Kourepini, E., M. Aggelakopoulou, T. Alissafi, N. Paschalidis, D.C. Simoes, and V. Panoutsakopoulou. 2014. Osteopontin expression by CD103-dendritic cells drives intestinal inflammation. Proc Natl Acad Sci U S A 111:E856–865.

Künzli, M., and D. Masopust. 2023. CD4(+) T cell memory. Nature immunology 24:903–914.

Kurd, N.S., Z. He, T.L. Louis, J.J. Milner, K.D. Omilusik, W. Jin, M.S. Tsai, C.E. Widjaja, J.N. Kanbar, J.G. Olvera, T. Tysl, L.K. Quezada, B.S. Boland, W.J. Huang, C. Murre, A.W. Goldrath, G.W. Yeo, and J.T. Chang. 2020. Early precursors and molecular determinants of tissue-resident memory CD8(+) T lymphocytes revealed by single-cell RNA sequencing. Science immunology 5:

Laan, M., Z.H. Cui, H. Hoshino, J. Lötvall, M. Sjöstrand, D.C. Gruenert, B.E. Skoogh, and A. Lindén. 1999. Neutrophil recruitment by human IL-17 via C-X-C chemokine release in the airways. Journal of immunology (Baltimore, Md. : 1950) 162:2347–2352.

Li, J., E. Lu, T. Yi, and J.G. Cyster. 2016. EBI2 augments Tfh cell fate by promoting interaction with IL-2-quenching dendritic cells. Nature 533:110–114.

Liang, S.C., X.Y. Tan, D.P. Luxenberg, R. Karim, K. Dunussi-Joannopoulos, M. Collins, and L.A. Fouser. 2006. Interleukin (IL)-22 and IL-17 are coexpressed by Th17 cells and cooperatively enhance expression of antimicrobial peptides. The Journal of experimental medicine 203:2271–2279.

Lim, J.M.E., A.T. Tan, N. Le Bert, S.K. Hang, J.G.H. Low, and A. Bertoletti. 2022. SARS-CoV-2 breakthrough infection in vaccinees induces virus-specific nasal-resident CD8+ and CD4+ T cells of broad specificity. J Exp Med 219:

Lin, Y., S. Ritchea, A. Logar, S. Slight, M. Messmer, J. Rangel-Moreno, L. Guglani, J.F. Alcorn, H. Strawbridge, S.M. Park, R. Onishi, N. Nyugen, M.J. Walter, D. Pociask, T.D. Randall, S.L. Gaffen, Y. Iwakura, J.K. Kolls, and S.A. Khader. 2009. Interleukin-17 is required for T helper 1 cell immunity and host resistance to the intracellular pathogen Francisella tularensis. Immunity 31:799–810.

Liu, J., L. Stoler-Barak, H. Hezroni-Bravyi, A. Biram, S. Lebon, N. Davidzohn, M. Kedmi, M. Chemla, D. Pilzer, M. Cohen, O. Brenner, M. Biton, and Z. Shulman. 2024. Turbinate-homing IgA-secreting cells originate in the nasal lymphoid tissues. Nature 632:637–646.

Mabrouk, N., T. Tran, I. Sam, I. Pourmir, N. Gruel, C. Granier, J. Pineau, A. Gey, S. Kobold, E. Fabre, and E. Tartour. 2022. CXCR6 expressing T cells: Functions and role in the control of tumors. Frontiers in immunology 13:1022136.

Marchesini Tovar, G., C. Gallen, and T. Bergsbaken. 2024. CD8+ Tissue-Resident Memory T Cells: Versatile Guardians of the Tissue. Journal of immunology (Baltimore, Md. : 1950) 212:361–368.

Maroof, A., Y.M. Yorgensen, Y. Li, and J.T. Evans. 2014. Intranasal vaccination promotes detrimental Th17-mediated immunity against influenza infection. PLoS pathogens 10:e1003875.

Mas-Orea, X., L. Rey, L. Battut, C. Bories, C. Petitfils, A. Abot, N. Gheziel, E. Wemelle, C. Blanpied, J.P. Motta, C. Knauf, F. Barreau, E. Espinosa, M. Aloulou, N. Cenac, M. Serino, L. Mouledous, N. Fazilleau, and G. Dietrich. 2023. Proenkephalin deletion in hematopoietic cells induces intestinal barrier failure resulting in clinical feature similarities with irritable bowel syndrome in mice. Communications biology 6:1168.

McCarthy, K.N., S. Hone, R.M. McLoughlin, and K.H.G. Mills. 2024. IL-17 and IFN-γ-producing respiratory tissue resident memory CD4 T cells persist for decades in adults immunized as children with whole cell pertussis vaccines. The Journal of infectious diseases

McGeachy, M.J. 2011. GM-CSF: the secret weapon in the T(H)17 arsenal. Nature immunology 12:521–522.

McKinstry, K.K., T.M. Strutt, Y. Kuang, D.M. Brown, S. Sell, R.W. Dutton, and S.L. Swain. 2012. Memory CD4+ T cells protect against influenza through multiple synergizing mechanisms. The Journal of clinical investigation 122:2847–2856.

McMaster, S.R., A.N. Wein, P.R. Dunbar, S.L. Hayward, E.K. Cartwright, T.L. Denning, and J.E. Kohlmeier. 2018. Pulmonary antigen encounter regulates the establishment of tissue-resident CD8 memory T cells in the lung airways and parenchyma. Mucosal immunology 11:1071–1078.

Miller, I., M. Min, C. Yang, C. Tian, S. Gookin, D. Carter, and S.L. Spencer. 2018. Ki67 is a Graded Rather than a Binary Marker of Proliferation versus Quiescence. Cell Rep 24:1105–1112.e1105.

Miller, M.A., A.P. Ganesan, N. Luckashenak, M. Mendonca, and L.C. Eisenlohr. 2015. Endogenous antigen processing drives the primary CD4+ T cell response to influenza. Nature medicine 21:1216–1222.

Mills, K.H.G. 2023. IL-17 and IL-17-producing cells in protection versus pathology. Nature reviews. Immunology 23:38–54.

Moguche, A.O., S. Shafiani, C. Clemons, R.P. Larson, C. Dinh, L.E. Higdon, C.J. Cambier, J.R. Sissons, A.M. Gallegos, P.J. Fink, and K.B. Urdahl. 2015. ICOS and Bcl6-dependent pathways maintain a CD4 T cell population with memory-like properties during tuberculosis. The Journal of experimental medicine 212:715–728.

Ng, S.S., F. De Labastida Rivera, J. Yan, D. Corvino, I. Das, P. Zhang, R. Kuns, S.B. Chauhan, J. Hou, X.Y. Li, T.C.M. Frame, B.A. McEnroe, E. Moore, J. Na, J.A. Engel, M.S.F. Soon, B. Singh, A.J. Kueh, M.J. Herold, M. Montes de Oca, S.S. Singh, P.T. Bunn, A.R. Aguilera, M. Casey, M. Braun, N. Ghazanfari, S. Wani, Y. Wang, F.H. Amante, C.L. Edwards, A. Haque, W.C. Dougall, O.P. Singh, A.G. Baxter, M.W.L. Teng, A. Loukas, N.L. Daly, N. Cloonan, M.A. Degli-Esposti, J. Uzonna, W.R. Heath, T. Bald, S.K. Tey, K. Nakamura, G.R. Hill, R. Kumar, S. Sundar, M.J. Smyth, and C.R. Engwerda. 2024. Author Correction: The NK cell granule protein NKG7 regulates cytotoxic granule exocytosis and inflammation. Nature immunology 25:716.

Nguyen, K.D., A. Fohner, J.D. Booker, C. Dong, A.M. Krensky, and K.C. Nadeau. 2008. XCL1 enhances regulatory activities of CD4+ CD25(high) CD127(low/-) T cells in human allergic asthma. Journal of immunology (Baltimore, Md. : 1950) 181:5386–5395.

O’Hara, J.M., N.S. Redhu, E. Cheung, N.G. Robertson, I. Patik, S.E. Sayed, C.M. Thompson, M. Herd, K.B. Lucas, E. Conaway, C.C. Morton, D.L. Farber, R. Malley, and B.H. Horwitz. 2020. Generation of protective pneumococcal-specific nasal resident memory CD4(+) T cells via parenteral immunization. Mucosal immunology 13:172–182.

Omokanye, A., L.C. Ong, C. Lebrero-Fernandez, V. Bernasconi, K. Schon, A. Stromberg, M. Bemark, X. Saelens, P. Czarnewski, and N. Lycke. 2022. Clonotypic analysis of protective influenza M2e-specific lung resident Th17 memory cells reveals extensive functional diversity. Mucosal immunology 15:717–729.

Paget, J., P. Spreeuwenberg, V. Charu, R.J. Taylor, A.D. Iuliano, J. Bresee, L. Simonsen, and C. Viboud. 2019. Global mortality associated with seasonal influenza epidemics: New burden estimates and predictors from the GLaMOR Project. Journal of global health 9:020421.

Pandiyan, P., N. Bhaskaran, M. Zou, E. Schneider, S. Jayaraman, and J. Huehn. 2019. Microbiome Dependent Regulation of T(regs) and Th17 Cells in Mucosa. Frontiers in immunology 10:426.

Peng, C., M.A. Huggins, K.M. Wanhainen, T.P. Knutson, H. Lu, H. Georgiev, K.L. Mittelsteadt, N.N. Jarjour, H. Wang, K.A. Hogquist, D.J. Campbell, H. Borges da Silva, and S.C. Jameson. 2022. Engagement of the costimulatory molecule ICOS in tissues promotes establishment of CD8(+) tissue-resident memory T cells. Immunity 55:98–114.e115.

Pizzolla, A., T.H.O. Nguyen, J.M. Smith, A.G. Brooks, K. Kedzieska, W.R. Heath, P.C. Reading, and L.M. Wakim. 2017a. Resident memory CD8(+) T cells in the upper respiratory tract prevent pulmonary influenza virus infection. Sci Immunol 2:

Pizzolla, A., Z. Wang, J.R. Groom, K. Kedzierska, A.G. Brooks, P.C. Reading, and L.M. Wakim. 2017b. Nasal-associated lymphoid tissues (NALTs) support the recall but not priming of influenza virus-specific cytotoxic T cells. Proc Natl Acad Sci U S A 114:5225–5230.

Plasek, L.M., and S. Valadkhan. 2021. lncRNAs in T lymphocytes: RNA regulation at the heart of the immune response. American journal of physiology. Cell physiology 320:C415–c427.

Ramirez, S.I., F. Faraji, L.B. Hills, P.G. Lopez, B. Goodwin, H.D. Stacey, H.J. Sutton, K.M. Hastie, E.O. Saphire, H.J. Kim, S. Mashoof, C.H. Yan, A.S. DeConde, G. Levi, and S. Crotty. 2024. Immunological memory diversity in the human upper airway. Nature 632:630–636.

Randall, T.D. Structure, Organization, and Development of the Mucosal Immune System of the Respiratory Tract. Mucosal Immunology. 2015:43–61. doi: 10.1016/B978-0-12-415847-4.00004-5. Epub 2015 Mar 13.,

Rauen, T., C.M. Hedrich, K. Tenbrock, and G.C. Tsokos. 2013. cAMP responsive element modulator: a critical regulator of cytokine production. Trends in molecular medicine 19:262–269.

Richard, M., J.M.A. van den Brand, T.M. Bestebroer, P. Lexmond, D. de Meulder, R.A.M. Fouchier, A.C. Lowen, and S. Herfst. 2020. Influenza A viruses are transmitted via the air from the nasal respiratory epithelium of ferrets. Nature communications 11:766.

Satija, R., J.A. Farrell, D. Gennert, A.F. Schier, and A. Regev. 2015. Spatial reconstruction of single-cell gene expression data. Nature biotechnology 33:495–502.

Sato, A., A. Suwanto, M. Okabe, S. Sato, T. Nochi, T. Imai, N. Koyanagi, J. Kunisawa, Y. Kawaguchi, and H. Kiyono. 2014. Vaginal memory T cells induced by intranasal vaccination are critical for protective T cell recruitment and prevention of genital HSV-2 disease. Journal of virology 88:13699–13708.

Sekine, T., A. Perez-Potti, O. Rivera-Ballesteros, K. Strålin, J.B. Gorin, A. Olsson, S. Llewellyn-Lacey, H. Kamal, G. Bogdanovic, S. Muschiol, D.J. Wullimann, T. Kammann, J. Emgård, T. Parrot, E. Folkesson, O. Rooyackers, L.I. Eriksson, J.I. Henter, A. Sönnerborg, T. Allander, J. Albert, M. Nielsen, J. Klingström, S. Gredmark-Russ, N.K. Björkström, J.K. Sandberg, D.A. Price, H.G. Ljunggren, S. Aleman, and M. Buggert. 2020. Robust T Cell Immunity in Convalescent Individuals with Asymptomatic or Mild COVID-19. Cell 183:158–168.e114.

Shiow, L.R., D.B. Rosen, N. Brdicková, Y. Xu, J. An, L.L. Lanier, J.G. Cyster, and M. Matloubian. 2006. CD69 acts downstream of interferon-alpha/beta to inhibit S1P1 and lymphocyte egress from lymphoid organs. Nature 440:540–544.

Solans, L., A.S. Debrie, L. Borkner, N. Aguiló, A. Thiriard, L. Coutte, S. Uranga, F. Trottein, C. Martín, K.H.G. Mills, and C. Locht. 2018. IL-17-dependent SIgA-mediated protection against nasal Bordetella pertussis infection by live attenuated BPZE1 vaccine. Mucosal immunology 11:1753–1762.

Son, Y.M., I.S. Cheon, Y. Wu, C. Li, Z. Wang, X. Gao, Y. Chen, Y. Takahashi, Y.X. Fu, A.L. Dent, M.H. Kaplan, J.J. Taylor, W. Cui, and J. Sun. 2021. Tissue-resident CD4(+) T helper cells assist the development of protective respiratory B and CD8(+) T cell memory responses. Science immunology 6:

Sonnenberg, G.F., M.G. Nair, T.J. Kirn, C. Zaph, L.A. Fouser, and D. Artis. 2010. Pathological versus protective functions of IL-22 in airway inflammation are regulated by IL-17A. The Journal of experimental medicine 207:1293–1305.

Swarnalekha, N., D. Schreiner, L.C. Litzler, S. Iftikhar, D. Kirchmeier, M. Kunzli, Y.M. Son, J. Sun, E.A. Moreira, and C.G. King. 2021. T resident helper cells promote humoral responses in the lung. Sci Immunol 6:

Szabo, P.A., H.M. Levitin, M. Miron, M.E. Snyder, T. Senda, J. Yuan, Y.L. Cheng, E.C. Bush, P. Dogra, P. Thapa, D.L. Farber, and P.A. Sims. 2019a. Single-cell transcriptomics of human T cells reveals tissue and activation signatures in health and disease. Nature communications 10:4706.

Szabo, P.A., M. Miron, and D.L. Farber. 2019b. Location, location, location: Tissue resident memory T cells in mice and humans. Science immunology 4:

Takamura, S., H. Yagi, Y. Hakata, C. Motozono, S.R. McMaster, T. Masumoto, M. Fujisawa, T. Chikaishi, J. Komeda, J. Itoh, M. Umemura, A. Kyusai, M. Tomura, T. Nakayama, D.L. Woodland, J.E. Kohlmeier, and M. Miyazawa. 2016. Specific niches for lung-resident memory CD8+ T cells at the site of tissue regeneration enable CD69-independent maintenance. The Journal of experimental medicine 213:3057–3073.

Teijaro, J.R., D. Turner, Q. Pham, E.J. Wherry, L. Lefrancois, and D.L. Farber. 2011. Cutting edge: Tissue-retentive lung memory CD4 T cells mediate optimal protection to respiratory virus infection. Journal of immunology (Baltimore, Md. : 1950) 187:5510–5514.

The, L. 2024. H5N1: international failures and uncomfortable truths. *Lancet (London*, England) 403:2455.

Thomas, P.G., S.A. Brown, W. Yue, J. So, R.J. Webby, and P.C. Doherty. 2006. An unexpected antibody response to an engineered influenza virus modifies CD8+ T cell responses. Proc Natl Acad Sci U S A 103:2764–2769.

Thome, J.J., N. Yudanin, Y. Ohmura, M. Kubota, B. Grinshpun, T. Sathaliyawala, T. Kato, H. Lerner, Y. Shen, and D.L. Farber. 2014. Spatial map of human T cell compartmentalization and maintenance over decades of life. Cell 159:814–828.

Tuomela, K., A.R. Ambrose, and D.M. Davis. 2022. Escaping Death: How Cancer Cells and Infected Cells Resist Cell-Mediated Cytotoxicity. Frontiers in immunology 13:867098.

Turner, D.L., K.L. Bickham, J.J. Thome, C.Y. Kim, F. D’Ovidio, E.J. Wherry, and D.L. Farber. 2014. Lung niches for the generation and maintenance of tissue-resident memory T cells. Mucosal immunology 7:501–510.

Uddbäck, I., S.E. Michalets, A. Saha, C. Mattingly, K.N. Kost, M.E. Williams, L.A. Lawrence, S.L. Hicks, A.C. Lowen, H. Ahmed, A.R. Thomsen, C.J. Russell, C.D. Scharer, J.M. Boss, K. Koelle, R. Antia, J.P. Christensen, and J.E. Kohlmeier. 2024. Prevention of respiratory virus transmission by resident memory CD8(+) T cells. Nature 626:392–400.

Vijayanand, S., K.B. Gomes, R.P. Gala, M.N. Uddin, and M.J. D’Souza. 2020. Exploring the Potential of T-Cells for a Universal Influenza Vaccine. Vaccines 8:

Wakim, L.M., A. Woodward-Davis, and M.J. Bevan. 2010. Memory T cells persisting within the brain after local infection show functional adaptations to their tissue of residence. Proc Natl Acad Sci U S A 107:17872–17879.

Wang, X., C.C. Chan, M. Yang, J. Deng, V.K. Poon, V.H. Leung, K.H. Ko, J. Zhou, K.Y. Yuen, B.J. Zheng, and L. Lu. 2011. A critical role of IL-17 in modulating the B-cell response during H5N1 influenza virus infection. Cellular & molecular immunology 8:462–468.

Wang, X., X. Shen, S. Chen, H. Liu, N. Hong, H. Zhong, X. Chen, and W. Jin. 2022. Reinvestigation of Classic T Cell Subsets and Identification of Novel Cell Subpopulations by Single-Cell RNA Sequencing. Journal of immunology (Baltimore, Md. : 1950) 208:396–406.

Wang, Z., S. Wang, N.P. Goplen, C. Li, I.S. Cheon, Q. Dai, S. Huang, J. Shan, C. Ma, Z. Ye, M. Xiang, A.H. Limper, E.C. Porquera, J.E. Kohlmeier, M.H. Kaplan, N. Zhang, A.J. Johnson, R. Vassallo, and J. Sun. 2019. PD-1(hi) CD8(+) resident memory T cells balance immunity and fibrotic sequelae. Science immunology 4:

Wein, A.N., S.R. McMaster, S. Takamura, P.R. Dunbar, E.K. Cartwright, S.L. Hayward, D.T. McManus, T. Shimaoka, S. Ueha, T. Tsukui, T. Masumoto, M. Kurachi, K. Matsushima, and J.E. Kohlmeier. 2019. CXCR6 regulates localization of tissue-resident memory CD8 T cells to the airways. The Journal of experimental medicine 216:2748–2762.

Weiss, I.D., O. Wald, H. Wald, K. Beider, M. Abraham, E. Galun, A. Nagler, and A. Peled. 2010. IFN-gamma treatment at early stages of influenza virus infection protects mice from death in a NK cell-dependent manner. Journal of interferon & cytokine research : the official journal of the International Society for Interferon and Cytokine Research 30:439–449.

Wilbanks, A., S.C. Zondlo, K. Murphy, S. Mak, D. Soler, P. Langdon, D.P. Andrew, L. Wu, and M. Briskin. 2001. Expression cloning of the STRL33/BONZO/TYMSTRligand reveals elements of CC, CXC, and CX3C chemokines. Journal of immunology (Baltimore, Md. : 1950) 166:5145–5154.

Wimmers, F., A.R. Burrell, Y. Feng, H. Zheng, P.S. Arunachalam, M. Hu, S. Spranger, L.E. Nyhoff, D. Joshi, M. Trisal, M. Awasthi, L. Bellusci, U. Ashraf, S. Kowli, K.C. Konvinse, E. Yang, M. Blanco, K. Pellegrini, G. Tharp, T. Hagan, R.S. Chinthrajah, T.T. Nguyen, A. Grifoni, A. Sette, K.C. Nadeau, D.B. Haslam, S.E. Bosinger, J. Wrammert, H.T. Maecker, P.J. Utz, T.T. Wang, S. Khurana, P. Khatri, M.A. Staat, and B. Pulendran. 2023. Multi-omics analysis of mucosal and systemic immunity to SARS-CoV-2 after birth. Cell 186:4632–4651.e4623.

Woodring, T., C.N. Dewey, L.D. Santos Dias, X. He, H.E. Dobson, M. Wüthrich, and B. Klein. 2022. Distinctive populations of CD4(+)T cells associated with vaccine efficacy. iScience 25:104934.

Xu, Q., P. Milanez-Almeida, A.J. Martins, A.J. Radtke, K.B. Hoehn, C. Oguz, J. Chen, C. Liu, J. Tang, G. Grubbs, S. Stein, S. Ramelli, J. Kabat, H. Behzadpour, M. Karkanitsa, J. Spathies, H. Kalish, L. Kardava, M. Kirby, F. Cheung, S. Preite, P.C. Duncker, M.M. Kitakule, N. Romero, D. Preciado, L. Gitman, G. Koroleva, G. Smith, A. Shaffer, I.T. McBain, P.J. McGuire, S. Pittaluga, R.N. Germain, R. Apps, D.M. Schwartz, K. Sadtler, S. Moir, D.S. Chertow, S.H. Kleinstein, S. Khurana, J.S. Tsang, P. Mudd, P.L. Schwartzberg, and K. Manthiram. 2023. Adaptive immune responses to SARS-CoV-2 persist in the pharyngeal lymphoid tissue of children. Nature immunology 24:186–199.

Yan, Y., G.X. Zhang, M.S. Williams, G.B. Carey, H. Li, J. Yang, A. Rostami, and H. Xu. 2012. TCR stimulation upregulates MS4a4B expression through induction of AP-1 transcription factor during T cell activation. Molecular immunology 52:71–78.

Yang, H.Q., Y.S. Wang, K. Zhai, and Z.H. Tong. 2021. Single-Cell TCR Sequencing Reveals the Dynamics of T Cell Repertoire Profiling During Pneumocystis Infection. Frontiers in microbiology 12:637500.

Yount, K.S., J.M. Hall, K. Caution, M.M. Shamseldin, M. Guo, K. Marion, A.R. Fullen, Y. Huang, J.A. Maynard, S.A. Quataert, R. Deora, and P. Dubey. 2023. Systemic priming and intranasal booster with a BcfA-adjuvanted acellular pertussis vaccine generates CD4+ IL-17+ nasal tissue resident T cells and reduces B. pertussis nasal colonization. Frontiers in immunology 14:1181876.

Zens, K.D., J.K. Chen, and D.L. Farber. 2016. Vaccine-generated lung tissue-resident memory T cells provide heterosubtypic protection to influenza infection. JCI insight 1:

Zens, K.D., J.K. Chen, R.S. Guyer, F.L. Wu, F. Cvetkovski, M. Miron, and D.L. Farber. 2017. Reduced generation of lung tissue-resident memory T cells during infancy. The Journal of experimental medicine 214:2915–2932.

Zheng, M.Z.M., T.K. Tan, F. Villalon-Letelier, H. Lau, Y.M. Deng, S. Fritzlar, S.A. Valkenburg, H. Gu, L.L.M. Poon, P.C. Reading, A.R. Townsend, and L.M. Wakim. 2023. Single-cycle influenza virus vaccine generates lung CD8(+) Trm that cross-react against viral variants and subvert virus escape mutants. Science advances 9:eadg3469.

