## Supplementary Figures for "Nasal tissue-resident memory CD4^+^ T cells persist after influenza A virus infection and provide heterosubtypic protection"

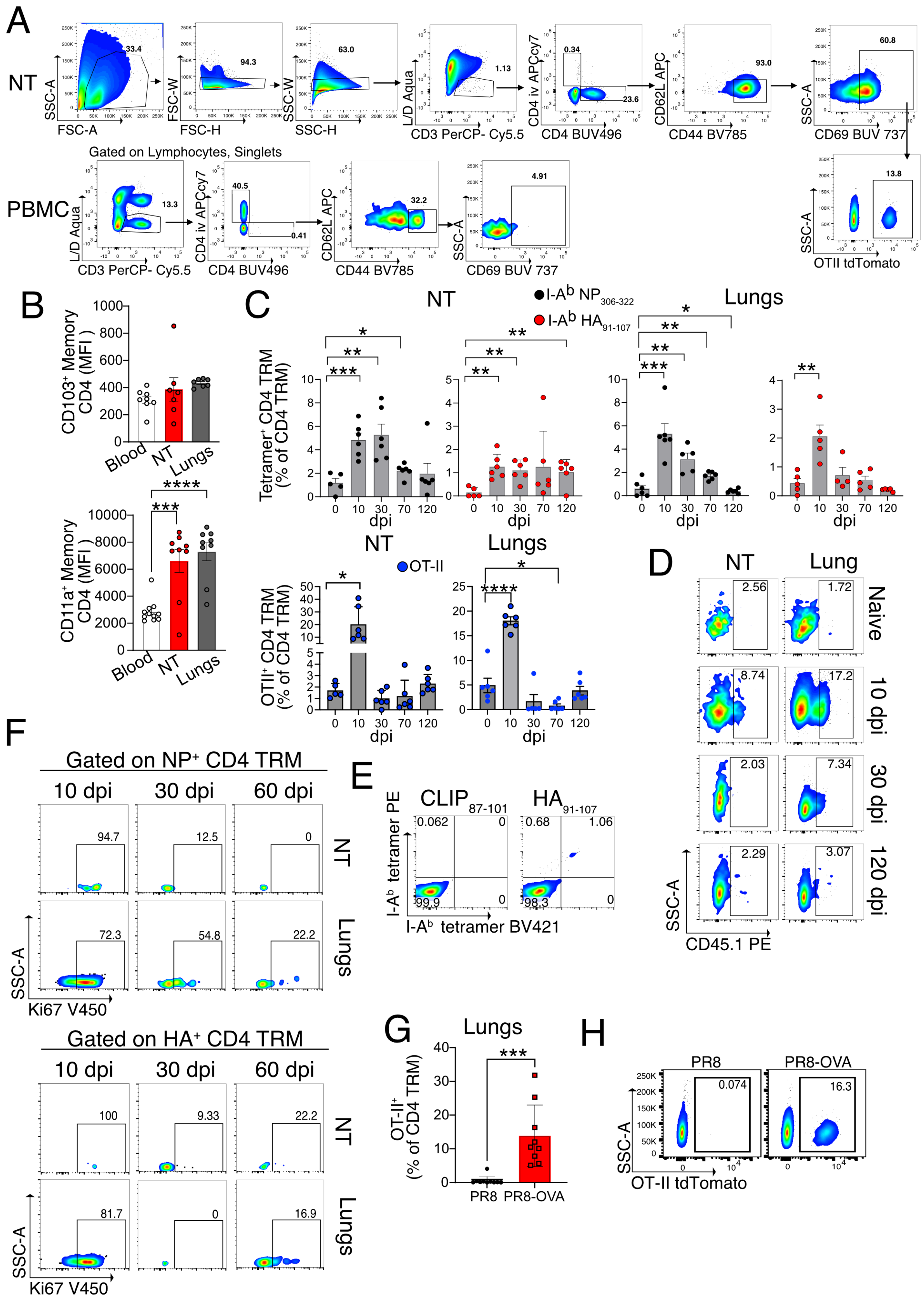

**Figure S1: Characterization of CD4 TRM in lungs and NT after IAV infection**

A) Gating strategy to identify OT-II$^+$ CD4 TRM in the NT.

B) Expression of CD103 and CD11a (median fluorescence intensity) on OT-II CD4 TRM in NT and lungs, and on total CD4 TEM of blood of CD45.2$^+$ recipients on day 30 following IAV infection. The experiment was repeated thrice and the results (mean ± s.e.m.) were pooled. NS, not significant, *****P*<0.0001; ****P*<0.001; ***P*<0.01; **P*<0.05 by one-way ANOVA, with Tukey's multiple comparison test

C) Scatter plot indicating the percentage of I-A$^b$ $NP_{306-322}$ tetramer$^+$, I-A$^b$ $HA_{91-107}$ tetramer$^+$ and OT-II CD4 TRM in lung (right panel) and NT (left panel) on different days post infection with PR8 Ova IAV intranasally. Each data point indicates an individual mouse. The experiment was performed once and the results (mean ± s.e.m.) were pooled. NS, not significant, *****P*<0.0001; ****P*<0.001; ***P*<0.01; **P*<0.05 by unpaired two-tailed t-test.

D) Flow cytometry plot showing the percentage of OT-II$^+$ CD4 TRM in the NT and lungs from different days post PR8 infection.

E) Flow cytometry plot showing the percentage of I-A$^b$ $HA_{91-107}$ PE tetramer$^+$ I-A$^b$ $HA_{91-107}$ BV421tetramer$^+$ CD4 TRM and I-A$^b$ human $CLIP_{87-101}$ PE tetramer$^+$ I-A$^b$ human $CLIP_{87-101}$ BV421 tetramer$^+$ CD4 TRM in the NT of mice on day 60 post PR8 infection.

F) Representative flow cytometry plots of Ki67$^+$ cells among I-A$^b$ $HA_{91-107}$ tetramer$^+$ and I-A$^b$ $NP_{306-322}$ tetramer+ CD4 TRM of lungs and NT isolated on day 10, 30 and 60 post PR8 infection.

G-H) Frequency of OT-II CD4 TRM in the lung on day 22 following infection with PR8 or PR8-Ova. F) Scatter plot showing the frequency of OT-II CD4 TRM. The experiment was performed twice and the results (mean ± s.e.m.) are pooled. NS, not significant, *****P*<0.0001; ****P*<0.001; ***P*<0.01; **P*<0.05 by unpaired two-tailed t-test. G) A representative flow cytometry plot indicating the frequency of OT-II CD4 TRM.

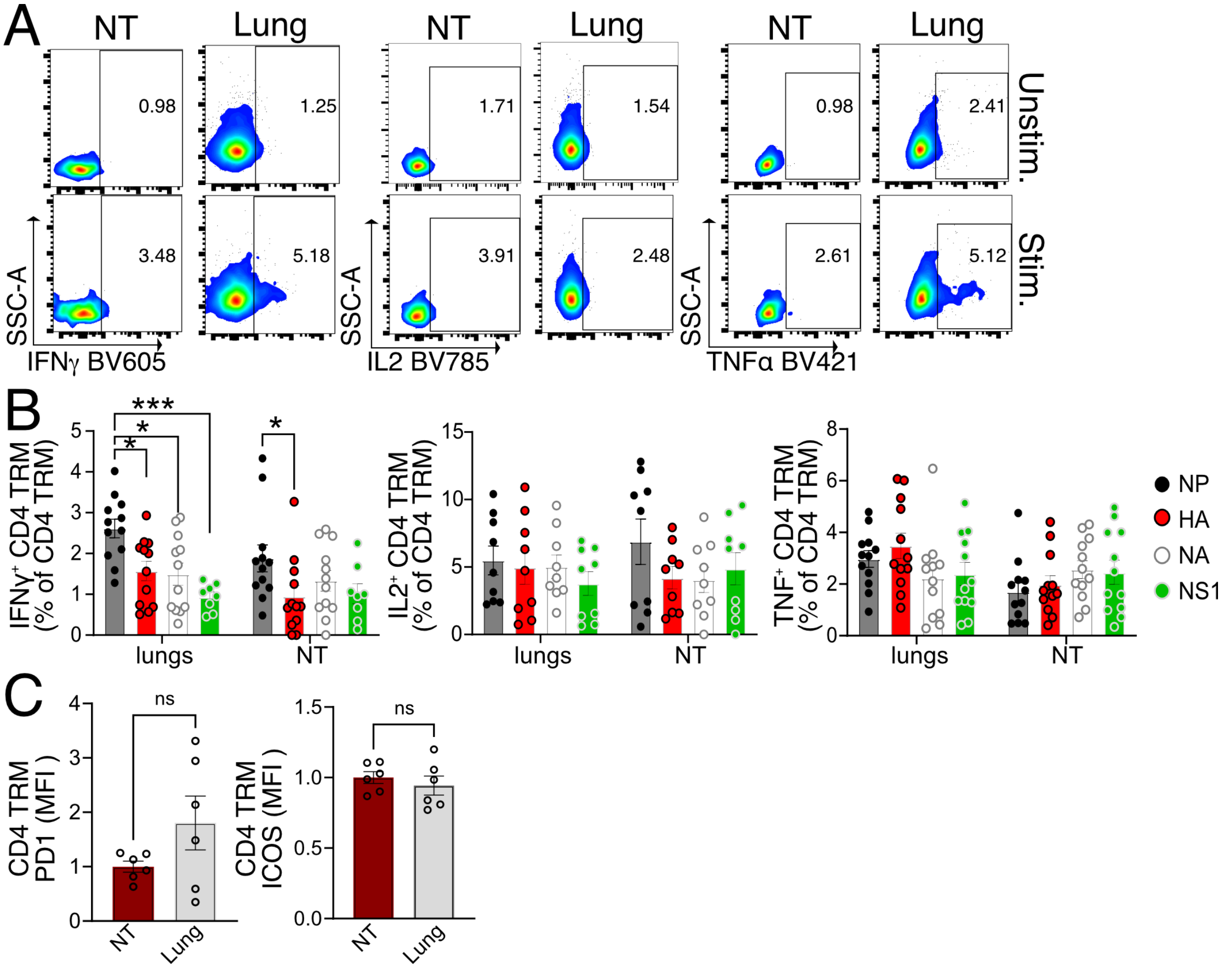


**Figure S2: IAV specific CD4 TRM of the NT are functional and exhibit immunodominance hierarchy similar to the lungs**

A) Flow cytometry plots indicating the percentage of CD4 TRM expressing IFN-γ, IL-2 and TNF with and without stimulation with IAV NP peptide pool. The organs are isolated on day 30 following PR8 IAV infection of mice.

B) Scatter plot indicating the percentage of CD4 TRM expressing IFN-γ, IL-2 and TNF after stimulation with IAV NP peptide pool, IAV HA peptide pool, IAV NA peptide pool and IAV NS1 peptide pool. The organs are isolated on day 30 following PR8 IAV infection of mice. The experiment was performed thrice and the results (mean ± s.e.m.) were pooled. NS, not significant, ****$P$<0.0001; ***$P$<0.001; **$P$<0.01; *$P$<0.05 by by two-way ANOVA, with Tukey's multiple comparison test.

C) Scatter plot showing the expression of PD-1 and ICOS (MFI normalized to average MFI of respective markers on NT CD4 TRM) on CD4 TRM of lungs and NT on day 30 following infection with PR8 IAV. The experiment was repeated twice and the results (mean ± s.e.m.) are pooled. NS, not significant, ****$P$<0.0001; ***$P$<0.001; **$P$<0.01; *$P$<0.05 by unpaired two-tailed t-test.

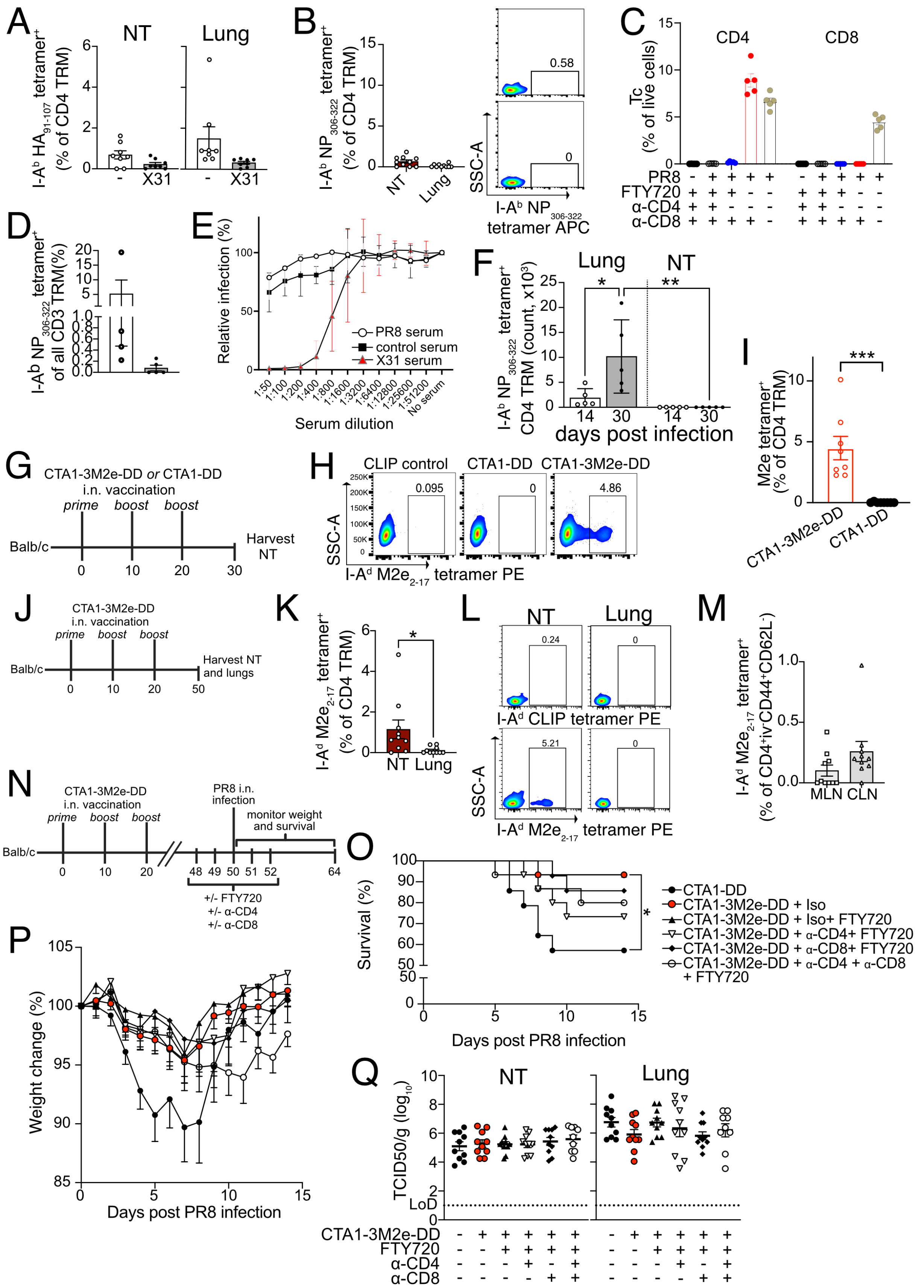

A
NT
Lung
I-A^b HA91-107 tetramer+ (% of CD4 TRM)
-
X31
B
I-A^b NP306-322 tetramer+ (% of CD4 TRM)
NT
Lung
0.58
0
SSC-A
I-A^b NP306-322 tetramer APC
C
CD4
CD8
Tc (% of live cells)
PR8
FTY720
α-CD4
α-CD8
D
I-A^b NP306-322 tetramer+ of all CD3 TRM(%)
E
Relative infection (%)
PR8 serum
control serum
X31 serum
1:50
1:100
1:200
1:400
1:800
1:1600
1:3200
1:6400
1:12800
1:25600
1:51200
No serum
Serum dilution
F
Lung
NT
I-A^b NP306-322 tetramer+ CD4 TRM (count, x10^3)
*
**
14
30
days post infection
G
CTA1-3M2e-DD or CTA1-DD
i.n. vaccination
prime
boost
boost
Balb/c
Harvest NT
H
CLIP control
CTA1-DD
CTA1-3M2e-DD
0.095
0
4.86
SSC-A
I-A^d M2e2-17 tetramer PE
I
M2e tetramer+ (% of CD4 TRM)
***
CTA1-3M2e-DD
CTA1-DD
J
CTA1-3M2e-DD
i.n. vaccination
prime
boost
boost
Balb/c
Harvest NT and lungs
K
I-A^d M2e2-17 tetramer+ (% of CD4 TRM)
*
NT
Lung
L
NT
Lung
0.24
0
I-A^d CLIP tetramer PE
5.21
0
SSC-A
I-A^d M2e2-17 tetramer PE
M
I-A^d M2e2-17 tetramer+ (% of CD4+iv-CD44+CD62L-)
MLN
CLN
N
CTA1-3M2e-DD
i.n. vaccination
prime
boost
boost
PR8 i.n.
infection
monitor weight
and survival
Balb/c
+/- FTY720
+/- α-CD4
+/- α-CD8
O
Survival (%)
CTA1-DD
CTA1-3M2e-DD + Iso
CTA1-3M2e-DD + Iso+ FTY720
CTA1-3M2e-DD + α-CD4+ FTY720
CTA1-3M2e-DD + α-CD8+ FTY720
CTA1-3M2e-DD + α-CD4 + α-CD8
+ FTY720
*
Days post PR8 infection
P
Weight change (%)
Days post PR8 infection
Q
NT
Lung
TCID50/g (log10)
LoD
CTA1-3M2e-DD
FTY720
α-CD4
α-CD8

**Figure S3: IAV specific NT CD4 TRM are induced by vaccination and provide protection.**

A) I-$A^b$ $HA_{91-107}$ tetramer- specific CD4 TRM in the NT and lungs of mice that were infected with X31 intranasally or left uninfected on day 30 following PR8 infection. The mice were treated with FTY720 and the organs were analyzed 6 days post X31 infection. Scatter plot indicating the percentage of I-$A^b$ $HA_{91-107}$ tetramer$^+$ cells among CD4 TRM in lungs and NT. The experiment was done twice and the results (mean ± s.e.m.) were pooled. NS, not significant, ****$P$<0.0001; ***$P$<0.001; **$P$<0.01; *$P$<0.05 by unpaired two-tailed t-test.

B) I-Ab NP306-322 tetramer-specific CD4 TRM in the NT and lungs of mice 6 days post X31 infection. The mice were treated with FTY720 from day -2 to day 5 post infection. Left: scatter plot indicating the percentage of I-Ab NP306-322 tetramer+ cells among CD4 TRM of lungs and NT. The experiment was done twice and the results (mean ± s.e.m.) were pooled. Right: representative flow cytometry plots indicating the percentages of I-Ab NP306-322 tetramer- specific CD4 TRM are shown.

C) Percentage of CD4$^+$ and CD8$^+$ T cells in the blood of mice on day 3 following secondary X31 infection from different groups as indicated in the fig. 3D.

D) Scatter plot showing the frequency of I-$A^b$ $NP_{306-322}$ tetramer$^+$ cells among CD3$^+$ T cells in the NT of mice treated with isotype control antibody or anti-CD4 and anti-CD8 antibody. The experiment was performed once. The result is shown as (mean ± s.e.m.). NS, not significant, ****$P$<0.0001; ***$P$<0.001; **$P$<0.01; *$P$<0.05 by unpaired two-tailed t-test.

E) Percentage of relative infection of MDCK cells by X31 IAV after incubation with serum derived on day 30 post infection of mice with PR8 or X31. The serum from mice immunized with irrelevant protein (Covid Spike or streptavidin) was used as controls. The experiment was performed twice and the results (mean ± s.e.m.) are pooled.

F) Graph indicating the absolute number of I-$A^b$ $NP_{306-322}{}^+$ CD4 TRM in lungs and NT of mice on day 14 and day 30 post infection with PR8 intratracheally. The experiment was done once. The experiment was performed once. The result is shown as (mean ± s.e.m.) NS, not significant, ****$P$<0.0001; ***$P$<0.001; **$P$<0.01; *$P$<0.05 by two-way ANOVA, with Tukey's multiple comparison test.

G-I) I-$A^d$ $M2e_{2-17}$ tetramer$^+$ CD4 TRM in the NT of BALB/c on day 10 after 3rd immunization with CTA1-DD or CTA1-3M2e-DD intranasally. G) Schematic representation of the experimental setup. H) Representative flow cytometry plots indicating the percentage of I-$A^d$ $M2e_{2-17}$ tetramer$^+$ CD4 TRM from different treatment groups. Staining with I-$A^d$ Human $CLIP_{87-101}$ tetramer (CLIP control) is used as the negative control. I) Scatter plot indicating the percentage of I-$A^d$ $M2e_{2-17}$ tetramer$^+$ cells among all CD4 TRM. The experiment was repeated twice and the results (mean ± s.e.m.) were pooled. NS, not significant, ****$P$<0.0001; ***$P$<0.001; **$P$<0.01; *$P$<0.05 by unpaired two-tailed t-test.

J-M) I-$A^d$ $M2e_{2-17}$ tetramer$^+$ CD4 TRM in the NT and lungs of BALB/c on day 30 post last immunization with CTA1-3M2e-DD intranasally. J) Schematic representation of the experimental setup. K) Scatter plot indicating the percentage of I-$A^d$ $M2e_{2-17}$ tetramer$^+$ cells among all CD4 TRM of lungs and NT. The experiment was repeated twice and the results (mean ± s.e.m.) were pooled. NS, not significant, ****$P$<0.0001; ***$P$<0.001; **$P$<0.01; *$P$<0.05 by unpaired two-tailed t-test. L) Representative flow cytometry plots indicating the percentage of I-$A^d$ $M2e_{2-17}$ tetramer$^+$ CD4 TRM and I-$A^d$ human

$CLIP_{87\text{-}101}$ tetramer$^+$ CD4 TRM from lungs and NT. M) Scatter plot indicating the percentage of I-A$^d$ $M2e_{2\text{-}17}$ tetramer$^+$ cells among all CD4 TEM of MLN and CLN in mice immunized as shown in fig S3J. The experiment was repeated twice and the results (mean ± s.e.m.) were pooled. NS, not significant, ****$P$<0.0001; ***$P$<0.001; **$P$<0.01; *$P$<0.05 by unpaired two-tailed t-test.

N-P) Survival rate and weight loss of CTA1-3M2e-DD or CTA1-DD immunized BALB/c mice that were infected with PR8 in URT restricted manner and were treated with or without FTY720 and anti-CD4 antibody or anti-CD8 antibody or isotype control antibody. N) Schematic representation of the experimental set up. O) Kaplan-Meier survival curves of mice from different groups after PR8 infection. The experiment was repeated thrice and the results were pooled. NS, not significant, ***$P$<0.0001; ***$P$<0.001; **$P$<0.01; *$P$<0.05 by log rank Mantel-Cox test. P) Weight loss curve for mice from different treatment groups as indicated. The experiment was repeated thrice and the results (mean ± s.e.m.) were pooled. Statistical comparisons between treatment groups (NS, not significant, ****$P$<0.0001; ***$P$<0.001; **$P$<0.01; *$P$<0.05) were performed using a linear mixed-effects model with comparisons of weight across treatment groups (averaged over time) using estimated marginal means (EMMs) and pairwise comparisons with Tukey's Honest Significant Difference (HSD) adjustment. CTA1-DD vs CTA1-3M2e-DD+Iso=*, CTA1-DD vs CTA1-3M2e-DD+Iso+FTY720=** and CTA1-DD vs CTA1-3M2e-DD + FTY720 + a-CD8=*. Statistical comparisons across groups for each day are reported in Table S2.

Q) Scatter plot indicating viral titers ($TCID_{50}$/g) in NT and lungs of immunized mice as shown in fig S3N on day 3 following infection with PR8 IAV. The experiment was repeated thrice and the results (mean ± s.e.m.) are pooled. NS, not significant, ****$P$<0.0001; ***$P$<0.001; **$P$<0.01; *$P$<0.05 by unpaired two-tailed t-test.

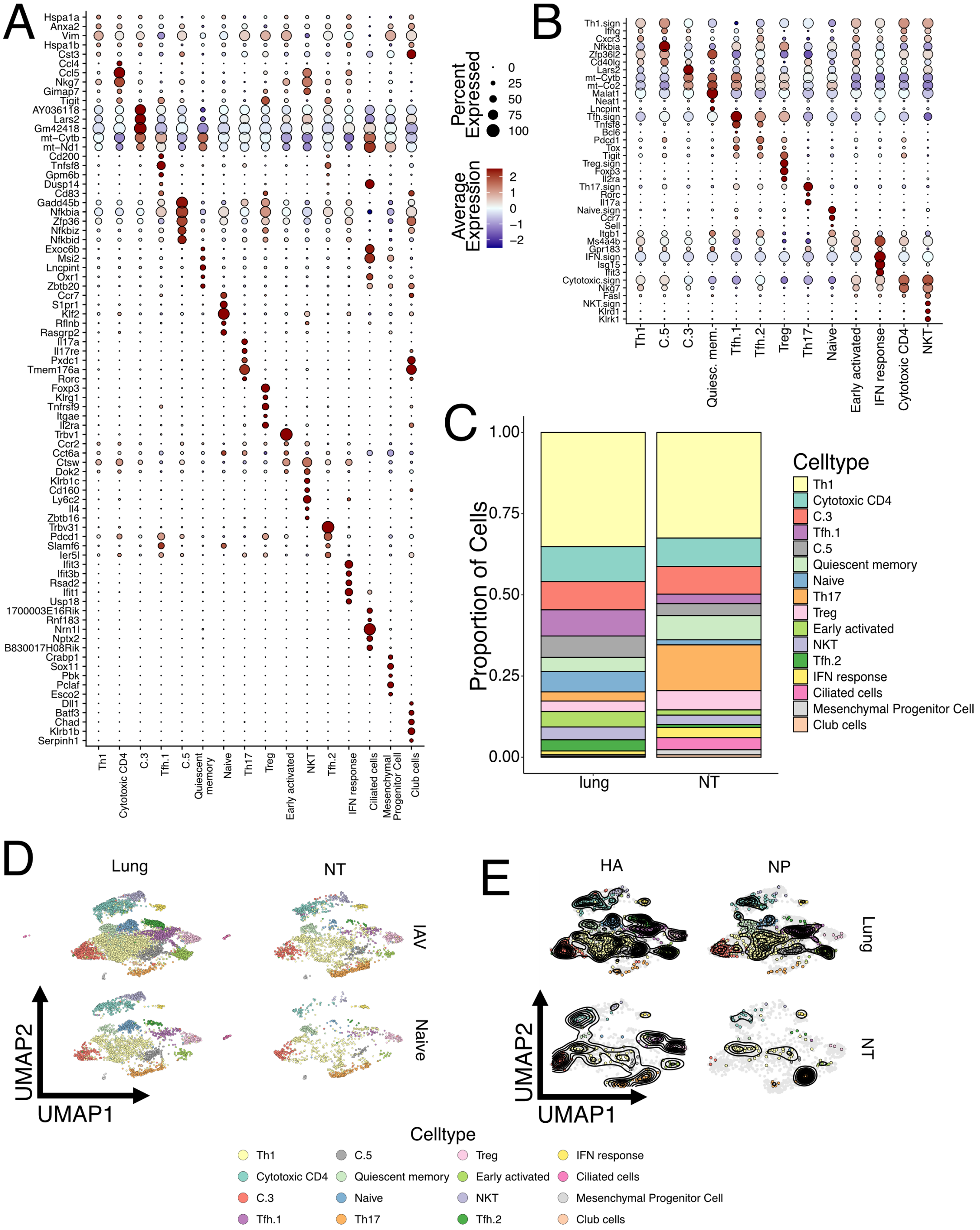
A
Hspa1a
Anxa2
Vim
Hspa1b
Cst3
Ccl4
Ccl5
Nkg7
Gimap7
Tigit
AY036118
Lars2
Gm42418
mt−Cytb
mt−Nd1
Cd200
Tnfsf8
Gpm6b
Dusp14
Cd83
Gadd45b
Nfkbia
Zfp36
Nfkbiz
Nfkbid
Exoc6b
Msi2
Lncpint
Oxr1
Zbtb20
Ccr7
S1pr1
Klf2
Rflnb
Rasgrp2
Il17a
Il17re
Pxdc1
Tmem176a
Rorc
Foxp3
Klrg1
Tnfrsf9
Itgae
Il2ra
Trbv1
Ccr2
Cct6a
Ctsw
Dok2
Klrb1c
Cd160
Ly6c2
Il4
Zbtb16
Trbv31
Pdcd1
Slamf6
Ier5l
Ifit3
Ifit3b
Rsad2
Ifit1
Usp18
1700003E16Rik
Rnf183
Nrn1l
Nptx2
B830017H08Rik
Crabp1
Sox11
Pbk
Pclaf
Esco2
Dll1
Batf3
Chad
Klrb1b
Serpinh1
Th1
Cytotoxic CD4
C.3
Tfh.1
C.5
Quiescent memory
Naive
Th17
Treg
Early activated
NKT
Tfh.2
IFN response
Ciliated cells
Mesenchymal Progenitor Cell
Club cells
Percent Expressed
0
25
50
75
100
Average Expression
2
1
0
−1
−2
B
Th1.sign
Ifng
Cxcr3
Nfkbia
Zfp36l2
Cd40lg
Lars2
mt−Cytb
mt−Co2
Malat1
Neat1
Lncpint
Tfh.sign
Tnfsf8
Bcl6
Pdcd1
Tox
Tigit
Treg.sign
Foxp3
Il2ra
Th17.sign
Rorc
Il17a
Naive.sign
Ccr7
Sell
Itgb1
Ms4a4b
Gpr183
IFN.sign
Isg15
Ifit3
Cytotoxic.sign
Nkg7
Fasl
NKT.sign
Klrd1
Klrk1
Th1
C.5
C.3
Quiesc. mem.
Tfh.1
Tfh.2
Treg
Th17
Naive
Early activated
IFN response
Cytotoxic CD4
NKT
C
Proportion of Cells
1.00
0.75
0.50
0.25
0.00
lung
NT
Celltype
Th1
Cytotoxic CD4
C.3
Tfh.1
C.5
Quiescent memory
Naive
Th17
Treg
Early activated
NKT
Tfh.2
IFN response
Ciliated cells
Mesenchymal Progenitor Cell
Club cells
D
Lung
NT
IAV
Naive
UMAP2
UMAP1
E
HA
NP
Lung
NT
UMAP2
UMAP1
Celltype
Th1
Cytotoxic CD4
C.3
Tfh.1
C.5
Quiescent memory
Naive
Th17
Treg
Early activated
NKT
Tfh.2
IFN response
Ciliated cells
Mesenchymal Progenitor Cell
Club cells

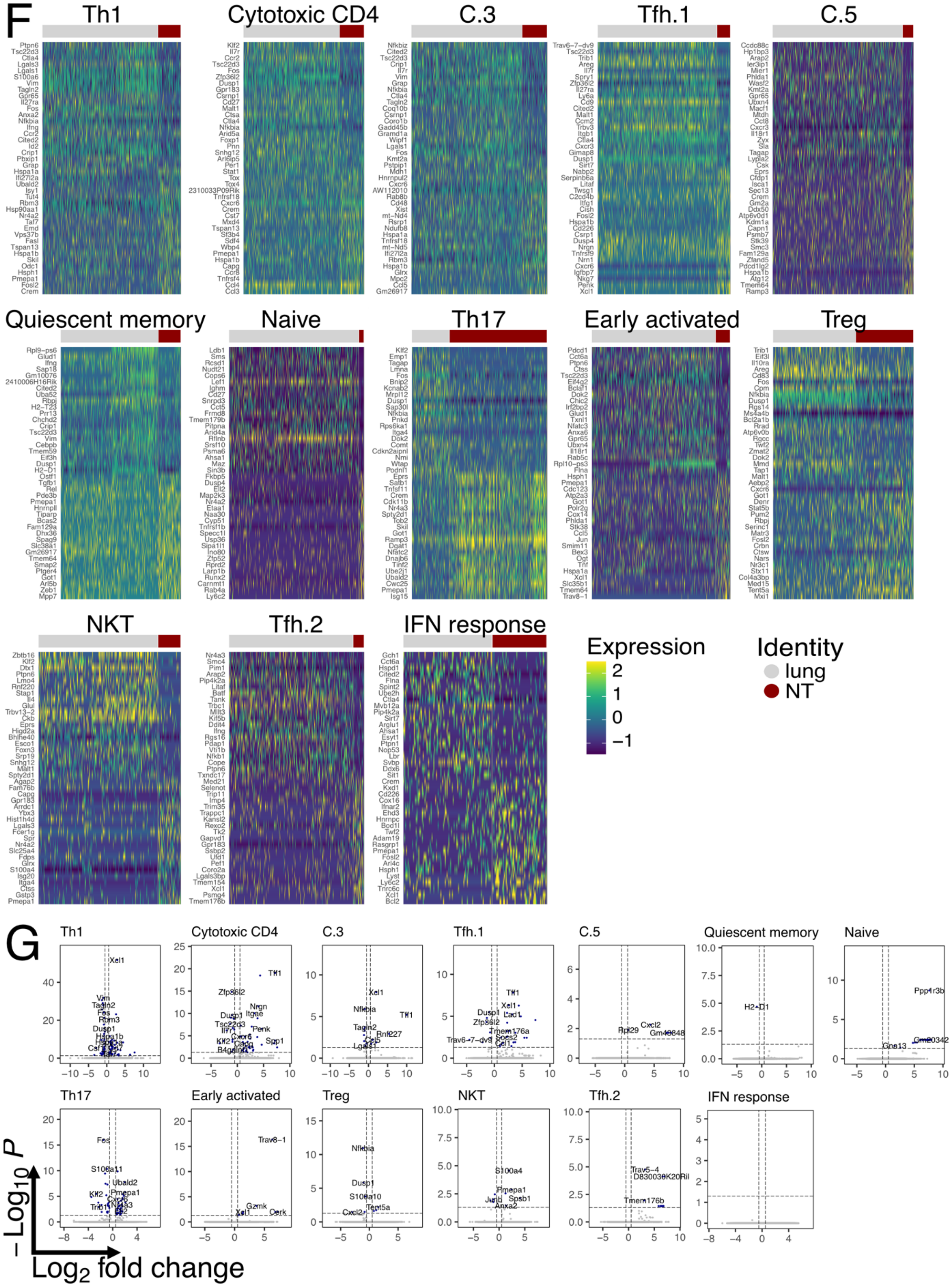

F
Th1
Cytotoxic CD4
C.3
Tfh.1
C.5
Quiescent memory
Naive
Th17
Early activated
Treg
NKT
Tfh.2
IFN response
Expression
2
1
0
−1
Identity
lung
NT
G
Th1
Cytotoxic CD4
C.3
Tfh.1
C.5
Quiescent memory
Naive
Th17
Early activated
Treg
NKT
Tfh.2
IFN response
−Log10 P
Log2 fold change

**Figure S4: scRNA-Seq of CD4 TRM in the NT vs lungs**

A) Dot plot representing mean expression of top 5 marker genes for each T cell cluster, identified by the FindAllMarker function and ordered according log2 fold change. Color intensity from blue to red indicates average expression of genes and size of the dot depicts percentage of cells expressing the gene within the clusters.

B) Dot plot representing mean expression of selected marker genes for each T cell cluster. Color intensity from blue to red indicates average expression of genes and size of the dot depicts percentage of cells expressing the gene within the clusters. Only T cell clusters were included in the analysis.

C) Bar graph showing proportion of each UMAP cluster divided by organ. All cells from all mice were included in the analysis.

D) UMAP plot split by tissue (lung or NT) and infection status (IAV infected or naïve) and colored according to identified clusters.

E) UMAP plot split by tissue (lung or NT) and antigen specificity (I-$A^b$ $NP_{306-322}$ tetramer or I-$A^b$ $HA_{91-107}$ tetramer) and colored according to identified clusters. Grey dots indicate cells which were not specific for the selected antigen within the specific organ. Contour lines indicate the density of the cells.

F) Heat maps showing the top 20 differentially expressed genes between NT (NT; dark red) and lungs (grey) in different cell clusters. Color scale indicates expression intensity.

G) Volcano plot showing differentially expressed genes upregulated in the NT *vs* lungs among different cell clusters. The dotted lines indicate *p* value and Log2 fold change cutoffs.

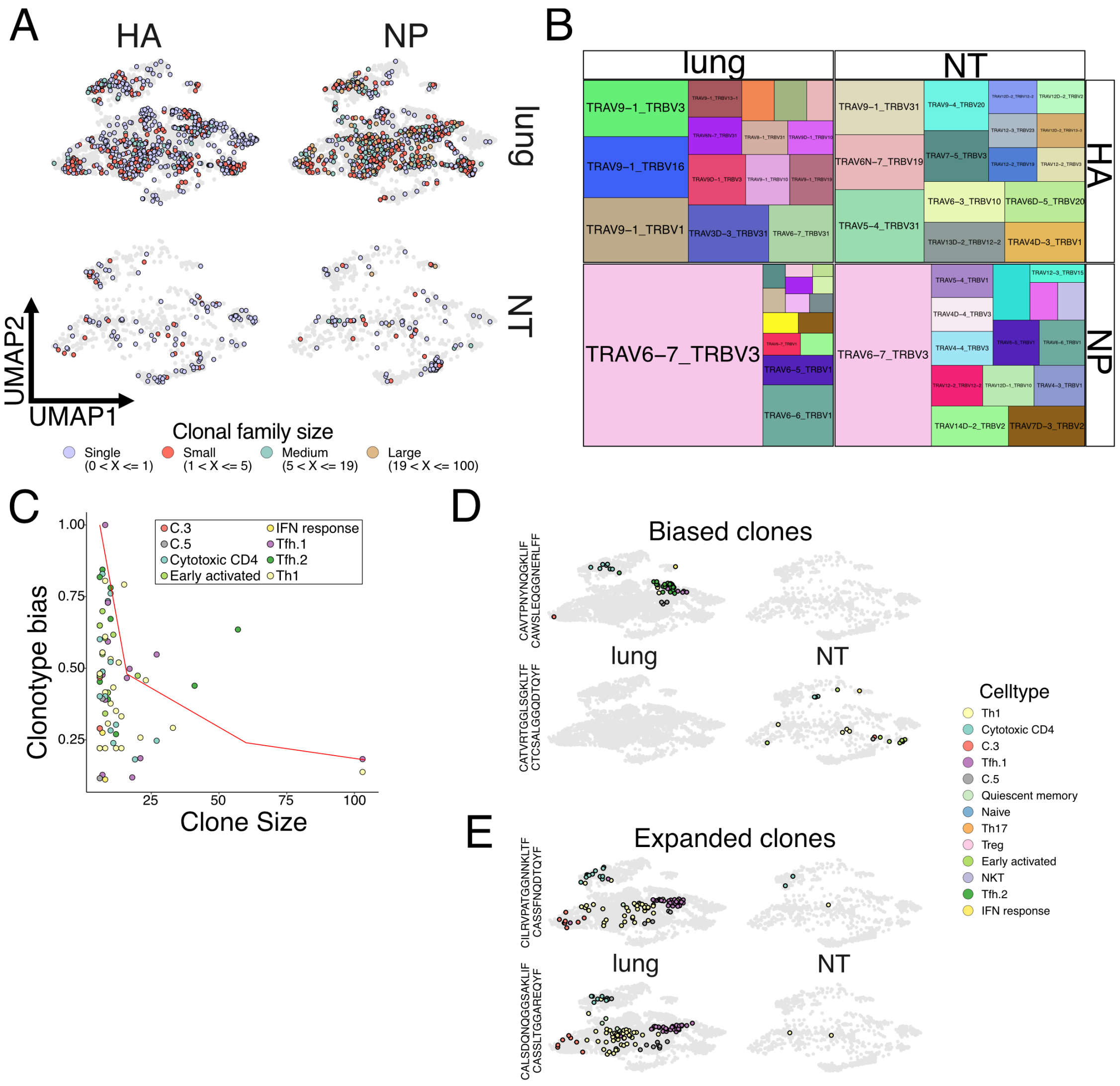


**Figure S5: Clonality of antigen specific CD4 TRM from lungs and NT**

A) UMAP plot split by tissue (lung or NT) and antigen specificity (I-A$^b$ NP$_{306-322}$ tetramer or I-A$^b$ HA$_{91-107}$ tetramer) and colored according to clone size for CD4 TRM in IAV infected mice. Clones were binned into single clones (1 member; light blue), small (2 to5 members; red), medium (6 to 19 members; green), large (over 20 members; gold). Light grey dots represent cells where TCR clonotype could not be assigned.

B) Tile plot indicating paired TRAV and TRBV genes usage split by tissue (lung or NT) and antigen specificity (I-A$^b$ NP$_{306-322}$ tetramer or I-A$^b$ HA$_{91-107}$ tetramer) in CD4 TRM of IAV infected mice. The size of the tile is proportional to TRAV usage frequency.

C) Clonotype bias plot for clones with at least 6 cells. Clonotypes are colored according to predominant cluster. The red line highlights the null distribution (background distribution) for each clone size. The clones to the right of the red line are biased clones.

D) UMAP plot, split by tissue (lung or NT), depicting two biased clones. Cells belonging to the clonal family are colored according to the cluster while other cells are in light grey.

E) UMAP plot, split by tissue (lung or NT), depicting the two most expanded clones. Cells belonging to the clonal family are colored according to the cluster while other cells are in light grey.

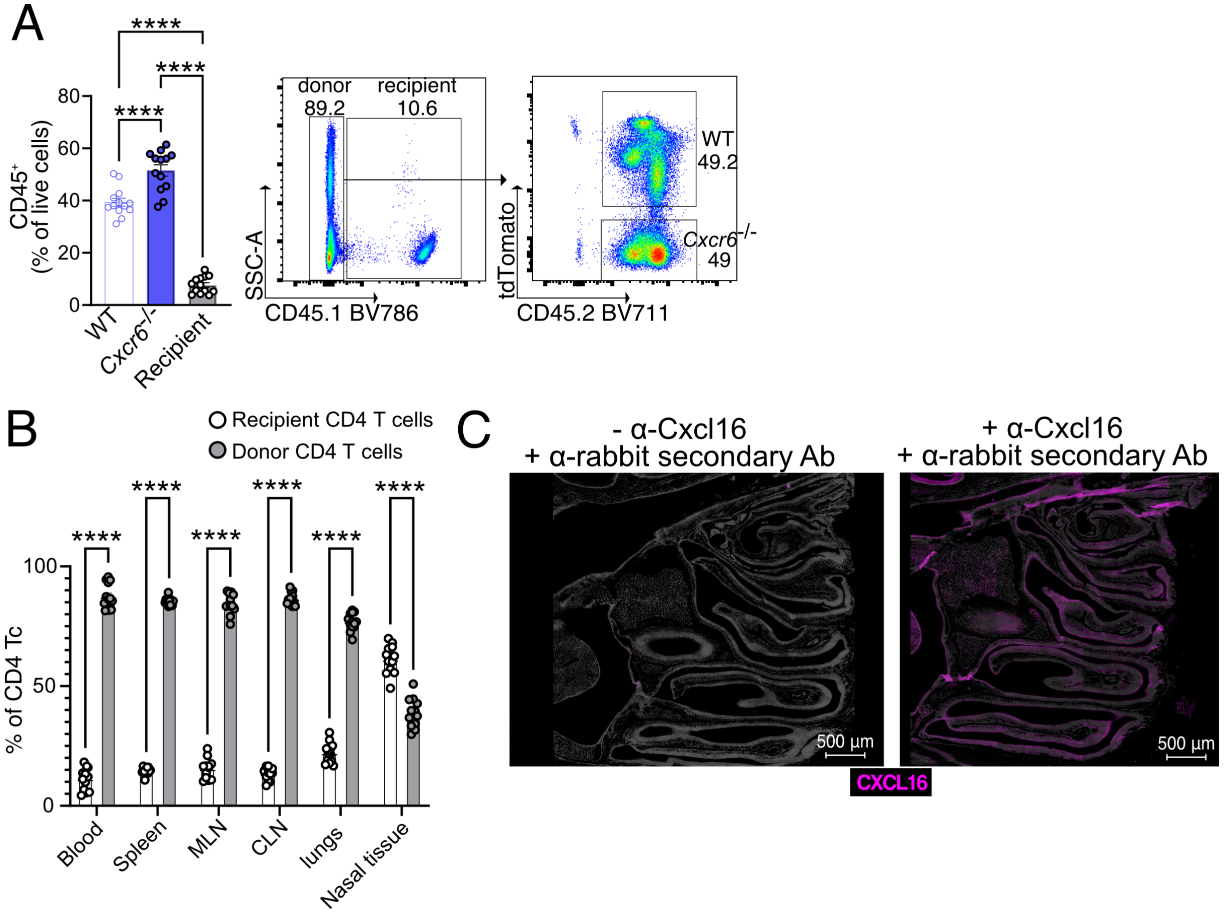


**Figure S6: CXCR6-CXCL16 axis promotes NT CD4 TRM establishment**

A) Percentage of $CD45^{+}$ donor cells (both *Cxcr6*$^{-/-}$ and WT) cells and recipient cells in the blood of BM chimera on day 67 following BM transplantation. Left panel: Scatter plot for the percentage of donor cells and recipient cells. The experiment was repeated thrice and the results (mean ± s.e.m.) were pooled. NS, not significant, *****P*<0.0001; ****P*<0.001; ***P*<0.01; **P*<0.05 by one-way ANOVA, with Tukey's multiple comparison test. Right panel: A representative flow cytometry plot showing the percentage of donor cells and recipient cells.

B) Scatter plot showing the percentage of donor and recipient cells among CD4 T cells in different organs in the BM chimera on day 30 post infection with PR8. The experiment was repeated thrice and the results (mean ± s.e.m.) were pooled. NS, not significant, *****P*<0.0001; ****P*<0.001; ***P*<0.01; **P*<0.05 by two-way ANOVA, with Tukey's multiple comparison test.

C) Representative microscopic image of CXCL16 expression (magenta) in the olfactory epithelium of the murine NT is shown on the right panel. A negative control for CXCL16 staining showing NT stained with α-rabbit secondary IgG Texas red only. NT is isolated on day 30 post PR8-Ova infection from mice that received OT-II CD4 T cells.

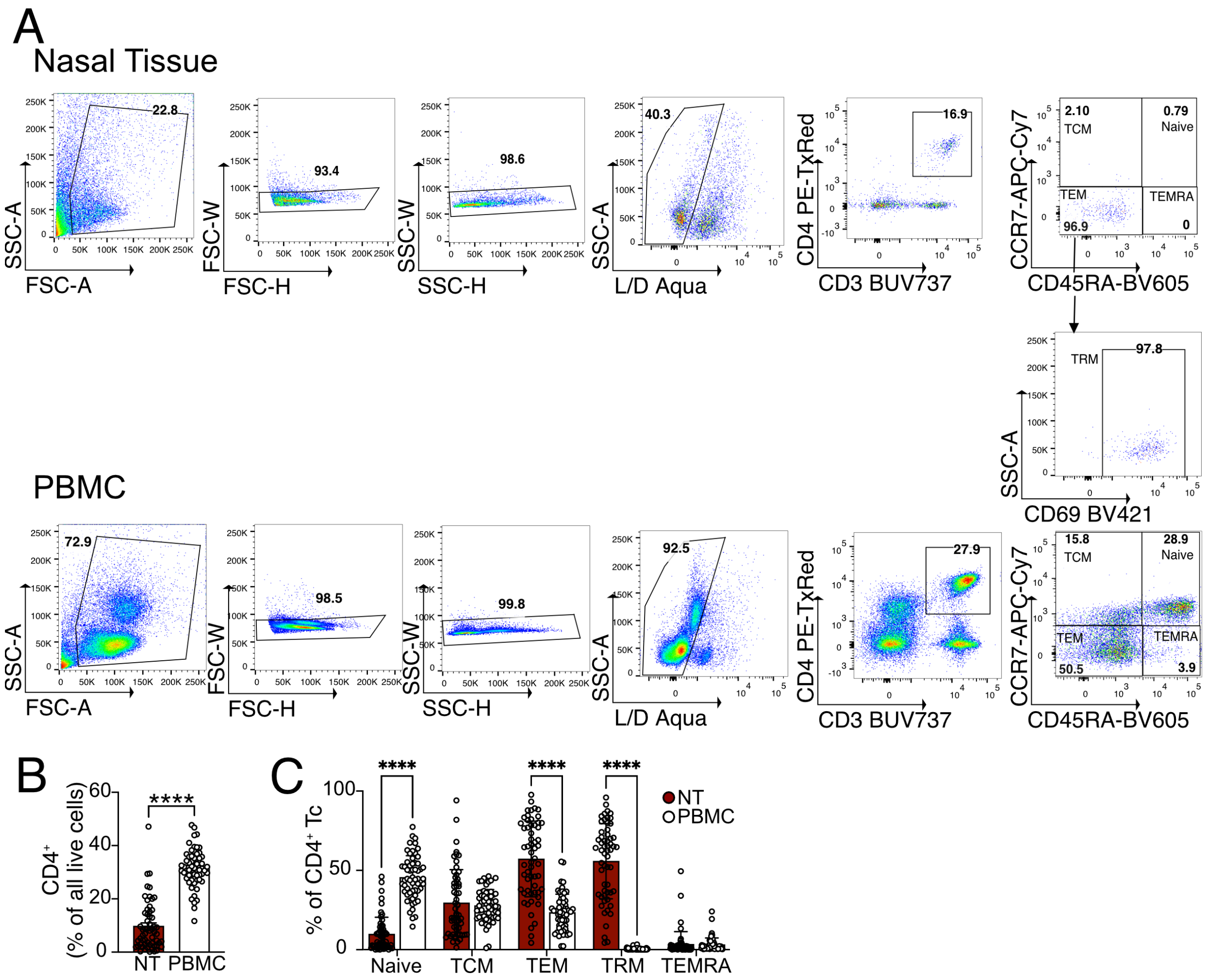


**Figure S7: IAV-responsive CD4 TRM in the nasal tissue and CD4 TEM in the PBMC of healthy controls**

A) Gating strategy to identify CD4 TEM ($CD4^{+}CCR7^{-}CD45RA^{-}$), CD4 TCM ($CD4^{+}CCR7^{+}CD45RA^{-}$), Naïve CD4 Tc ($CD4^{+}CCR7^{+}CD45RA^{+}$), CD4 TEMRA($CD4^{+}CCR7^{-}CD45RA^{+}$) and CD4 TRM ($CD4^{+}CCR7^{-}CD45RA^{-}CD69^{+}$) in NT and blood.

B) Scatter plot showing the frequency of CD4 Tc in the NT and PBMC. The experiment was performed seven times and the results (mean ± s.e.m.) are pooled. NS, not significant, ****$P$<0.0001; ***$P$<0.001; **$P$<0.01; *$P$<0.05 by unpaired two-tailed t-test.

C) Scatter plot showing the frequency of naïve CD4 Tc ($CD4^{+}CCR7^{+}CD45RA^{+}$), CD4 TCM ($CD4^{+}CCR7^{+}CD45RA^{-}$), CD4 TEM ($CD4^{+}CCR7^{-}CD45RA^{-}$), CD4 TEMRA($CD4^{+}CCR7^{-}CD45RA^{+}$) and CD4 TRM ($CD4^{+}CCR7^{-}CD45RA^{-}CD69^{+}$) in NT and blood. NS, not significant, ****$P$<0.0001; ***$P$<0.001; **$P$<0.01; *$P$<0.05 by two-way ANOVA, with Tukey's multiple comparison test.

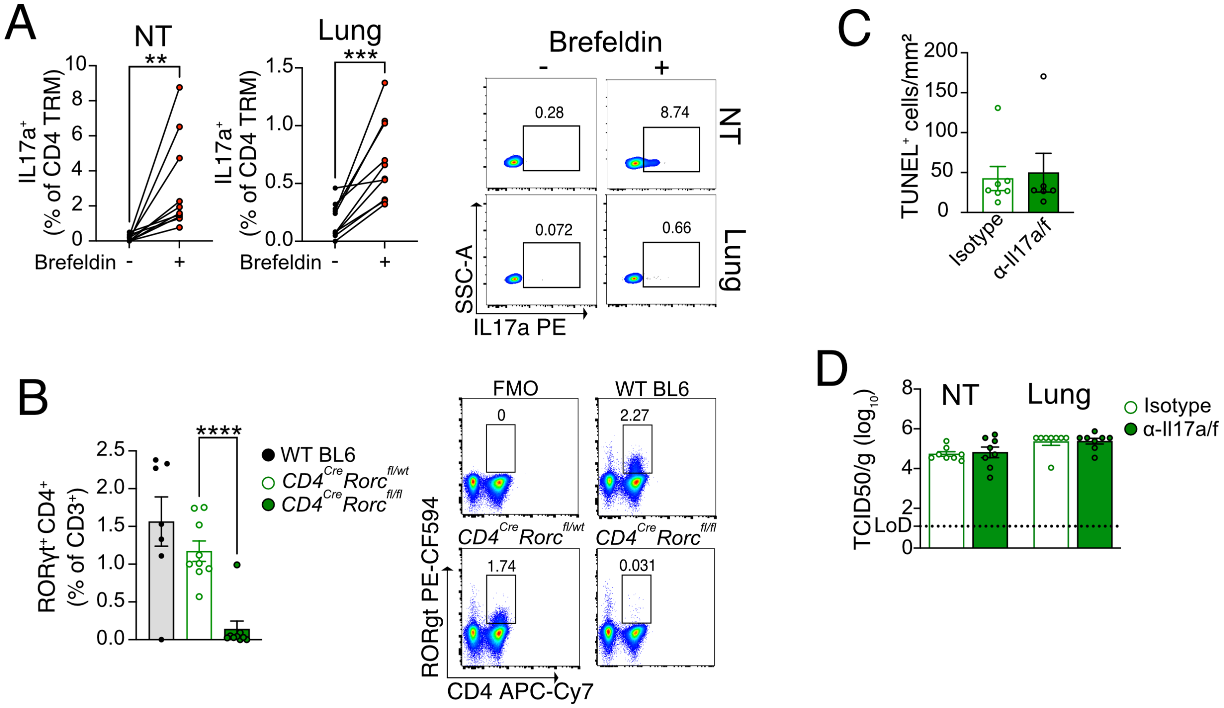


**Figure S8: IL-17 is dispensable for viral clearance in the NT**

A) Percentage of IL-17a+ cells among CD4 TRM of the lungs and NT that are stimulated with $NP_{306-322}$ peptide in the presence or absence of Brefeldin A. Left panel: Graph showing expression of IL-17a from the cells that are stimulated in the presence or absence of Brefeldin A connected by a line. Each line on the graph indicates the cells from individual mouse. The experiment was repeated twice and the results are pooled. NS, not significant, ****$P$<0.0001; ***$P$<0.001; **$P$<0.01; *$P$<0.05 by two-sided Wilcoxon matched-pairs signed-rank test. Right panel: Representative flow cytometry plots showing the expression of IL17-a among CD4 TRM of lungs and NT.

B) RORγt$^+$CD4$^+$ T cells in the Peyer's patches of WT C57BL/6 mice, *CD4$^{cre}$Rorc$^{fl/wt}$ and CD4$^{cre}$Rorc$^{fl/fl}$* mice. Left panel: Scatter plot showing the frequency of RORγt$^+$CD4$^+$ T cells among all CD3$^+$ T cells. The experiment was done twice and the results (mean ± s.e.m.) were pooled. NS, not significant, ****$P$<0.0001; ***$P$<0.001; **$P$<0.01; *$P$<0.05 by unpaired two-tailed t-test. Right panel: A representative flow cytometry plot for the percentage of RORγt$^+$CD4$^+$ T cells in different groups.

C-D) Microscopy for TUNEL$^+$ cells and viral titer ($TCID_{50}$/g) from organs of mice infected with PR8 and reinfected with X31 IAV on day 30 following PR8 IAV infection. C) Scatter plot showing number of TUNEL$^+$ cells in the nasal septum (respiratory region) on day 4 post X31 IAV infection. D) Viral titer ($TCID_{50}$/g) from NT and lungs on day 3 post X31 IAV infection.
